# Landscape-scale forest loss as a catalyst of population and biodiversity change

**DOI:** 10.1101/473645

**Authors:** Gergana N. Daskalova, Isla H. Myers-Smith, Anne D. Bjorkman, Shane A. Blowes, Sarah R. Supp, Anne Magurran, Maria Dornelas

**Affiliations:** School of GeoSciences, University of Edinburgh, West Mains Road, Edinburgh EH9 3FF, Scotland; Biological and Environmental Sciences, University of Gothenburg, 405 30 Gothenburg, Sweden; German Centre for Integrative Biodiversity Research (iDiv), Deutscher Pl. 5E, Leipzig 04103, Germany; Data Analytics Program, Denison University, 100 W College St, Granville, OH 43023, USA; Centre for Biological Diversity, University of St Andrews, Greenside Place, St Andrews KY16 9TF, Scotland

## Abstract

Global assessments have highlighted land-use change as a key driver of biodiversity change. However, we lack real-world global-scale estimates of how habitat transformations such as forest loss and gain are reshaping biodiversity over time. Here, we quantify the influence of 150 years of forest cover change on populations and ecological assemblages worldwide and across taxa by analyzing change in 6,667 time series. We found that forest loss simultaneously intensified ongoing increases and decreases in abundance, species richness and temporal species replacement (turnover) by up to 48%. Temporal lags in these responses extended up to 50 years and increased with species’ generation time. Our findings demonstrate that land-use change precipitates divergent population and biodiversity change, highlighting the complex biotic consequences of deforestation and afforestation.

**One Sentence Summary:** Declines in forest cover amplify both gains and losses in population abundance and biodiversity over time.

## Main Text

Accelerating human impacts are reshaping Earth’s ecosystems (*1*). The abundance of species’ populations (*2*) and the richness (*3*–*5*) and composition (*5*) of ecological assemblages at sites around the world are being altered over time in complex ways (*6*–*8*). However, we currently have only a limited quantitative understanding of how global change drivers, such as land-use change, produce heterogeneous local-scale patterns in population abundance and biodiversity over time (*7, 9, 10*), as highlighted by the recent IPBES Global Assessment (*1*). In terrestrial ecosystems, much of our current knowledge stems from space-for-time (*11, 12*) and modelling projection approaches (*13, 14*) that attribute population and richness declines to different types of land-use change, including reductions in forest cover. Yet, space-for-time methods can overestimate the effects of global change drivers compared to long-term monitoring, because they do not account for ecological lags (*7, 15, 16*) and community self-regulation (*17*). Ongoing controversy about the diverse impacts of habitat fragmentation on biodiversity (*18*–*20*), could be in part attributable to a lack of data from real-world contexts encompassing the full spectrum of forest fragmentation. As tree planting is increasingly viewed as a global climate change mitigation tool (*21*), we must understand the ecosystem and biodiversity consequences of tree cover loss and gain. Recent global-scale datasets of past land cover reconstructions (*22*) and high-resolution remote-sensing observations (*23, 24*) provide a novel opportunity to quantify decreases and increases in forested areas (hereafter, “forest loss and gain”), at landscape scales and at sites across the world. By integrating landscape-scale forest loss estimates with over five million population and biodiversity records (*25, 26*), our analysis provides unprecedented insight into the influence of land-use change on population and biodiversity change.

In our study, we ask how the timing and magnitude of landscape-scale forest loss affects time series of 4,228 species’ populations and 2,339 ecological assemblages (Fig. 1A). We assess variation in rates of population change (trends in numerical abundance, hereafter called “populations”) and biodiversity change (trends in species richness and community composition) across four vertebrate taxa, terrestrial invertebrates and plants (Figs. 1-2, S1-S2). First, we compared population and biodiversity change before and after the year of the largest reduction in forested area across the duration of each time series (hereafter, “contemporary peak forest loss”, Fig. 1B). Second, we asked how population and biodiversity change were related to the overall magnitude of forest loss experienced from 2000 to 2016 (the duration of the Forest Cover Change Dataset, Fig. 1B). Third, we investigated among-taxa variation in temporal lags (the time period between contemporary peak forest loss and maximum change in populations and assemblages, Fig. 1B). We provide the first temporal global assessment of the consequences of forest cover change for biodiversity for 2,157 geographic locations across the planet, allowing us to quantify the immediate and lagged impacts of forest loss and gain on population and biodiversity change.

**Fig 1.**
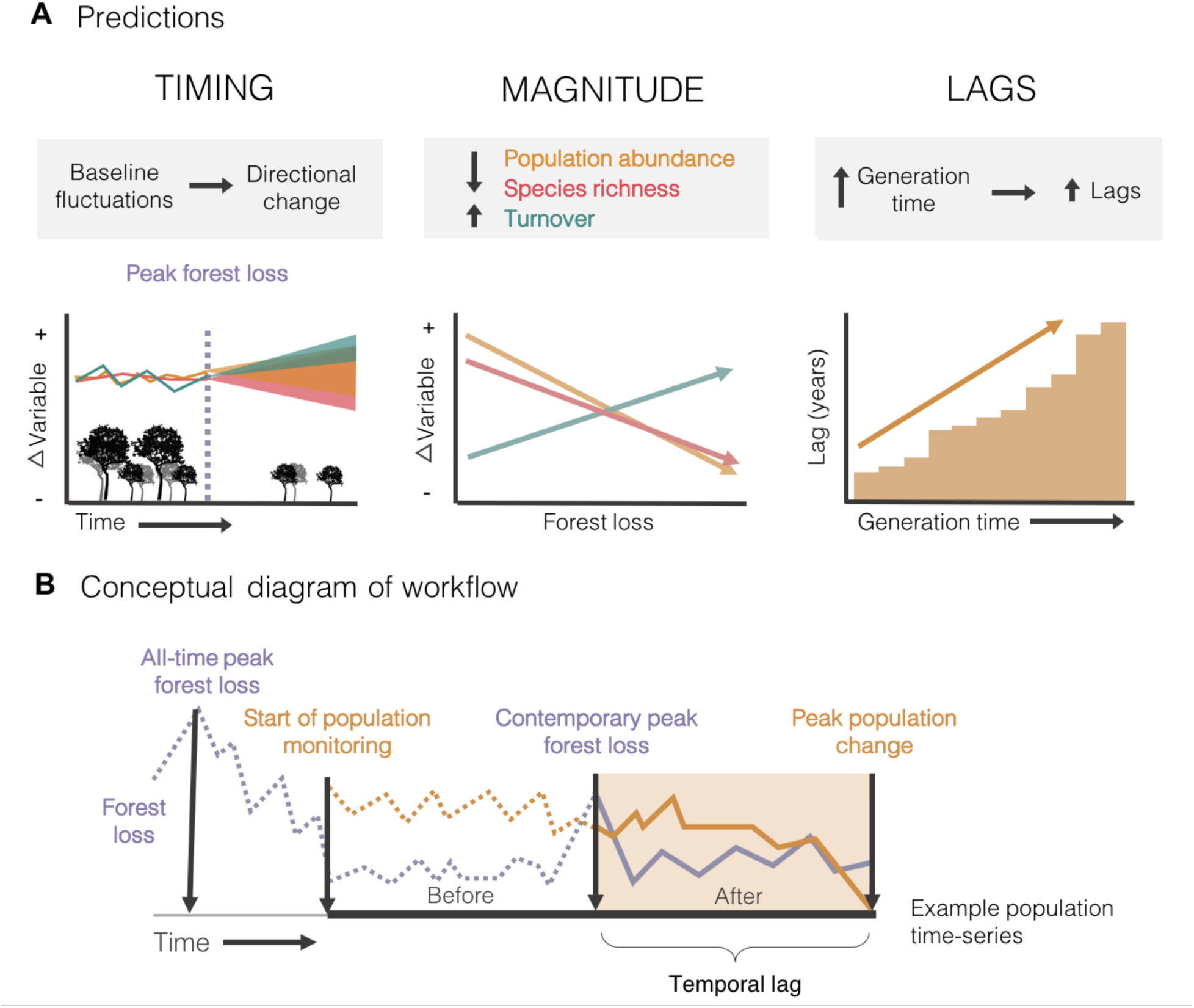
Our study tests three pathways through which forest loss can influence the population abundance of species and the richness and turnover of ecological assemblages: the timing, magnitude and temporally delayed effects of forest loss. **A**, Conceptual diagrams of our predictions outlined with respect to population change, richness change and turnover (temporal species replacement). **B**, Analytical workflow. See Figs. S1 and S2 for workflow of analyses and Materials and Methods sections one through three for further details.

**Fig. 2.**
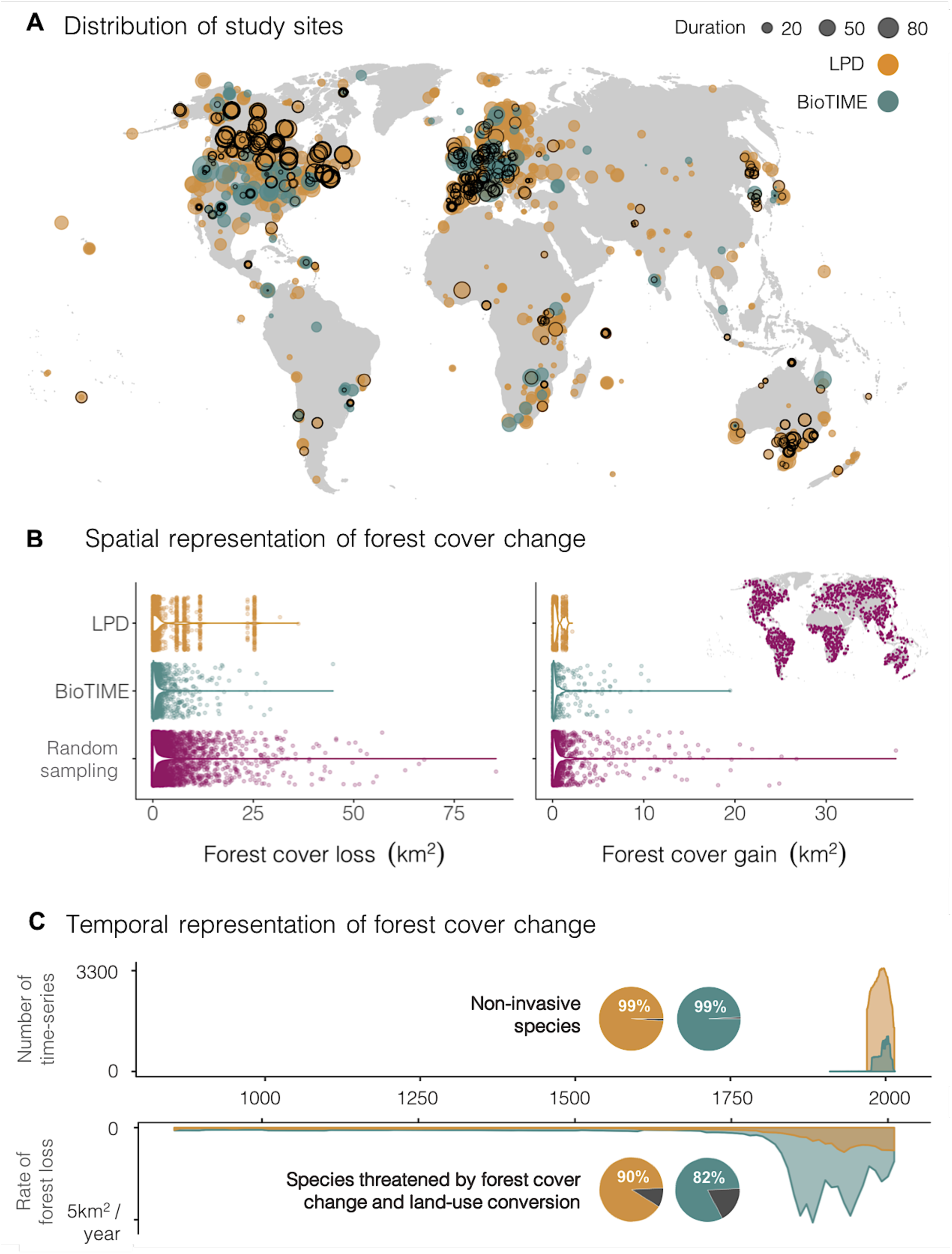
Population and biodiversity monitoring over time at sites around the world spans the global variation in forest cover change. **A**, Locations and duration of 542 Living Planet Database (LPD) and 199 BioTIME studies, containing 6,497 time series from 2,157 sites (black outline shows sites that were forested at the start of the monitoring (1,247 sites); see Table S2 for sample size in each woody biome). **B**, 44% of all time series experienced historic or contemporary forest loss of comparable magnitude to forest cover change across a random sample of geographical locations (shown on map inset in **B**) from the global distribution of forest cover loss and gain. We did not detect directional effects of forest gain across monitored sites. We present results concerning forest gain in Figs. S8-S9. **C**, presents the timeline of monitoring and forest loss across the sites represented in the LPD and BioTIME databases (for variation in monitoring periods among time series, see Figs. S6-S7). Insets in panel **C** show the proportion of study species that are not classified as invasive (top) and that are threatened by land-use change, based on species’ IUCN threat assessments (bottom, see Fig. S3E for details).

Land-use change, and specifically forest cover loss, alters habitat and resource availability (*11, 27, 28*) and is a global threat for the persistence of terrestrial species (*32*, Figs. 2, S3E). We predicted the greatest population and species richness declines after contemporary peak forest loss across time series. We also expected that greater population and richness declines will coincide with larger losses in forest cover. Additionally, we predicted greater temporal species replacement (hereafter, “turnover”) with greater forest loss. Finally, species with longer generation times typically respond more slowly to environmental change (*30*). We thus expected lags in ecological responses to forest loss to increase with longer generation times across taxa. Overall, our analyses provide real-world estimates of the magnitude of landscape-scale forest loss impacts and the pace at which they are altering terrestrial ecosystems around the planet.

To quantitatively test the ecological consequences of land-use change in the world’s woody biomes, we used two global databases. The Living Planet Database includes 133,092 records of the number of individuals from a species in a given area over time, hereafter called “populations” (*25*) and the BioTIME database comprises 4,970,128 records of the number and abundance of species in ecological assemblages over time (*26*). These are currently the two largest databases of population and community time series, respectively. Together, these temporal records represent a range of taxa including amphibians, birds, mammals, reptiles, invertebrates and plants (Fig. S3C-D) and 2,157 sites which cover almost the entire spectrum of forest loss and gain around the world (Fig. 2). We used a standardized cell size of 96 km^2^ to match response variables to landscape-scale forest change but note that analyses were robust to cell size (see Methods and Figure S16).

We calculated population change (*μ*) using state-space models that accounted for observation error and random fluctuations (*31*), and richness change (slopes of rate of change over time) using mixed effects models. We quantified temporal change in species composition as the turnover component of Jaccard’s dissimilarity measure (change due to species replacement, *31*). Turnover is often independent of changes in species richness (*8*) and is the dominant component of compositional change across the BioTIME time series (*33*). We measured historic and contemporary forest loss by integrating information from the Land Use Harmonization (*30*) and Global Forest Change (*23*) datasets. We also examined whether our results were robust to land use change data sources using the ESA Landcover (*28*) and KK09 (*29*) datasets (Figs. S4-S5). To test our predictions, we used a hierarchical Bayesian modelling framework, with individual time series nested within biomes (*36*) to account for the spatial structure of the data (see Methods for details).

Global biodiversity datasets have been criticized for spatial underrepresentation of sites modified by human activities (*37*). However, we found that over the duration of the time series in our study, population and biodiversity monitoring has occurred on sites which represent over half of the spectrum of global variation in forest cover gain and loss (Fig. 2B). Many forested ecosystems have undergone dramatic changes in the last two centuries, with all-time peak forest loss typically occurring in the early 1800’s, before biodiversity and population monitoring began, and before detailed satellite remote-sensing spatial layers are available (Fig. 2C). Nevertheless, the time periods and geographical locations sampled corresponded with intense transformations of forest landscapes in a high proportion of geographic locations (Fig. 2B).

Contrary to our first prediction (“*timing*”), we found that forest loss led to both increases and decreases in populations and biodiversity over time (Figs. 3-4 and S8-S9). Surprisingly, contemporary peak forest loss acted as a catalyst amplifying both positive and negative population and biodiversity change. Over time, contemporary peak forest loss intensified population declines, population increases and richness losses, but not richness gains, relative to the period before peak forest loss (Fig. 3A-D, F). In nearly a third of time series (32%), more than 10% of original species in the assemblage were replaced by new species (Fig. 3G-H). After peak forest loss, the assemblages that experienced the most richness change also experienced the most turnover (Pearson’s correlation = 0.37, 95% confidence intervals = 0.31 to 0.43). Across time series, more than half of all populations and ecological assemblages (61%) experienced higher rates of change after the largest forest loss event within each time series. However, forest loss events can occur at multiple time points at a given geographic location. Forest loss occurring outside of the period of population or biodiversity monitoring might influence our ability to detect a causal link between forest loss and biodiversity change (*16, 38*). For example, some sites experienced their largest reductions in forested areas over a century ago when we have very few ecological records. In a *post-hoc* analysis, we found that local-scale population declines were most pronounced when the monitoring occurred during the period of all-time peak forest loss (the 10-year period which included the largest reduction in forested areas at each site between 850 and 2015; Land-use Harmonization Dataset, Figs. 1B and 4A). Our findings of a full spectrum of population and biodiversity responses to forest loss highlight the complexity of real-world communities that might be overlooked in analyses that use a space-for-time substitution and thus do not capture the temporal dynamics of ecological responses to land-use change (*11, 12, 14, 39*).

**Fig. 3.**
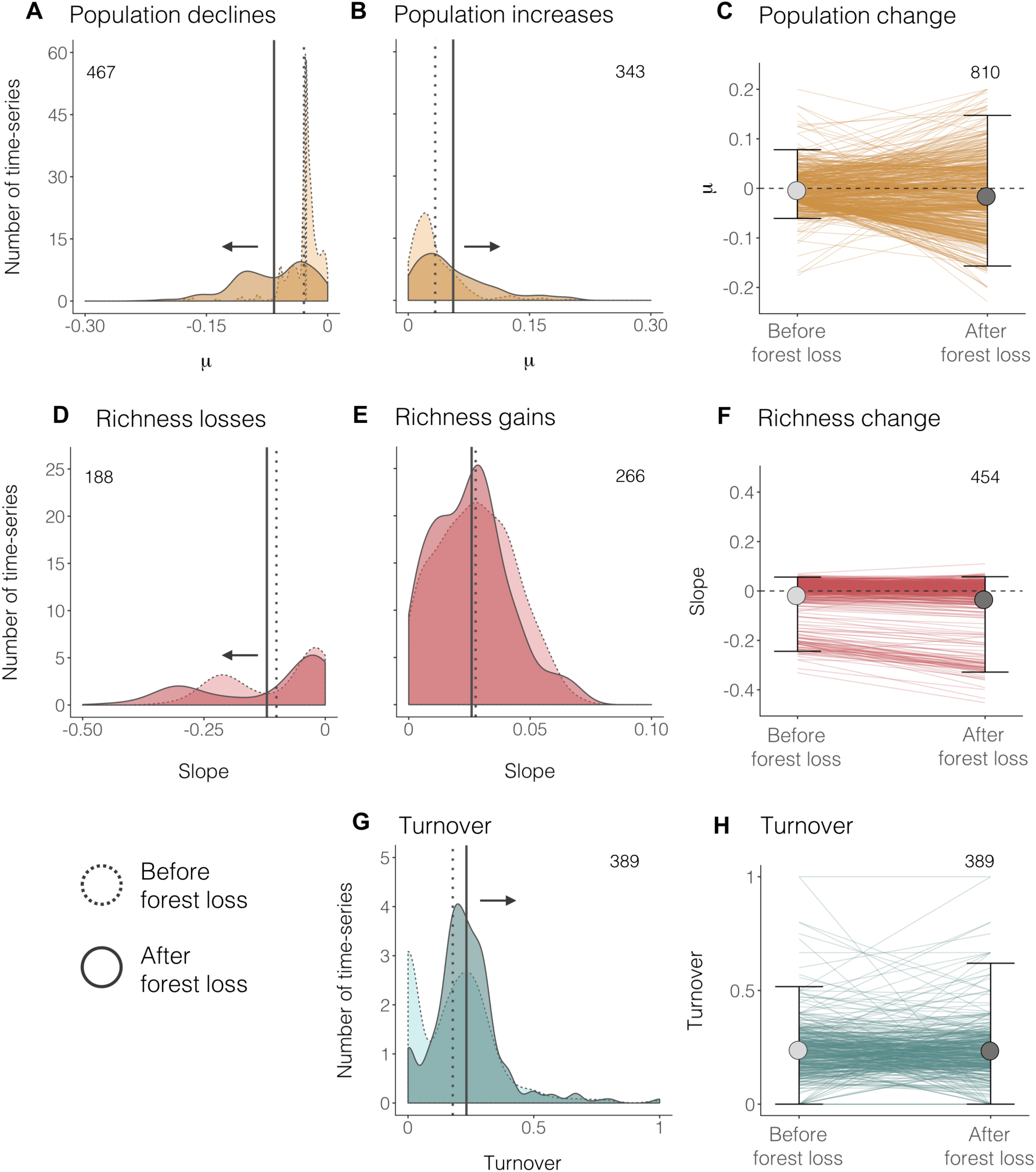
At the site level, population and biodiversity change increase after contemporary peak forest loss. In total, population, richness and turnover change increased across 61% and decreased across 39% of the 1,007 time series for which baseline comparisons were possible. Only turnover included instances of no difference in the amount of turnover before and after peak forest loss (6% of time series). Distributions compare **A**, population declines (*μ*), **B**, population increases (*μ*), **D**, richness losses (slopes), **E**, richness gains (slopes) and **G**, turnover (Jaccard’s dissimilarity) in the periods before and after peak forest loss. Vertical lines over distributions show the mean for each category (dotted – before; solid – after). Temporal change before and after contemporary peak forest loss, the largest forest loss event during the monitoring of each site, (**C, F, H)** is indicated with lines for individual time series. Duration varied among time series but was consistent for each individual time series (*i.e., n* years before forest loss = *n* years after forest loss, n ≥ 5 years). Light and dark grey points and error bars show mean values and 2.5 and 97.5% quantiles. Numbers on plots indicate sample size. See Table S1 for model outputs.

**Fig. 4.**
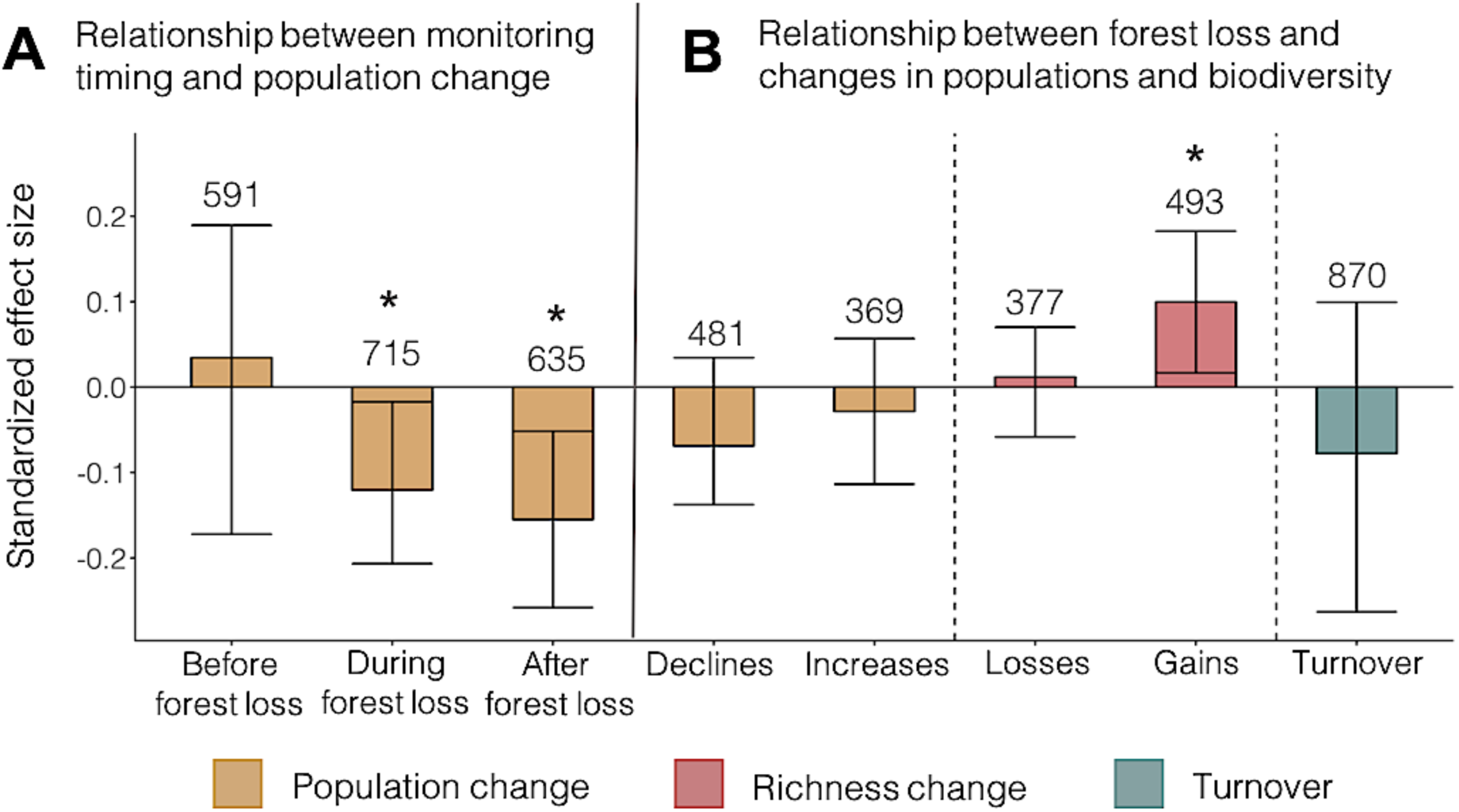
Population change, richness change and turnover across sites experiencing forest loss span a spectrum from increases to no net changes and declines. **A**, Population declines were most acute when all-time peak forest loss occurred during the population monitoring period (slope = -0.01, CI = -0.01 to -0.01;). Low sample size precluded a similar analysis for biodiversity change (Fig. S3B). **B**, The magnitude of forest loss experienced across sites over the duration of the GFC dataset (2000-2016) did not influence population declines and increases, richness losses, or turnover. Standardized effect sizes were calculated by dividing the model slopes by the standard deviation of the dependent variable. In approximately 5% of monitored time series, forest loss led to a conversion in the dominant habitat type (*e.g.*, from primary forest to urban areas). Habitat conversions corresponded with both gains and losses in population and biodiversity change, with turnover being highest when primary forests were converted to agricultural and urban areas (Fig. S12). For the effects of habitat transitions detected by the MODIS Land Cover Database (*24*), see Fig. S13. Numbers indicate sample size. For model visualizations, see Figs. S8-9.

Contrary to our second prediction (“*magnitude*”), we did not detect an effect of the magnitude of forest change on population and biodiversity losses. Greater magnitudes of forest loss did not correspond with larger increases in turnover or greater declines in populations and richness, and richness gains increased with forest loss (Figs. 4B and S9). Similarly, gains in forest cover did not correspond with gains in population abundance and species richness (Fig. S9). The decoupling between the magnitude of total forest loss across the time series duration and population and biodiversity declines could be due to a number of factors, such as: 1) temporal lags in population or community responses (*16, 30*), 2) lower forest loss during the monitoring period relative to historic clearing (*16, 29, 38*, Figs. S6-S7), 3) an influence of drivers other than forest loss, 4) variation in species vulnerability to disturbance (but note that generalist species were more likely to decline that forest-affiliated species, Fig. S8), and/or 5) mismatch between the assemblages monitored and the localized impacts of forest cover loss. It is also possible that forest loss amplified both positive and negative trends at local scales, but at larger scales favored the same species across sites leading to biotic homogenization (*41*). However, in our test of change in compositional similarity among sites between 1978 and 2007, we did not find evidence of biotic homogenization at monitored sites (Figure S10). Furthermore, rare and common species, as reflected by their range size, mean population size and habitat specificity, responded in similar ways to forest loss (Fig. S11). In contrast, space-for-time comparisons which do not account for temporal dynamics and lagged responses have found that land-use change impacts rare species more negatively than common species (*42*). Accounting for heterogeneity in the effects of forest cover change over time is key when scaling from localized impacts of human activities to global-scale biodiversity patterns and attribution of change (*1*).

In line with our third prediction (“*lags*”), we found evidence for up to half-century ecological lags in local-scale changes in population abundance, species richness and turnover following forest loss (Fig. 5). On average, we documented maximum change in populations and biodiversity six to 13 years after contemporary peak forest loss across taxa. Yet, nearly half of population and biodiversity change (40%) occurred within three years of peak forest loss, demonstrating that rapid shifts in populations and ecological assemblages occur frequently after habitat change (Fig. S14). As we predicted, the period between peak forest loss and peak change in populations and biodiversity was longer for taxa with longer generation times (*e.g.*, large mammals and birds, Fig. 5 and S14, Table S1). Population declines and increases occurred on similar timescales (Fig. S14). Losses in species richness lagged behind richness gains by only approximately half a year (slope = 0.5, CI = 0.1 – 1.05), indicating that extinction debts and immigration credits accumulated at roughly the same speed across taxa. The similar pace and temporal delay of population declines and increases, and richness gains and losses could be a contributing factor towards previous findings of no net population change (*2, 9, 43*) and richness change (*4, 5*) at local scales, in line with community self-regulation (*17*). Temporal lags in biodiversity change have also been observed in post-agricultural forests (*3, 44*) and fragmented grasslands (*30*), where agricultural activity has ceased decades to centuries ago, yet richness and community composition change continue to the modern-day. Overall, our results indicate that increasing rates of land-use change in the Anthropocene (*45, 46*) will alter ecosystems on both short- and long-term timescales that need to be captured in ongoing and future biodiversity monitoring.

**Fig. 5.**
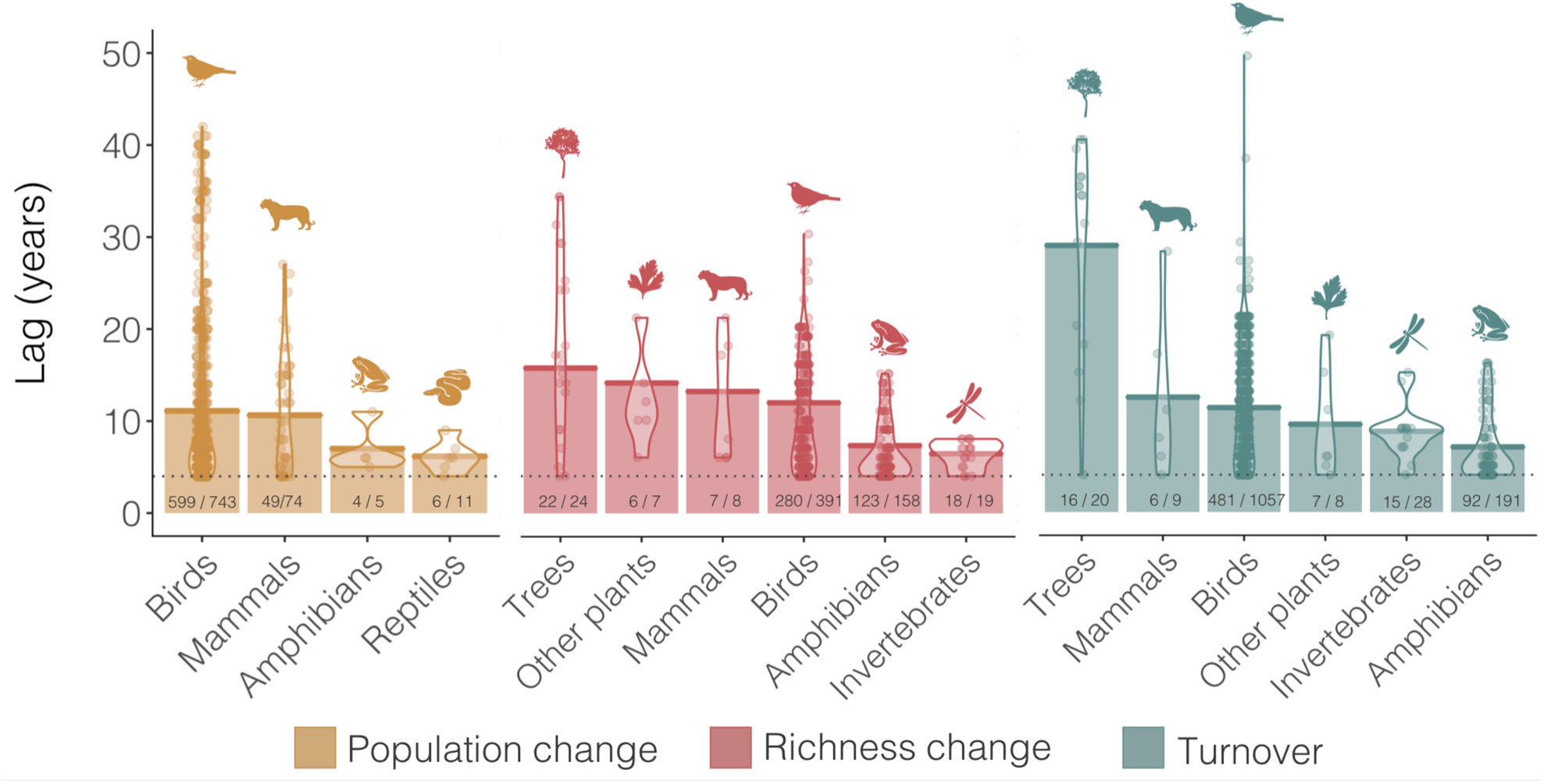
Population and community change after peak forest loss may be delayed by up to half a century, with taxa with long generation times showing the largest temporal lags. We categorized lags as time periods of three years (dashed horizontal line) or more between peak forest loss during the monitoring for each time series (from a total of 3,187 time series), and peak population/biodiversity change (Fig. 2B). Bars show mean lags for each taxon; violins show the distribution of lag values and the points are lag values for each time series. Numbers on bars indicate how many time series experienced lags out of the total sample size for each taxon. See Table S1 for model outputs.

Taxonomic, spatial and temporal imbalances in sampling can make large-scale attribution analyses of biodiversity trends and global change drivers challenging. Firstly, tropical species and locations are under-represented in current open-source temporal biodiversity databases (Fig. 2A). In a *post-hoc* test, we found that for the available tropical time series, the effects of forest loss are stronger and more negative relative to the rest of the globe (Fig. S15). Secondly, the spatial scales at which biodiversity is monitored and the resolution of forest cover datasets often differ by several orders of magnitude, which can introduce spatial mismatches between the driver and response. Nevertheless, we found that our richness change results were consistent across forest loss calculated on scales from 10km^2^ to 500km^2^ (Fig. S16A-B). Thirdly, temporal mismatches and lags (Figs. 1C and 5) can obscure relationships between forest loss and population and biodiversity change. We found that attribution signals were strongest when contemporary or all-time peak forest loss occurred during the monitoring time series (Figs. 3 and 4A). Biodiversity monitoring and global change attribution analyses will be improved by better spatial and temporal matching of biodiversity and environmental impact data.

In summary, our analysis reveals an intensification of both increases and decreases of populations and biodiversity by up to 48% at local scales after forest loss at sites around the planet. This finding challenges the widely-held assumption that land-use change universally leads to population declines and species richness loss (*11, 13, 39*). A current assumption underlying existing projections of biodiversity responses to land-use change (*11, 13*) is that space-for-time approaches accurately reflect longer-term population and biodiversity dynamics (*45*). In contrast, we found up to half of a century of temporal lags in population and biodiversity change following forest loss that varied by taxon and generation time. Our analyses highlight that real-world responses of population and assemblage to forest cover loss and gain are complex and variable over time. Forest loss was concurrent with both declines and increases in populations and ecological assemblages, similarly to the varied and often positive effects of habitat fragmentation on biodiversity metrics such as species richness (*18*). Our finding that forest cover gain does not correspond to gains in population abundance and species richness contribute to a growing body of literature indicating that afforestation efforts might have unintended biodiversity consequences (*47*), warranting caution with recent calls for global afforestation as a climate change mitigation tool (*21*). Incorporating the full spectrum of population and biodiversity change in response to land-use change will improve projections of future impacts of global change on biodiversity and thus contribute to the conservation of the world’s biota during the Anthropocene.

## Acknowledgments

We thank the WWF and ZSL for compiling the Living Planet Database, Robin Freeman and Louise McRae for useful discussions, the BioTime team for compiling the BioTime database (which was supported by ERC AdG BioTIME 250189 and ERC PoC BioCHANGE 727440), the creators of the Land Use Harmonization Database, The Hansen Lab for producing the Forest Cover Change Database and NASA for producing the MODIS Landcover Database. We thank Faye Moyes for managing the BioTIME database. We thank the Forest & Nature Lab at Ghent University for a stimulating discussion on historic and contemporary land-use change and choosing appropriate baselines for comparison of biodiversity change through time. We are grateful to Albert Phillimore and Kyle Dexter for providing advice during the conceptualization of the study, and to Laura Antão and Mark Vellend for providing feedback on the draft manuscript.

## Author contributions

G.N.D., M.A.D. and I.M.S. conceptualized the study. G.N.D. integrated databases and conducted statistical analyses with input from S.B., I.M.S., A.D.B. and M.A.D. G.N.D. created the figures with input from co-authors. S.B., M.A.D. and S.R.S. wrote the code for the rarefaction of the BioTIME studies. I.M.S. was the primary supervisor, M.A.D. the co-supervisor and A.D.B. is on the supervisory committee for G.N.D. M.A.D. and A.M. funded the compilation of the BioTime database. G.N.D. wrote the first draft and all authors contributed to revisions.

## Data and materials availability

Code for the rarefaction of the BioTIME Database is available from https://doi.org/10.5281/zenodo.1475218. Code for statistical analyses is available from http://doi.org/10.5281/zenodo.1490144. Population and biodiversity data are freely available in the Living Planet and BioTIME Databases (*25, 26*). The Living Planet Database can be accessed on http://www.livingplanetindex.org/data_portal. The BioTIME Database can be accessed on Zenodo (https://doi.org/10.5281/zenodo.1211105) or through the BioTIME website (http://biotime.st-andrews.ac.uk/). Land-use change data are publicly available in the Land Use Harmonization Database (*22*), the Forest Cover Change Database (*23*), and the MODIS Landcover Database (*24*).

