## Supplementary Information for "Landscape-scale forest loss as a catalyst of population and biodiversity change"

**This PDF file includes:**

Materials and Methods

Figs. S1 to S20

Tables S1 to S3

### Materials and Methods

For an illustration of the workflow of our analyses of forest cover and population and biodiversity change through time, see Figs. S1 and S2. All data and statistical analyses are described in detail below. We did not predetermine sample size and instead worked with all available temporal population, biodiversity and forest cover change data that met our duration criteria. For analyses of population change, we included time series with five or more survey points. For analyses of biodiversity change, we included time series with five or more data points when analyzing the full time series, and time series with two or more data points when matching the duration of time series comparisons to the 16-year duration of the Global Forest Change Database from 2000 to 2016. We calculated forest loss on a standardized landscape scale (~96 $km^2$). Our analyses were not sensitive to the cell size over which we calculated forest loss (tests from 10 $km^2$ to 500 $km^2$), as detected forest loss scaled proportionately with cell size across sites (Fig. S16A-B).

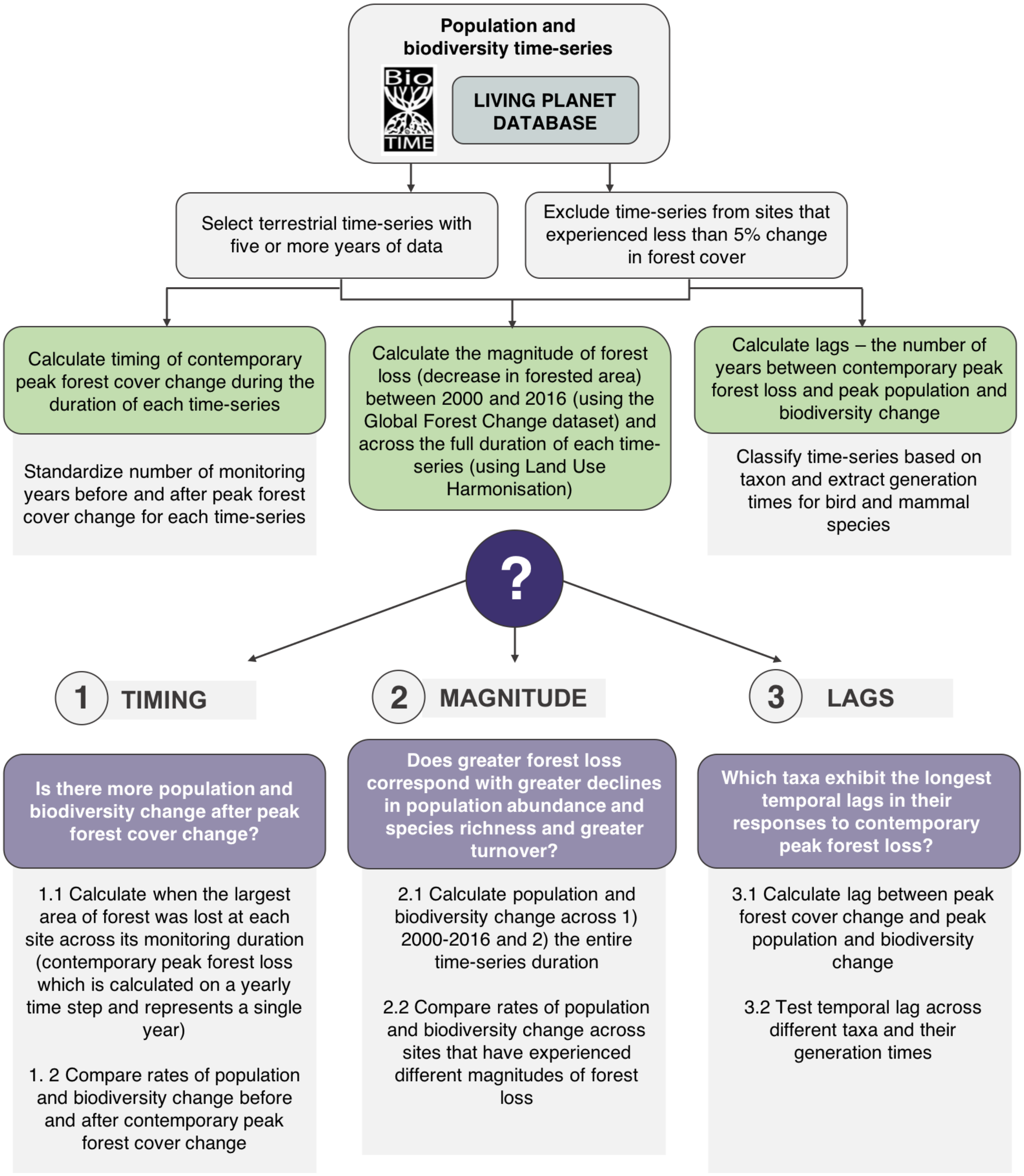


**Fig S1. Workflow of analyses to answer the following research questions: 1) Do population and biodiversity change increase after peak forest cover change? 2) Which type of habitat conversion yields the highest rates of population and biodiversity change? and, 3) Do lags in population and biodiversity responses to forest cover change vary across taxa?** See Fig. S2 for a worked example of our analyses for one time series and Table S1 for model outputs and sample sizes. For details on each step of our analyses, see correspondingly numbered sections in Methods and Materials.

**Databases**

*Forest cover change data*

To quantify historic and contemporary forest cover change, we extracted historic forest loss from the Land Use Harmonisation (LUH; 850 – 2015, forest loss and habitat transitions at a 0.25° degree resolution (around 20 km, *30*) and contemporary forest cover change and habitat conversions from the Global Forest Change (GFC, 2000 – 2016, forest loss and gain at a 30 m resolution, *21*), and MODIS Landcover (2000 – 2013, land-use transitions at 500m resolution, *22*) datasets.

*Land Use Harmonisation Database*

To estimate forest cover change across a time period matching the full duration of the biodiversity observations, we derived the change in primary forest cover from the Land Use Harmonisation database (LUH, *30*) for 96 $km^2$ cells around the location of each population in the LPD database (*25*) and for the standardized grid cells of the BioTIME database (~ 96 $km^2$ each, ). The v2h release of LUH includes annual gridded fractions of land-use states for the period from 850 to 2013 at 0.25° x 0.25° resolution. The estimates are based on historical reconstructions using Earth System models, with inputs such as regional and national rates of wood harvest and potential biomass density. The accuracy and precision of LUH increases towards the modern day, when there are more available data to inform the Earth System models. Note that unlike GFC, LUH estimates forest cover as a proportion (bounded between zero and one). For our analyses, we focused on time series from locations that have experienced at least 0.05 (equivalent to 5%) forest loss. To calculate total forest cover change over the period of a given population or biodiversity time series, we subtracted the proportion of forest cover in the first year of biodiversity monitoring from the proportion of forest cover in the last year. The type of forest cover change detected by the LUH database was predominantly forest loss, with forest gain occurring infrequently and at very small magnitudes (<0.001 out of maximum 1), thus we focus on forest loss.

To estimate the historic baseline of forest cover change, we calculated change in % forest cover across 10-year periods for each site from 850 to 2015 from the LUH data. We then determined all-time peak forest loss as the period when the most forest loss occurred (calculated using the difference in forest area at the start and end of standardized 10-year blocks). Time since all-time peak forest loss was a poor predictor of the variation in contemporary population and biodiversity change (Fig. S15F-H). To determine contemporary peak forest loss for each time series of monitoring data, we calculated yearly changes in forest cover across the duration of each time series and determined the year when the most change had occurred.

*Global Forest Change Database*

We derived overall forest loss and forest gain across the 2000-2016 period for 96 $km^2$ cells around the location of each population in the LPD database and for the standardized grid cells of the BioTIME database (~ 96 $km^2$ each) from the Global Forest Change (GFC, *21*) database using the Google Earth Engine (*48*). The GFC database provides high resolution forest cover change data, derived from Landsat satellite observations at a 30-meter spatial resolution. We calculated the total area of forest cover gain and loss separately (measured in $km^2$) for each 96 $km^2$ cell on a yearly time step. We then summed the yearly values for the period that coincided with population and biodiversity monitoring to estimate overall forest cover gain and loss (two separate metrics). For example, for a biodiversity time series spanning 2002 – 2009, our forest cover gain and loss metrics included the total amount of forest cover gained and lost during that same period. For our analyses, we focused on time series from locations that have experienced at least 0.5 $km^2$ of forest gain or loss. GFC does not distinguish between primary forest, secondary forest and plantations, but it does provide a very high-resolution measure of general forest cover. The drivers of the forest loss detected by GFC across our study sites are predominantly forestry, changes in agricultural practices and wildfires (*49*). Note that the GFC database spans from 2000 to 2016, whereas the earliest terrestrial biodiversity record in BioTIME is from 1858.

*MODIS Landcover Database*

To quantify habitat conversion for locations where we had population and biodiversity monitoring data, we used the MODIS Landcover Database (*24*). The MODIS Database has a resolution of 500 m, and it uses satellite-derived reflectance data to classify land cover around the world. To determine the types of habitat conversion between 2000 and 2013 (the time span of available MODIS data) across all monitoring locations, we calculated the dominant land cover type at the start and end of each population and biodiversity time series and split time series into categories such as "no habitat conversion" and "grassland to woody savannah". We focused on the eight most frequent types of habitat conversion (Fig. S13).

By synthesizing information from scenario data based on Earth Dynamics Models (LUH) and remote-sensing databases (GFC, MODIS), we were able to determine historic forest loss from the start of the monitoring period to 2016, as well as contemporary forest cover change (gain and loss) and habitat transitions from 2000 to 2016. GFC and MODIS detect forest cover, with no distinction between primary and secondary forests, thus we derived information on transitions from primary to secondary forest from the LUH database. We calculated overall forest cover change because we considered total change in habitat to be more meaningful for long-term population and biodiversity trends as opposed to an annual rate of forest cover change which does not capture cumulative effects. Together, the three databases (GFC, MODIS, LUH) encompass two different elements of land-use change: 1) land cover types and long-term historical reconstructions of past land-use and habitat conversions and 2) high-resolution satellite data from recent years of forest cover change and habitat conversion types. Thus, the combined analysis allows for a comprehensive test of the effects of land-use change on populations and biodiversity around the world.

*Population time series data (Living Planet Database)*

We analyzed 4,228 population time series with records distributed around the world. These time series represent repeated monitoring surveys of the number of individuals in a given area (species' abundance over time), to which we refer as "populations". Geographic representation is variable with, for example, an under-representation of tropical regions in the population data (Fig. 2A). In the LPD, some populations have precise coordinates, whereas the location of others are approximate. Because of the extent over which we are calculating forest cover change (96 km$^2$), we included both types of populations in our analysis. Duration varied across time series (Figs. S3C and S6) and we only included populations with at least five survey points. The overall range of the time series covered the period between the years 1970 and 2014. We calculated population change using state-space models which are particularly appropriate when quantifying change in data with varying collection methodology, as they take into account observation error and process noise (*50*, *51*). For more details on state-space model calculations, see Humbert *et al.* 2009 (*31*) and Daskalova *et al.* 2018 (*2*). We scaled the population size data to be between 0 and 1 to analyse within-population relationships and to make sure that we were not conflating within-population relationships and between-population relationships (*52*). State-space models partition the variance in abundance estimates into process error ($\sigma^2$) and observation or measurement error ($\tau^2$) and estimate population trends ($\mu$):

$$X_t = X_{t-1} + \mu + \varepsilon_t, (1)$$

where $X_t$ and $X_{t-1}$ are the scaled (observed) abundance estimates (between 0 and 1) in the present and past year, with process noise represented by $\varepsilon_t \sim gaussian(0, \sigma^2)$. We included measurement error following:

$$Y_t = X_t + F_t,$$

where $Yt$ is the estimate of the true (unobserved) population abundance with measurement error:

$$F_t \sim gaussian(0, \tau^2).$$

We substituted the estimate of population abundance ($Y_t$) into equation 1:

$$Y_t = X_{t-1} + \mu + \varepsilon_t + F_t.$$

Given $X_{t-1} = Y_{t-1} - F_{t-1}$, then:

$$Y_t = Y_{t-1} + \mu + \varepsilon_t + F_t - F_{t-1}.$$

For each time series, we calculated overall population change ($\mu$) experienced 1) across the periods before and after contemporary peak forest loss, 2) across the full duration of the time series, 3) from 2000 to 2016 (matching the temporal scale of the GFC database), and 4) from 2000 to 2013 (matching the temporal scale of the MODIS database). We standardized the number of years over which we calculated population change before and after peak forest loss on the population-level, meaning that the number of years before and after was the same within populations, but might differ among populations.

*Biodiversity time series data (BioTIME Database)*

We analyzed 2,339 time series from 190 studies from terrestrial biomes across the globe that make up a part of the BioTIME database (*25,* a full list of datasets including those the BioTIME database could not republish is included in Table S3). Similarly to the LPD, tropical regions and some taxa such as amphibians and reptiles were under-represented. Some of the study locations fall within protected areas (32%). Because those studies only had one time series each, overall only 1% of analyzed time series were from inside protected areas. To account for the different spatial extents of the BioTIME database, studies with multiple locations and extents > 72 $km^2$ were partitioned into 96 $km^2$ grids, and then sample-based rarefaction was applied to standardize sampling within each time series (*33*). The identity of each study was always kept intact, and we did not combine data from different studies, e.g., if there were data from two studies in the same cell, those represented separate time series. Duration varied across time series (Figs. S3D and S7) and the overall range of the time series covered the period between the years 1858 and 2016. For time series with five or more years of monitoring records, we calculated overall richness change and turnover experienced 1) across the periods before and after contemporary peak forest loss, 2) across the full duration of the time series. For time series with two or more years of monitoring records, we calculated overall richness change and turnover experienced 3) from 2000 to 2016 (matching the temporal scale of the GFC database), and 4) from 2000 to 2013 (matching the temporal scale of the MODIS database). The GFC and MODIS databases cover shorter time periods, thus we included biodiversity time series with shorter durations than the five-year cut off point that was used in the rest of our analyses using datasets with longer durations (but note that 76% of biodiversity time series had a duration of three or more years). To estimate richness change, we modelled species richness versus time (year, mean centered) with random slopes and intercepts for each rarefied cell and a Poisson error distribution with a log link.

$$log(\mu_{j,i,t}) = \beta_0 + \beta_{0j} + \beta_{0j,i} + (\beta_1 + \beta_{1j} + \beta_{1j,i})year_{j,i,t},$$

$$y_{j,i,t} \sim poisson(\mu_{j,i,t}),$$

where $year_{j,i,t}$ is the time in years, $\beta_0$ and $\beta_1$ are the global intercept and slope (fixed effects), $\beta_{0j}$ and $\beta_{1j}$ are the biome-level departures from $\beta_0$ and $\beta_1$ (respectively; biome-level random effects), $\beta_{0j,i}$ and $\beta_{1j,i}$ are the (nested) cell-level departures from $\beta_0$ and $\beta_1$ (cell-level random effects); $y_{j,i,t}$ is the (rarefied) species richness within the *jth* biome in the *ith* cell in year *t*.

From the richness over time model, we extracted the posterior means for richness change for each time series (i.e., the cell-level slope estimates), which then became the response variable in the second stage of our analyses where we tested richness change versus forest cover change (see Statistical analyses section).

To determine changes in community composition, we calculated the turnover component of beta diversity (changes due to species replacement rather than changes in species abundances (*32*, *33*), at the end of each time period outlined above relative to the first year of observation in the same period. Turnover is bound between zero and one, where zero is no change in species composition and one indicates that all of the original species of a community have been replaced with new species.

**Statistical analyses**

We always matched the temporal scales of the forest cover change data and the population and biodiversity data when investigating attribution signals (i.e., evidence that a predictor variable is a potential driver of population or biodiversity change). For example, when testing the effects of forest cover change and land-use transitions as detected by GFC (2000 – 2016) and MODIS (2000 – 2013), we calculated population and biodiversity change for the same time periods. Because of the longer duration of the LUH database, we were also able to extract forest and land cover information for the full duration of the LPD and BioTIME time series. We excluded locations which had less than 0.05 (out of maximum 1) forest cover change from our analyses of contemporary peak forest loss and overall forest loss (using the LUH database over a time period matching the duration of each time series). We excluded locations which had no forest cover across the duration of the time series in both the 96 $km^2$ cells and the 500 $km^2$ larger landscape cells from our analyses of population and biodiversity change versus forest cover gain and loss from 2000 to 2016 (using the GFC database). See Figure S2 for an example workflow and summary of all analyses done and Table S1 for the outputs of all statistical models and their respective sample sizes.

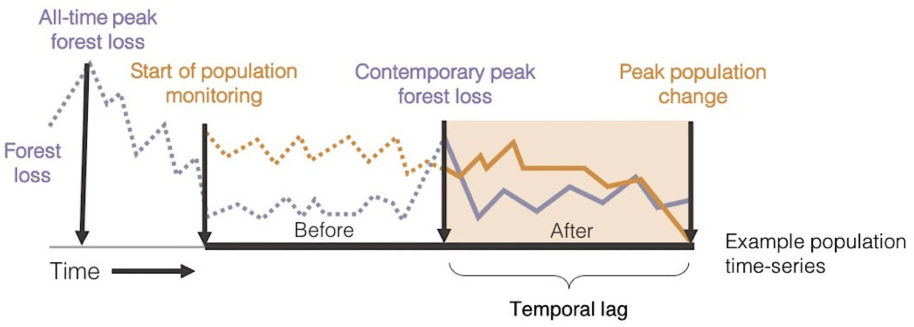


**1** Compare population trend before and after contemporary peak forest loss

**2** Model population trend as a function of the magnitude of forest loss. Forest loss refers to the total area of forested habitat lost 1) between 2000 and 2016 (to match GFC dataset) and 2) across the entire duration of each time-series. We always matched the temporal span of the population, biodiversity and forest cover data.

**3** Calculate temporal lag between peak contemporary forest loss and peak poulation change and compare it across time-series from different taxa

**4** Post-hoc analyses:

1. Are populations more likely to experience declines when the monitoring includes the period of all-time peak forest loss (the largest reduction in forest area between 850 and 2015)?
2. Within bird and mammal taxa, do species with longer generation times experience longer lags in population change following contemporary peak forest loss?
3. Are rare species more likely to respond negatively to forest loss compared to common species?
4. Is compositional similarity among sites increasing over time, in line with biotic homogenization?

**5** Sensitivity analyses:

1. Does calculating turnover relative to the first year of the time-series influence the amount of detected change in community composition?
2. How does forest loss influence population change when analyzing the BioTIME database on a population level?
3. How does nesting time-series id within study id influence the amount of detected richness change over time?
4. Are the effects of forest loss on population and biodiversity change in the tropics different relative to the rest of the world?
5. Is there a taxonomic signal in the relationships between forest loss and population change, richness change and turnover?
6. Are species associated with forests more likely to be negatively influenced by forest loss?
7. Does the spatial scale over which we quantify forest cover change influence its detected effects on population and biodiversity change?
8. Does the landscape context of forest loss (amount of forested area in 500km$^2$ cell around each time-series location) influence the relationships between forest loss and population and biodiversity change?
9. Does the type of forest (primary vs. secondary forest) influence the amount of detected forest cover change?
10. Does the amount of detected forest cover change vary depending on data sources?

**Fig. S2. Example workflow for one time series including main research questions, post-hoc analyses and sensitivity analyses.** See Fig. S1 for the overall workflow of analyses. The Living Planet Database includes repeated monitoring surveys of the number of individuals in a given area (species' abundance over time), to which we refer as "populations". See Fig. S2 for a worked example of our analyses for one time series and Table S1 for model outputs and sample sizes. For details on each step of our analyses and post-hoc analyses, see correspondingly numbered sections in Methods and Materials.

**1. Timing**

To test if temporal population and biodiversity change differed before and after peak forest loss at the site-level, we split each time series into two periods – before and after peak deforestation – and estimated population change, richness change and turnover for each period separately. Then, to infer if population and biodiversity change differed following peak forest loss, we modelled $\mu$ (population change), richness change (cell-level random slopes) and turnover as a function of period (categorical with two levels – before or after forest loss) and time series duration (numeric) as fixed effects, with a biome random effect to account for the spatial clustering of the data. For population and richness change, we modelled the positive and negative components of the distributions of change separately, e.g., one model for populations with positive $\mu$ values and one model for populations with negative $\mu$ values. This approach allowed us to test if the effects of forest loss differ across the positive and negative dimensions of population and biodiversity change. The models were as follows:

$$\mu_{j,i,p} = \beta_0 + \beta_{0j} + \beta_1 * duration_{j,i,p} + \beta_2 * period_{j,i,p},$$

$$y_{j,i,p} \sim gaussian(\mu_{j,i,p}, \sigma^2),$$

where $duration_{j,i,p}$ is the duration of the time series in years of cell $i$ within biome $j$ for period $p$, and $period_{j,i,p}$ is an indicator variable for the period (before or after forest loss); $\beta_0$, $\beta_1$ and $\beta_2$ are the global intercept and slope estimates for duration and the categorical period effect, respectively (fixed effects), $\beta_{0j}$ is the biome-level departures from $\beta_0$ (biome-level random effects); $y_{j,i,p}$ is the estimate for change in population size or species richness for the *ith* cell in the *jth* biome for the *pth* period.

To model the change in turnover before and after contemporary peak forest loss, we followed the same conceptual framework as outlined above, but we used a zero one inflated beta distribution to account for the properties of turnover (bounded between zero and one, inclusive, where one is a complete change in species composition). The probability density function for the zero one inflated beta distribution is:

$$betainf(y;\alpha,\gamma,\mu,\phi) = \begin{cases} \alpha(1-\gamma), & y = 0 \\ \alpha\gamma, & y = 1 \\ (1-\alpha)\gamma f(y;\mu,\phi), & 0 < y < 1, \end{cases}$$

where $\alpha$ is the probability that a zero or one occurs, $\gamma$ is the probability that a one occurs (given an observation is a zero or a one), and $\mu$ and $\phi$ are the mean and precision of the beta distribution, respectively. In the parameterization approach we used (*53*) $\phi$ is inversely related to the variance. Beta parameterization is also sometimes expressed through the parameters $p$ and $q$ that can be derived from our framework following $\phi = p + q$ (*54*). Because only 7% of time series did not experience any change in species composition ($y = 0$) in the time period after contemporary forest loss, and less than 1% of time series had a completely new set of species ($y = 1$) occupying the ecological communities, for $y = 0$ and $y = 1$, $\alpha$ and $\gamma$ were modelled assuming a Bernoulli distribution and logit-link function, and models were fit with only an intercept. For $0 < y < 1$, we assumed a beta error distribution and a logit-link function:

$$logit(\mu_{j,i,p}) = \beta_0 + \beta_{0j} + \beta_1 * duration_{j,i,p} + \beta_2 * period_{j,i,p},$$

$$y_{j,i,p} \sim Beta(\mu_{j,i,p}, \phi),$$

where $duration_{j,i,p}$ is the duration of the time series in years of cell $i$ within biome $j$ for period $p$, and $period_{j,i,p}$ is an indicator variable for the period (before or after forest loss); $\beta_0$, $\beta_1$ and $\beta_2$ are the global intercept and slope estimates for duration and the categorical period variable respectively (fixed effects), and $\beta_{0j}$ are the biome-level departures from $\beta_0$ (biome-level random intercepts); $y_{j,i,p}$ is the estimate of turnover for the *ith* cell in the *jth* biome for the *pth* period.

**2. Magnitude**

To test the effect of forest cover change on population and biodiversity change among sites, we modelled population and biodiversity change versus overall forest cover change (calculated as forest cover gain and forest cover loss (GFC database, 2000-2016) and forest loss (LUH database, across the duration of the time series). Models of population and richness change versus forest cover change were fitted assuming Gaussian error.

$$\mu_{j,i} = \beta_0 + \beta_{0j} + \beta_1 * duration_{j,i} + \beta_2 * forest\ change_{j,i},$$

$$y_{j,i} \sim gaussian(\mu_{j,i}, \sigma^2),$$

where $duration_{j,i}$ is the duration of the time series in years of cell $i$ within biome $j$, $forest\ change_{j,i}$ is the forest cover change in cell $i$ within biome $j$; $\beta_0$, $\beta_1$ and $\beta_2$ are the global intercept and slope estimates for duration and forest cover change respectively (fixed effects), and $\beta_{0j}$ are the biome-level departures from $\beta_0$ ( biome-level random intercepts); $y_{j,i}$ is the population or richness change metric (a separate model for population declines, population increases, richness losses and richness gains) in the *ith* cell within the *jth* biome.

Models of turnover versus forest cover change were fit with a zero one inflated beta distribution to account for the properties of turnover (bounded between zero and one). We used the same probability density function for the zero one inflated beta distribution as in the model for turnover before and after contemporary peak forest loss. For $y$ = 0 and $y$ = 1, $\alpha$ and $\gamma$ were modelled assuming a Bernoulli distribution and logit-link function, and we fit models with only an intercept. For 0 < $y$ < 1, we assumed a beta error distribution and a logit-link function:

$$logit(\mu_{j,i}) = \beta_0 + \beta_{0j} + \beta_1 * duration_{j,i} + \beta_2 * forest\ change_{j,i},$$

$$y_{j,i} \sim Beta(\mu_{j,i}, \phi),$$

where $duration_{j,i}$ is the duration of the time series in years of cell *i* within biome *j*, $forest\ change_{j,i}$ is the forest cover change in cell *i* within biome *j*; $\beta_0$, $\beta_1$ and $\beta_2$ are the global intercept and slope estimates for duration and forest cover change respectively (fixed effects), and $\beta_{0j}$ are the biome-level departures from $\beta_0$ (biome-level random intercepts); $y_{j,i}$ is turnover in the *ith* cell within the *jth* biome.

*Habitat conversion and population and biodiversity change*

To determine the influence of the type of forest cover change (i.e., land-use transitions) on population and biodiversity change, we compared the distributions of population and biodiversity change across transitions types (from primary forest to secondary forest, from primary forest to non-natural habitat, and from secondary forest to non-natural habitat, to which we refer as habitat conversion). Small sample sizes (on average 10 time series per transition type) precluded statistical analysis, thus we report findings from a visual inspection of distributions of population and biodiversity change across habitat conversion types.

**3. Lags**

To test for temporal lags in population and biodiversity responses to contemporary peak forest loss, we first calculated when population and biodiversity change were greatest following peak forest loss for each time series. Rates of population change were calculated using state-space models and a Kalman filter (*9*, *31*). Peak richness change and peak turnover were calculated as the maximum value of the absolute differences between consecutive observations of species richness and turnover. We then quantified lag as the number of years between contemporary peak forest loss and peak population/biodiversity change. We modelled lag as a function of taxa, as we expect that species with longer generation times will respond to disturbance more slowly.

$$\mu_{j,i} = \beta_{0j} + \beta_1 * taxa_{j,i},$$

$$y_{j,i} \sim gaussian(\mu_{j,i}, \sigma^2),$$

where $taxa_{j,i}$ is the taxa of the cell $i$ in the biome $j$ time series, $\beta_1$ is the slope for taxa effect (fixed effect), and$\beta_{0j}$ are the biome-level random intercepts; $y_{j,i}$ is the temporal lag in the population or biodiversity change metric (a separate model for population change, richness change and turnover) for the *ith* cell within the *jth* biome.

*Prior specification*

We used weakly regularizing normally-distributed priors for the global intercept and slope for all models except the model of turnover versus overall forest cover change (which was a zero one inflated model):

$$\beta_0 \sim gaussian(0, 6),$$

$$\beta_1 \sim gaussian(0, 6).$$

For the turnover models that had a zero one inflated beta distribution, we used the following priors:

$$\beta_0 \sim gaussian(0, 6),$$

$$\beta_1 \sim gaussian(0, 6),$$

$$zoi \sim gaussian(0, 0.5),$$

$$coi \sim gaussian(0, 0.5),$$

where *zoi* is the probability of being a zero or a one and *coi* is the conditional probability of being a one (given an observation is a zero or a one).

Group-level parameters (the rarefied cell random effect in the species richness over time model, *i*, and the biome random effect in all models, *j*) were all assumed to be *gaussian*(0, σ), and priors on the σ were the same for all models:

$$\sigma\beta_{0j} = \sigma\beta_{0j,i} \sim half\ \ Cauchy(0, 2).$$

All models were fitted in a Bayesian framework using the *brms* package v2.1.0 (*53*) in R v3.5.1 (*55*). Models were run for 6000 iterations, with a warm up of 2000 iterations. Convergence was assessed visually by examining trace plots and using *Rhat* values (the ratio of the effective sample size to the overall number of iterations, with values close to one indicating convergence).

**4. Post-hoc analyses**

*4.1 Are populations more likely to experience declines when the monitoring includes the period of all-time peak forest loss (the largest reduction in forest area between 850 and 2015)?*

To determine if population change differed based on whether population time series were recorded before, during, or after the period of all-time peak forest loss, we modelled μ (population change) as a function of when monitoring started. We defined all-time peak forest loss as the timing of the largest forest loss event at the location of each time series between the years 850 and 2015. We used a categorical variable with three levels – before, during or after peak forest loss – and time series duration (numeric) as fixed effects, with a biome random effect to account for the spatial clustering of the data. Low sample size precluded a similar analysis for biodiversity change (Fig. S3B). The model was as follows:

$$\mu j, i, m = \beta 0 + \beta 0 j + \beta 1 * duration j, i, m + \beta 2 * monitoring\ start j, i, m,$$

$$y j, i, m \sim gaussian(\mu j, i, m, \sigma 2),$$

where $duration_{j,i,m}$ is the duration of the time series in years of cell i within biome j for monitoring start m, and monitoring $start_{j,i,m}$ is an indicator variable denoting when monitoring started; $\beta_0$, $\beta_1$ and $\beta_2$ are the global intercept and slope estimates for duration and the categorical monitoring start variable respectively (fixed effects), $\beta_{0j}$ is the biome-level departures from $\beta_0$ (respectively; biome-level random effects); $y_{j,i,m}$ is the estimate for change in population size or species richness for the ith cell in the jth biome for the mth monitoring start.

*4.2 Within bird and mammal taxa, do species with longer generation times experience longer lags in population change following contemporary peak forest loss?*

We conducted a post-hoc analysis where we tested our temporal lag and generation time hypothesis in a more quantitative manner by modelling lag as a function of generation time in birds and mammals, the taxa for which generation time data were freely available (9, 52).

$$\mu g = \beta 0 + \beta 1 * generation\ time g,$$

$$y_g \sim gaussian(\mu_g, \sigma^2),$$

where generation $time_g$ is the mammal generation time in years, $\beta_0$ and $\beta_1$ are the global intercept and slope (fixed effect); $y_g$ is the temporal lag in population change for a species with generation time g.

*4.3 Are rare species more likely to respond negatively to forest loss compared to common species?*

We tested if rare species (based on species' geographic range, mean population size and habitat specificity) were more likely to be negatively influenced by forest loss by including a forest loss * rarity metric interaction term in our models (for full details on methods to determine rarity, see (*2*)). We found that regardless of whether species were rare or common, they experienced the full spectrum of forest loss effects (Figure S11, Table S1). Similar post-hoc analysis was not possible for the biodiversity time series because habitat preference and rarity data were not available for many of the species included in the BioTIME database.

*4.4 Is compositional similarity among sites increasing over time, in line with biotic homogenization?*

To test if forest loss favors or negatively influences the same kinds of species, we calculated spatial patterns of assemblage composition among sites within studies for 384 time series at two time points using Jaccard's dissimilarity (1978 and 2007). We found that over time, compositional dissimilarity has slightly increased, but overall, for these monitored sites, there was no support for biotic homogenization (Fig. S10).

### 5. Sensitivity analyses

*5.1 Does calculating turnover relative to the first year of the time series influence the amount of detected change in community composition?*

Our analyses were not sensitive to our calculation of turnover in the final year of the time series relative to the first year, and previous examinations of the BioTIME database have found that calculating turnover relative to the second year of observation produced similar results (*5*).

*5.2 How does forest loss influence population change when analyzing the BioTIME database on a population level?*

We quantified population change using the BioTIME database (following the same state-space modelling framework as with the LPD) and found similar lack of directional patterns in the relationships between population change and overall forest loss (Fig. S8F).

*5.3 How does nesting time series id within study id influence the amount of detected richness change over time?*

When we calculated slopes of richness change over time, we included a time series ID random effect. In a small set of occasions (28%), there were more than one time series from the same study because the studies covered very large areas (e.g., the whole USA). Calculating richness slopes with a model including a nested random effect (time series id within study) produced very similar estimates (Fig. S17).

*5.4 Are the effects of forest loss on population and biodiversity change in the tropics different relative to the rest of the world?*

When testing for geographic variation in the effects of forest loss, we found that the effects of forest loss were more likely to be negative in the tropics relative to the rest of the globe (Fig. S15).

*5.5 Is there a taxonomic signal in the relationships between forest loss and population change, richness change and turnover?*

We found no distinct taxonomic patterning in the relationships between population change, biodiversity change and forest cover change (Fig. S18). A very small proportion (around 1%) of the species represented in our analyses were classified as invasive or alien (based on the Global Invasive Species Database, (*57*), with around 3% of species identified as only morphospecies for which we cannot attribute species status (Fig. S19).

*5.6 Are species associated with forests more likely to be negatively influenced by forest loss?*

The relationships between population decreases and increases and forest loss were not influenced by whether species were tightly associated with forests or not (Fig. S8G-I).

*5.7 Does the spatial scale over which we quantify forest cover change influence its detected effects on population and biodiversity change?*

The cell size over which we calculated forest cover change (from 10 $km^2$ to 500 $km^2$) did not influence overall findings, as detected forest cover change scaled proportionately with cell size across locations (Fig. S16A-B).

*5.8 Does the landscape context of forest loss (amount of forested area in 500km² cell around each time series location) influence the relationships between forest loss and population and biodiversity change?*

Landscape context did not influence the relationship between forest cover change and population declines (Fig. S16C-D), but there were less richness losses when the landscape-scale forest cover was higher (Fig. S20).

*5.9 Does the type of forest (primary vs. secondary forest) influence the amount of detected forest cover change?*

Our findings were not influenced by the type of forest cover (primary vs secondary), as loss of secondary forest cover scaled proportionately to primary forest loss (Fig. S16E).

*5.10 Does the amount of detected forest cover change vary depending on data sources?*

Uncertainty in forest cover estimates increases for records further into the past. On a European scale, we were able to compare historic land cover estimates as quantified by the LUH and KK09 databases and found that they produced broadly consistent estimates (Fig. S4E). We then tested how the amount of detected forest loss varies across two remote-sensing datasets (GFC and ESA Landcover) for the time period over which they overlap (2000-2015) and found frequent mismatches in the magnitude of detected forest cover change (Fig. S4A-D and S5). In our analyses, we focus on using the Land Use Harmonization dataset because it includes the longest possible temporal records of forest loss, and the Global Forest Change dataset, because it provides the highest spatial resolution for forest cover change currently available. Our findings of both positive and negative associations of population and biodiversity change with forest cover change were broadly consistent regardless of the dataset with which forest cover was calculated.

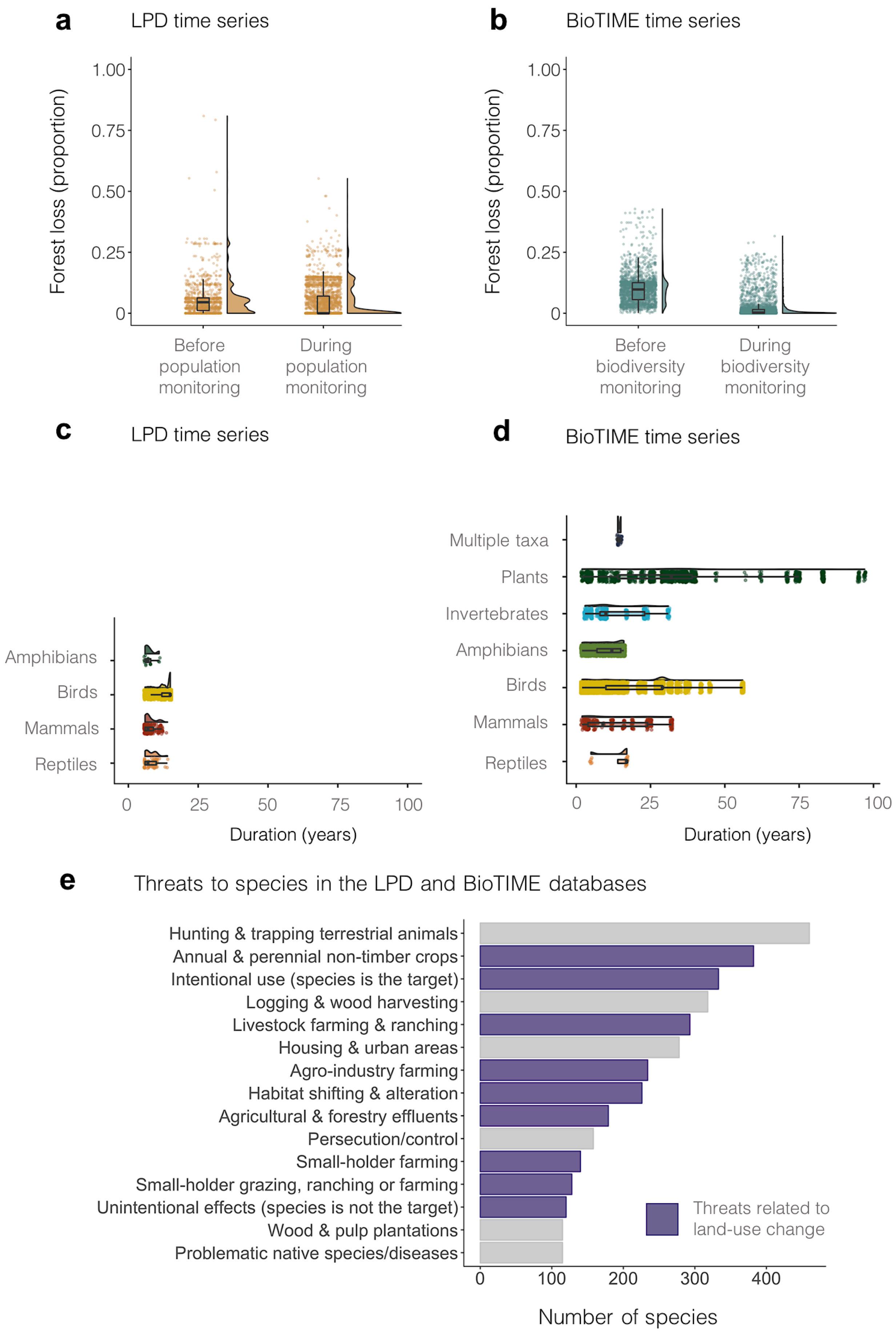
a
LPD time series
Forest loss (proportion)
1.00
0.75
0.50
0.25
0
Before population monitoring
During population monitoring
b
BioTIME time series
Forest loss (proportion)
1.00
0.75
0.50
0.25
0
Before biodiversity monitoring
During biodiversity monitoring
c
LPD time series
Amphibians
Birds
Mammals
Reptiles
0
25
50
75
100
Duration (years)
d
BioTIME time series
Multiple taxa
Plants
Invertebrates
Amphibians
Birds
Mammals
Reptiles
0
25
50
75
100
Duration (years)
e
Threats to species in the LPD and BioTIME databases
Hunting & trapping terrestrial animals
Annual & perennial non-timber crops
Intentional use (species is the target)
Logging & wood harvesting
Livestock farming & ranching
Housing & urban areas
Agro-industry farming
Habitat shifting & alteration
Agricultural & forestry effluents
Persecution/control
Small-holder farming
Small-holder grazing, ranching or farming
Unintentional effects (species is not the target)
Wood & pulp plantations
Problematic native species/diseases
Threats related to land-use change
0
100
200
300
400
Number of species

**Fig. S3. On average, there was more forest loss before population and biodiversity monitoring started compared to during the monitoring period, but the dominant threats to the species represented in the databases were related to land-use change.** Forest loss in **a** represented declines in primary forest cover for each time series, calculated using the LUH database (*22*) (cell size = 96 $km^2$). Population **c** and biodiversity **d** monitoring duration varied across taxa, but always included five or more data points per population time series, and five or more data points per biodiversity time series when analyzing the effects of contemporary peak forest loss and forest loss across the whole duration of the time series. When analyzing the effects of forest cover gain and loss (derived using the Global Forest Change Database which spans the period between 2000 and 2016), we included biodiversity time series with two or more records, with 76% or the time series having three or more records. Note that the locations of the reptile biodiversity time series had not experienced forest loss greater than 5% of the study cell and were thus not included in the subsequent analyses of forest loss effects on biodiversity change. Threats to species in **e** were extracted from the IUCN Red List Database (*29*) using the package rredlist v0.5.0 (*58*) for the 755 species for which global threat assessments were available. Boxplots in **a**-**d** show the mean, with the whiskers extending to the mean ± SD.

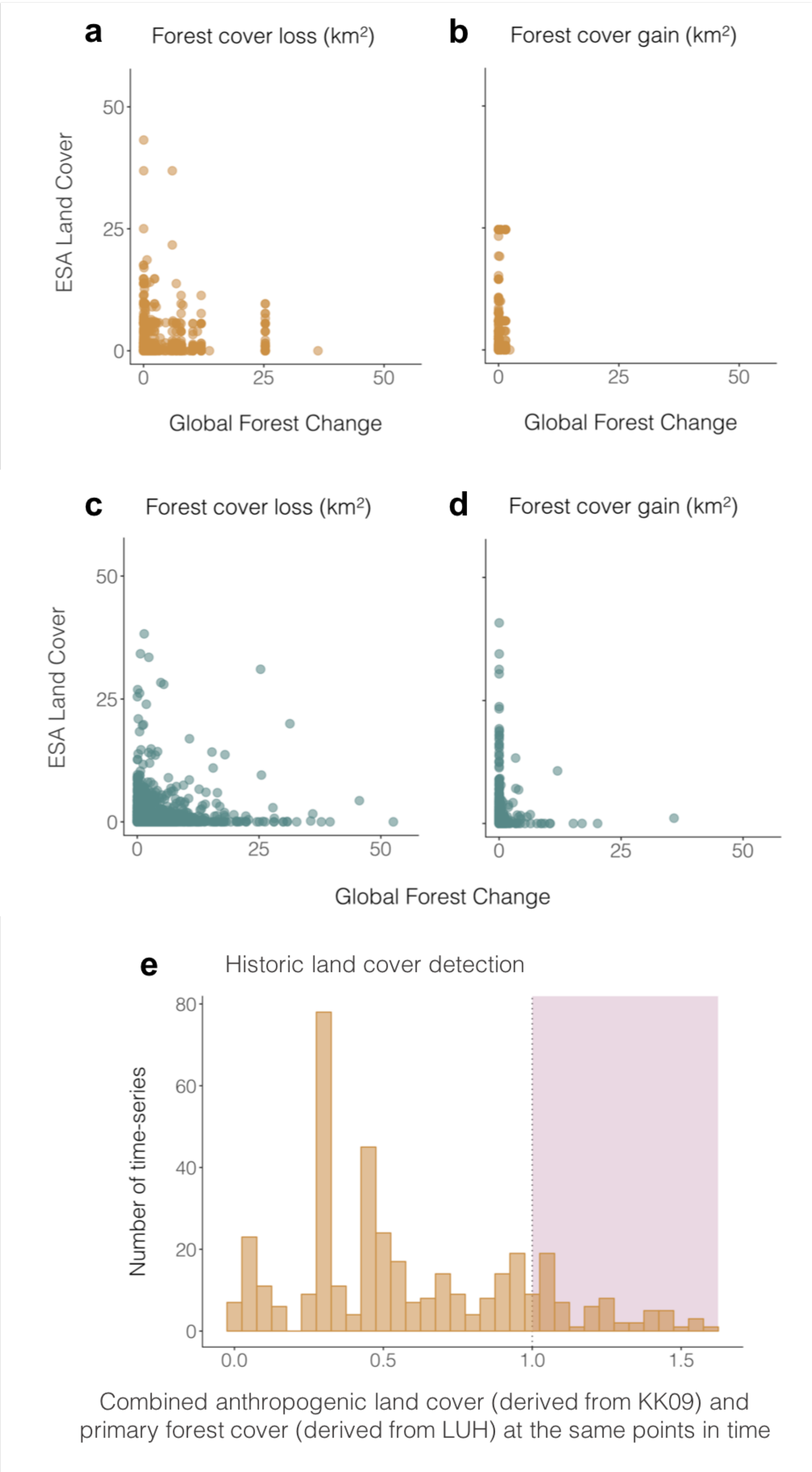
a
Forest cover loss (km²)
b
Forest cover gain (km²)
ESA Land Cover
50
25
0
0
25
50
Global Forest Change
Global Forest Change
c
Forest cover loss (km²)
d
Forest cover gain (km²)
ESA Land Cover
Global Forest Change
e
Historic land cover detection
Number of time-series
80
60
40
20
0
0.0
0.5
1.0
1.5
Combined anthropogenic land cover (derived from KK09) and
primary forest cover (derived from LUH) at the same points in time

**Fig. S4. There are mismatches in the amount of forest cover change detected by different global gridded datasets.** We calculated how much forest cover loss and gain are detected during the same time period (2000 – 2015) across sites with long-term monitoring of species' population abundance (**a**-**b**) and richness and composition of ecological assemblages (**c**-**d**). We compared the forest cover gain and loss estimates from two datasets – the Global Forest Change dataset (*23*) and the ESA Landcover dataset (*34*). While Global Forest Change focuses exclusively on forests, ESA Landcover provides estimates for 37 different land cover classes. Thus, the potential explanations for the mismatches between the two datasets include: 1) different definitions of a forest, 2) a ten-fold difference in resolution (30 m for Global Forest Change versus 300 m for ESA Landcover, and 3) differences in satellite data sources and processing methods. We also derived estimates for historic anthropogenic land cover in Europe from the KK09 dataset (*35*) and tested how they compare with the estimates for primary forest cover derived from the Land Use Harmonization dataset (*22*). We derived these estimates for the first year of each population time series for the same sized cells (~96 $km^2$) and combined them. This combined measure is shown on the x-axis of plot **e**. It is possible for the two estimates to be below one (suggesting at a given location there is some land under anthropogenic land use, some primary forest, but also another land cover type). Estimates above one indicate instances where there might be an error in either of the two datasets as the combined area of anthropogenic and forest cover in a cell cannot go above one in reality.

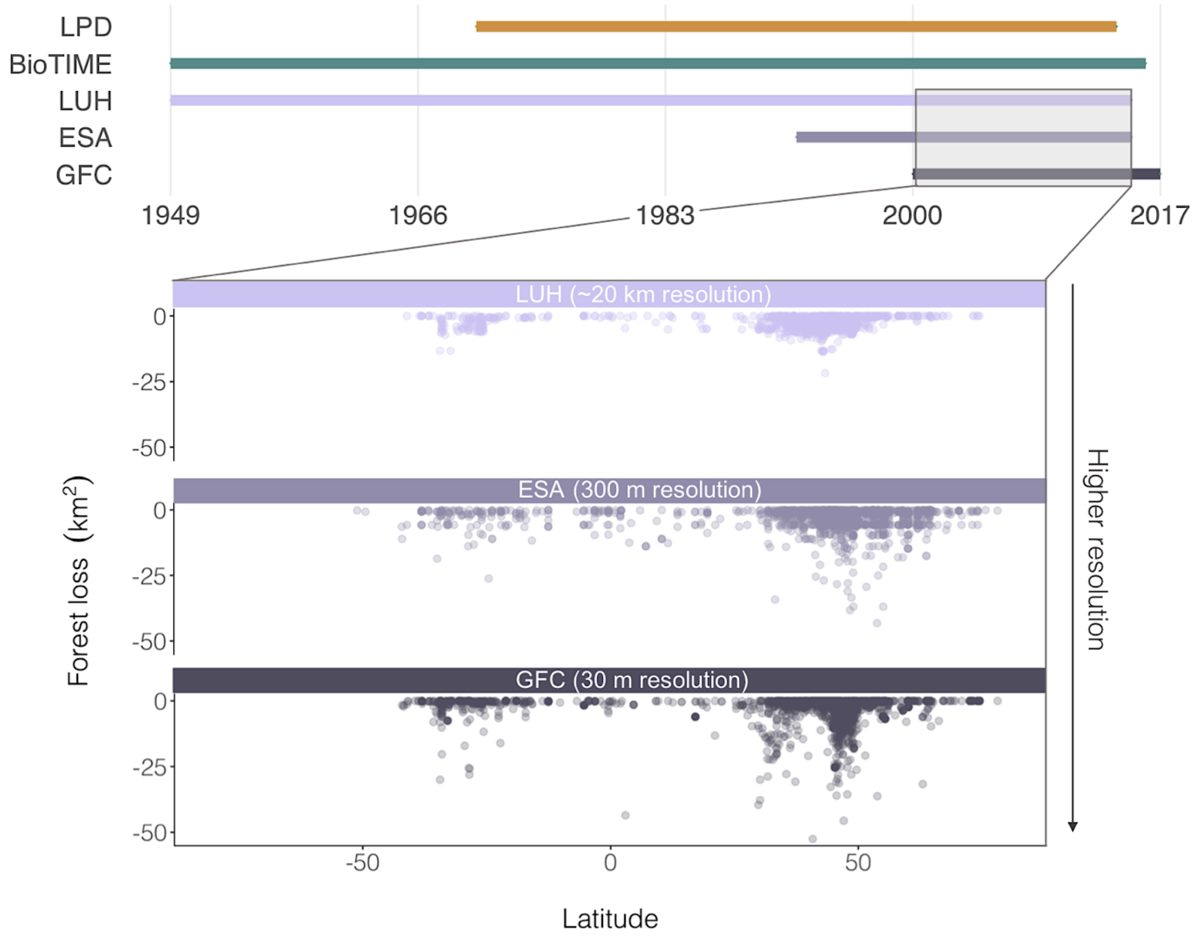


**Fig. S5. Population, biodiversity and forest cover databases cover different periods in time and the amount of detected forest loss increases with higher resolution of the spatial data.** The forest datasets used to calculate forest loss were the Land Use Harmonization dataset, the ESA Landcover dataset and the Global Forest Change dataset. Direct comparisons of detected forest loss and gain across sites are shown in Fig. S4.

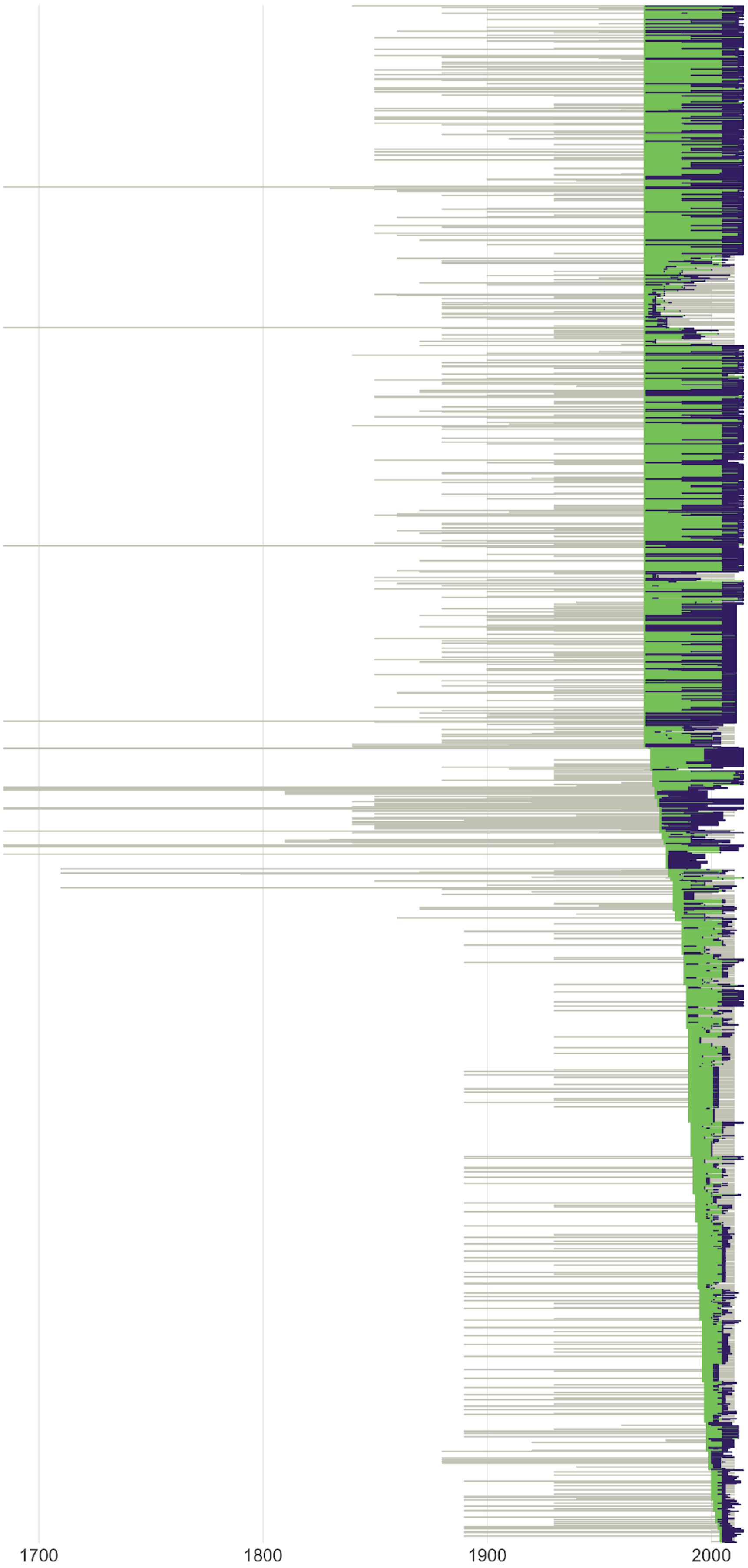
1700
1800
1900
2000

**Fig. S6. Historic peaks of forest cover change often occur decades to centuries before population monitoring starts (38% of time series), with some instances (23% of time series) of peaks in forest cover change occurring after population monitoring has ceased.** Each line represents a single population time series, part of the Living Planet Database (*24*). Grey lines start at the historic peak of forest cover change for each time series. The detected forest cover change constituted declines in forest cover, calculated based on the LUH database (*22*). Green and purple lines show the duration of population monitoring, with the break between green and purple indicating the largest forest cover change event across the duration of each time series. Note that for analysis, we compared population change across equal durations before and after recent peak forest cover change. E.g., if a time series included seven years of population abundance data before the peak in recent forest cover change and 10 years of data after that peak, we included only the first seven years of population data after the forest cover change peak (making for a total of 14 years of data included in the analysis for this example time series).

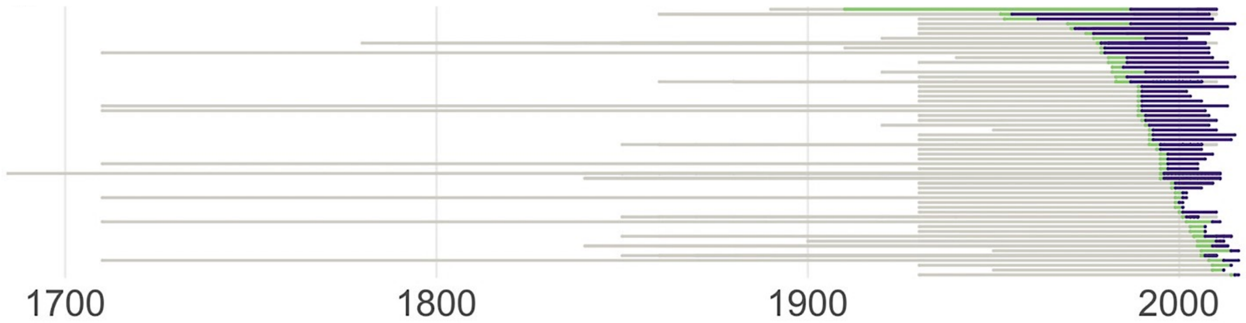


**Fig. S7. Historic peaks in forest cover change often occur decades to centuries before biodiversity monitoring starts (78% of time series).** The detected forest cover change constituted declines in forest cover, calculated based on the LUH database (*22*). Each line represents a single study from the terrestrial data within the BioTIME Database (*26*), which represented 190 studies. Note that the studies were rarefied into equally sized cells (~96 $km^2$), resulting in 2,339 time series and peaks in forest cover change were calculated on the time series level for analysis, but because of the large number of cells (> 2,000), here we visualize study-level data. Green and purple lines show the duration of biodiversity monitoring, with the break between green and purple indicating the largest forest cover change event across the duration of each time series. For our analyses, we compared biodiversity change across equal durations before and after recent peak forest cover change. E.g., if a time series included seven years of biodiversity data before the peak in recent forest cover change and 10 years of data after that peak, we included only the first seven years of biodiversity data after the forest cover change peak (making for a total of 14 years of data included in the analysis for this example time series).

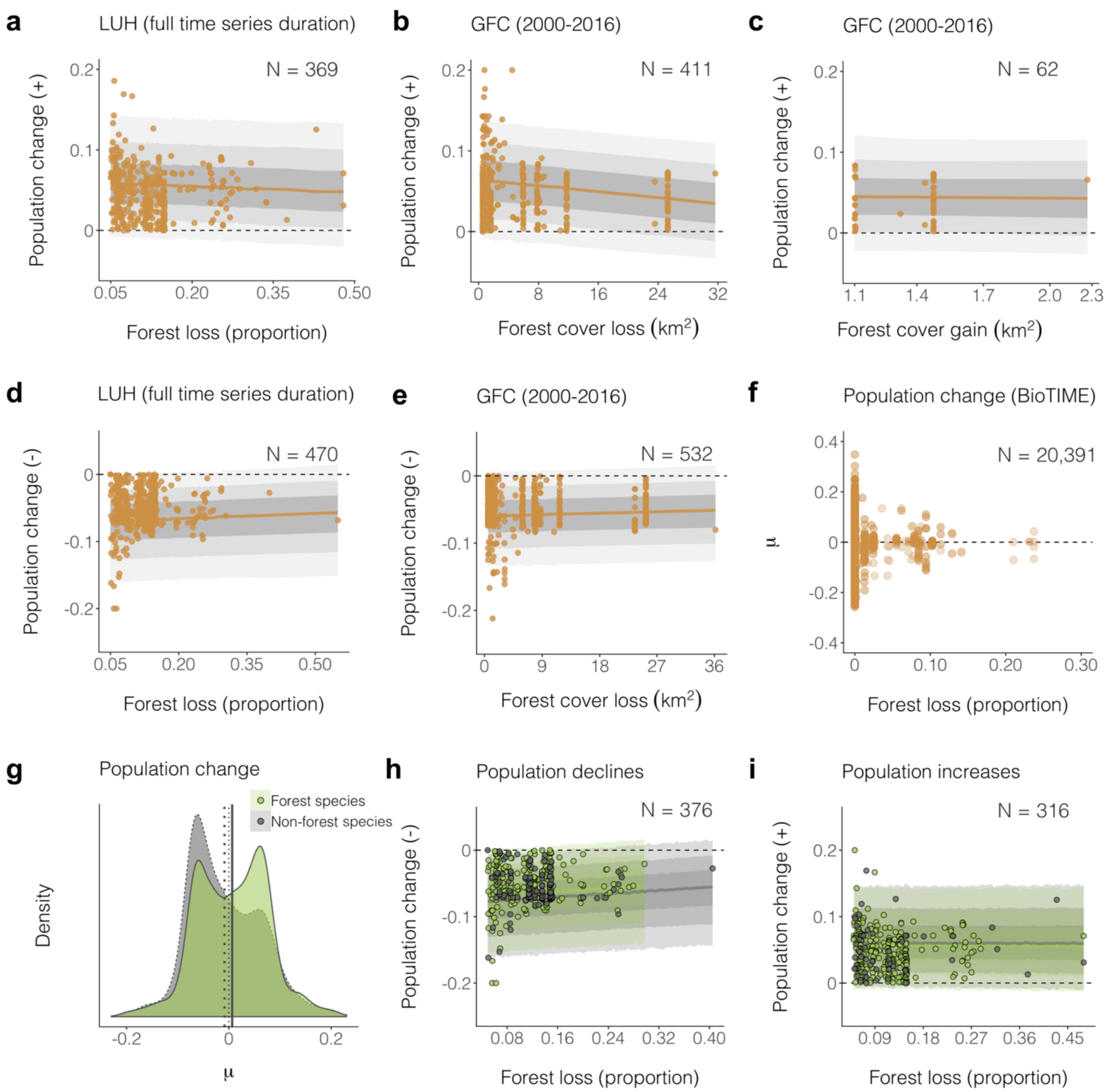


**Fig. S8. Model visualizations for forest cover change and population change.** Among time series, there were no directional trends between negative population change and forest cover change, quantified from both the GFC (*25*) and LUH (*22*) databases. Sample size was too low for the convergence of a model testing negative population change versus forest cover gain. **f**, Similar lack of directional trends between population change and forest loss were apparent when using population change, calculated based on the terrestrial vertebrate and invertebrate species population time series within the BioTIME Database. **g**, Non-forest specialist species experienced more population declines than forest species, where we found both declines and increases over time. **h**, **i**, Whether populations came from forest or non-forest species did not influence the relationships between forest loss and population declines and increases. Species were classified as forest or non-forest based on the habitat category from their IUCN Red List Assessments. Forest loss was calculated using the LUH database as proportions bounded between zero and one across the same duration as that of each individual time series. The GFC database is available for 2000 – 2016, so for the GFC models, we only used the time series records from 2000 – 2016. Only time series which had experienced over 0.5 $km^2$ forest cover change for the GFC database and 0.05 (equivalent to 5%) forest loss for the LUH database were included in the models. Population change was calculated using state-space models. Grey shades indicate the 95%, 80% and 50% credible intervals. See Table S1 for model outputs.

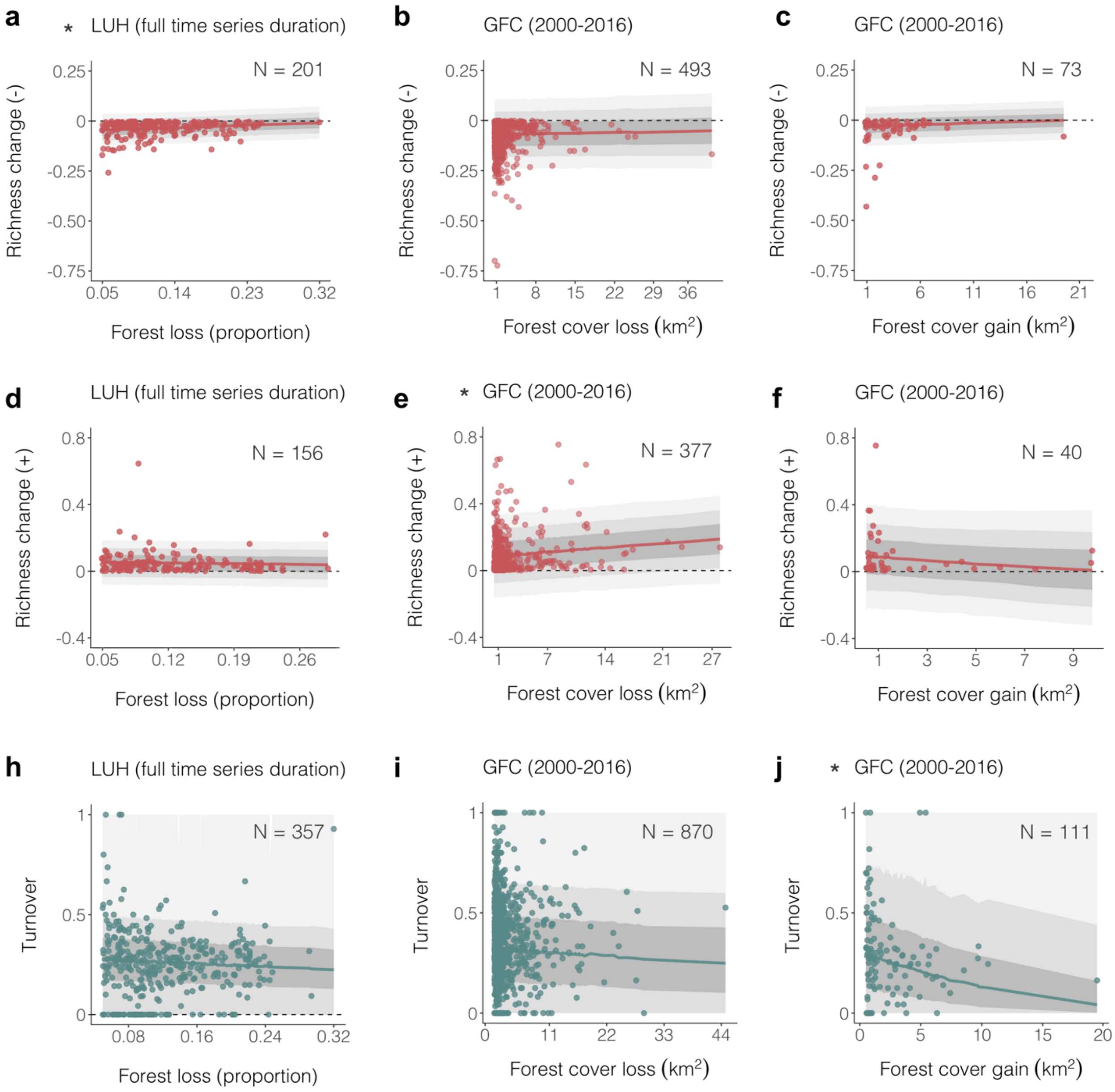


**Fig. S9. Model visualizations for forest cover change and biodiversity change (species richness and turnover).** Among time series, greater forest loss corresponded with lower species richness loss, though note that the effect size was small (slope = 0.01, CI = 0.01 to 0.01) and increases in forest cover were related to lower turnover (slope = -0.26, CI = -0.47 to -0.07). **d**, **e**, **f**, There were no directional trends between species richness losses and forest cover gain, and between turnover and forest cover loss. Larger forest cover loss corresponded with larger species richness gains (slope = 0.10, CI = 0.02 to 0.06, but note that this signal was stronger for time series with shorter durations, see Table S1 for outputs of models using time series with two or more survey points, and using time series with five or more survey points). Asterisks indicate relationships where the 95% credible intervals for the slope did not overlap zero (**a**, **e**, **j**). Forest loss was calculated using the LUH database[2] as proportions bounded between zero and one across the same duration as that of each individual time series. The GFC database is available for 2000 – 2016, so for the GFC models, we only used the time series records from 2000 – 2016. Only time series which had experienced over 0.5 $km^2$ forest cover change for the GFC database and 0.05 (equivalent to 5%) forest loss for the LUH database were included in the models. Richness change was calculated using a mixed effects model with a Poisson error distribution. Turnover refers to changes in species composition due to species replacement in the final year of the time series relative to the start of the time series. Model fits on **a**-**f** are mixed effects models with a biome random effect and a Gaussian error distribution. Model fits in **h**-**j** are zero-one-inflated beta models with a logit link function. Grey shades indicate the 95%, 80% and 50% credible intervals. See Table S1 for model outputs.

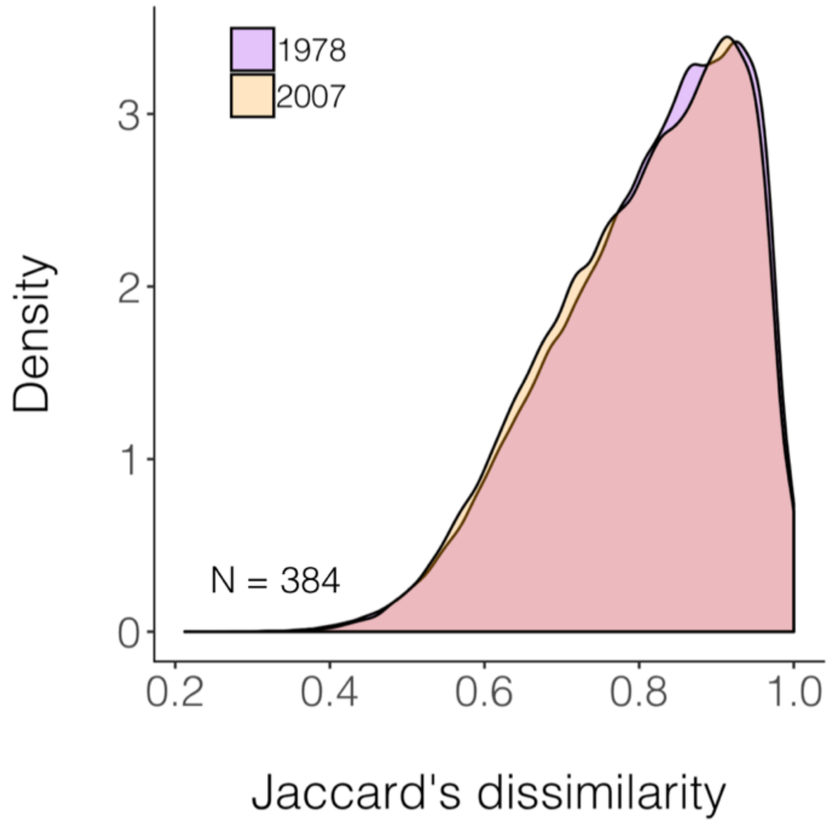


**Fig. S10. The compositional dissimilarity across sites within 384 assemblages did not decrease in 2007 relative to 1978, suggesting a lack of biotic homogenization across space among these monitored sites.** We used Jaccard’s dissimilarity index, for which zero indicates that the assemblages are exactly the same, and one indicates a completely different species composition.

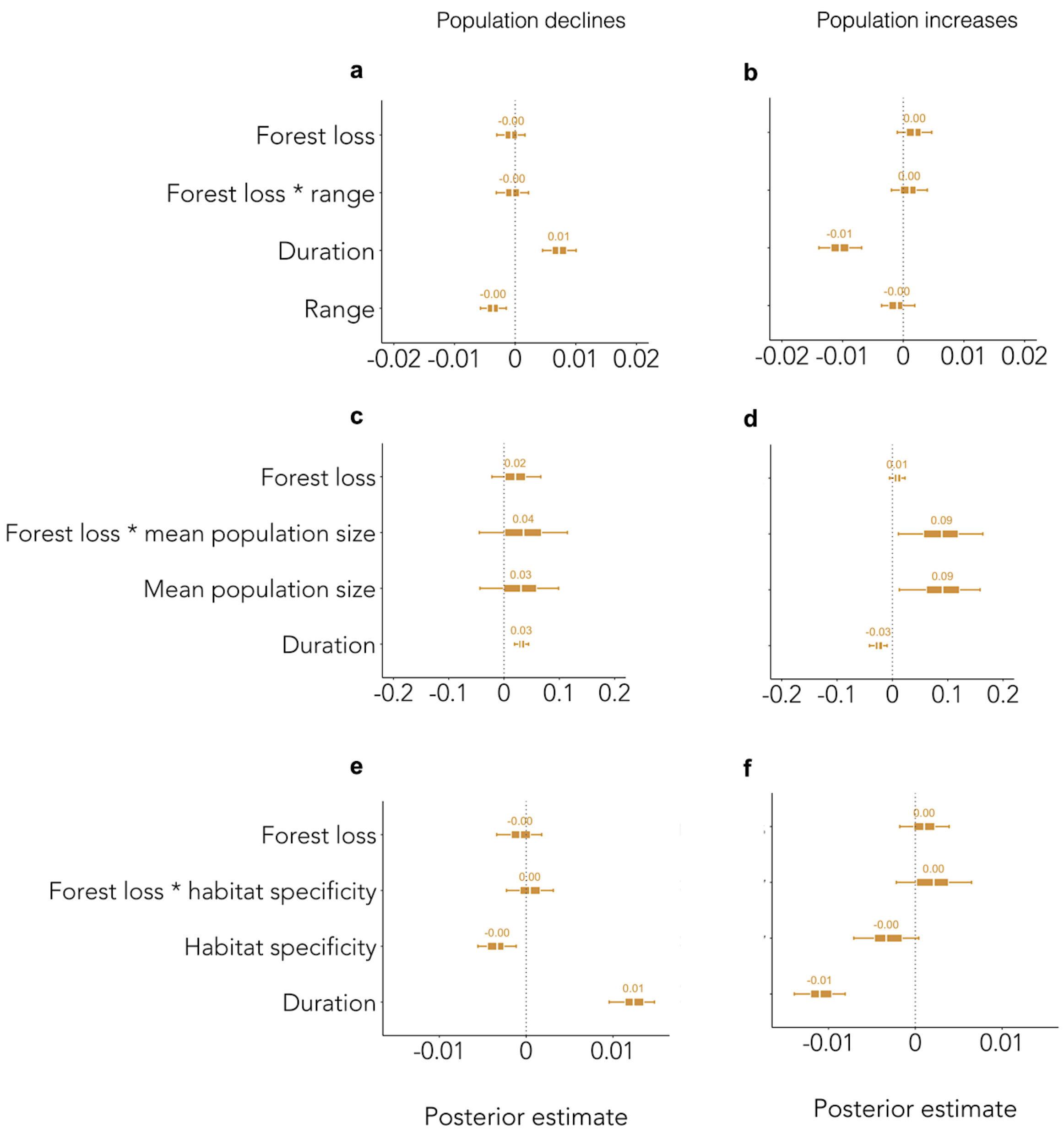


**Fig. S11. Species' geographic range, mean population size and habitat specificity, often used as proxies for rarity, were poor predictors of population responses to forest loss over time.** We tested if rare species (based on species' geographic range, mean population size and habitat specificity) were more likely to be negatively influenced by forest loss by including a forest loss * rarity metric interaction term in our models. For full details on methods to determine rarity, see (*2*). We found that regardless of whether species were rare or common, they experienced the full spectrum of forest loss effects (see Table S1 for full model outputs).

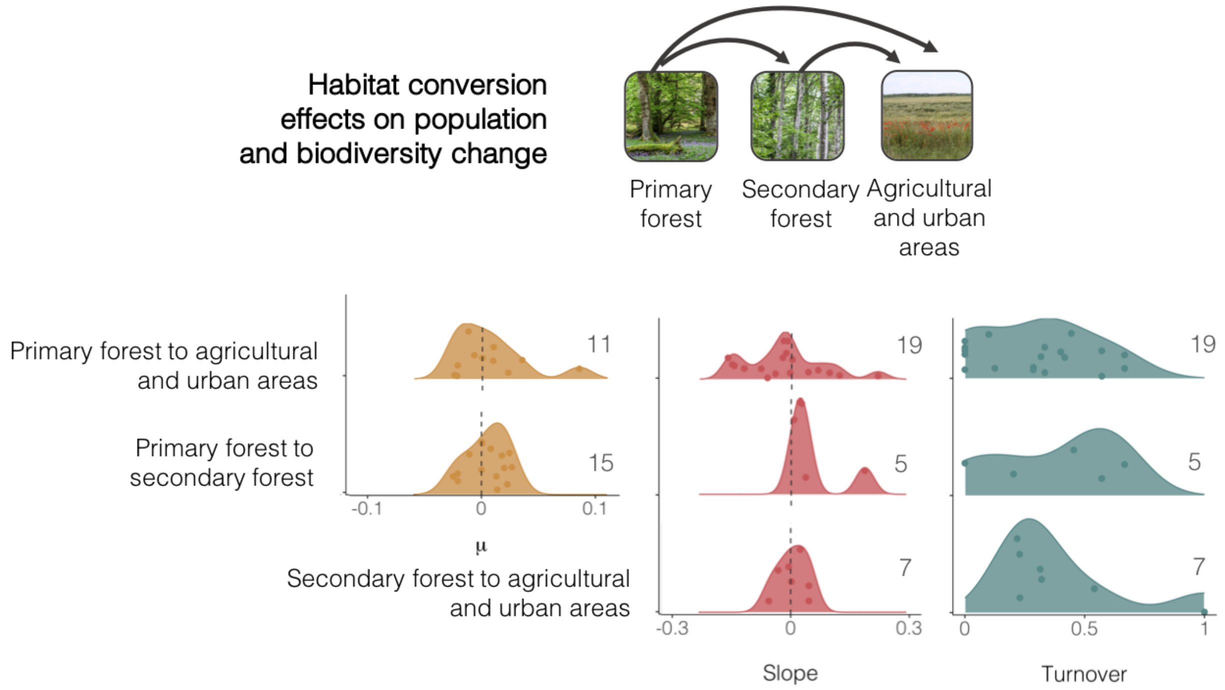


**Fig S12. Population and biodiversity change following transitions in dominant habitat type include instances declines, increases and no net changes across sites.** Distributions show *μ* values  for population change, posterior means (slopes) for richness change and Jaccard's dissimilarity for turnover under different habitat conversions. Line thickness corresponds with magnitude of detected change. Forest loss led to habitat conversion in 304 of 5795 (approximately 5%) of monitored population and biodiversity time series. There was only one instance of a population time series experiencing habitat conversion with secondary forest as the starting dominant land cover, thus no distribution is plotted for that category. The y-axis refers to the probability density function for the kernel density estimation per unit on the x-axis, and the distributions are relative to one another. Numbers in plots indicate number of time series for each category. Small sample sizes of an average 10 time series per transition types of interest for this analysis precluded statistical analysis and inferences on the effects of habitat transitions were drawn by visually inspecting the density distributions. See Figs. 7-8 and Table S1 for models of forest cover gain and loss (GFC database, 2000-2016) and forest cover loss (LUH database, across the time series) and population and biodiversity change. See Fig. S13 for distributions of population and biodiversity change following habitat transitions detected by the MODIS Land Cover Database (*24*).

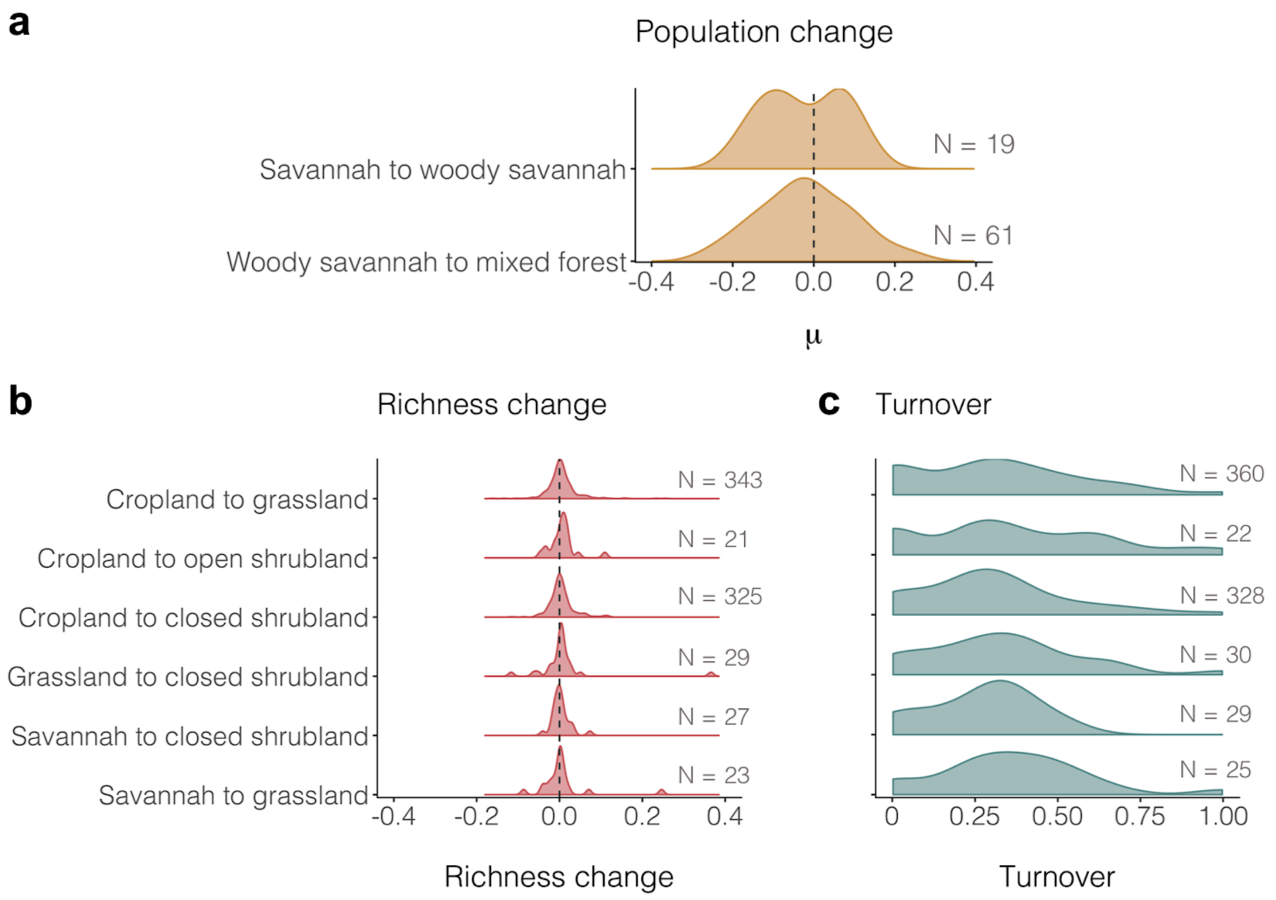


**Fig. S13. Population and biodiversity change following land-use transitions as detected by the MODIS Landcover Database** (*24*)**. a**, Population change, **b**, richness change and **c**, turnover were calculated for the period between 2001 to 2013 to match the time frame of the MODIS Landcover Database. Numbers indicate number of time series in each category. Land cover change was calculated for standardized cells of 96 km$^2$. Small and uneven sample sizes precluded statistical analysis. Visual inspection of the distributions of population and biodiversity change does not suggest directional patterns for any specific land-use transition types.

610

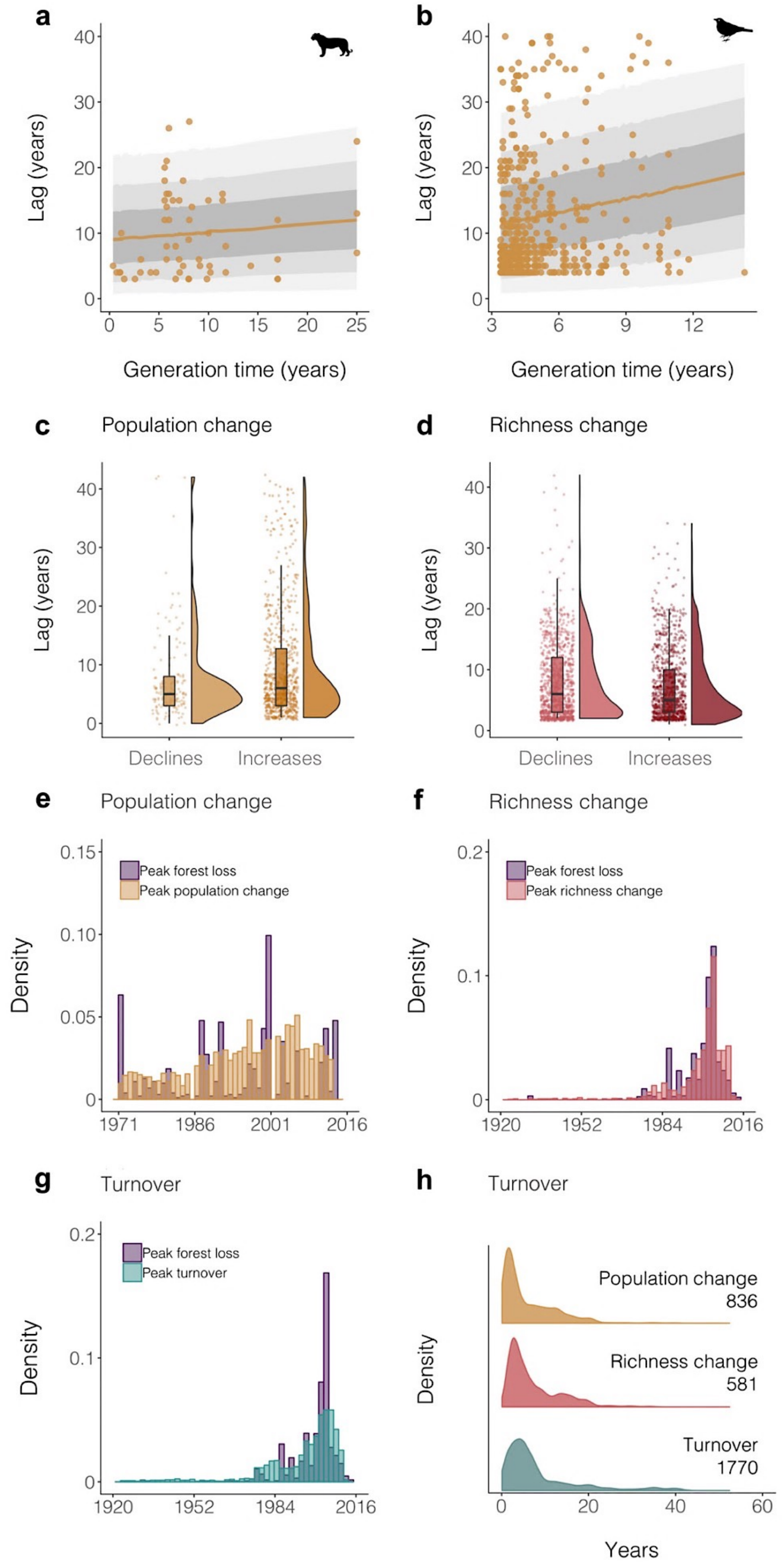
a
Lag (years)
Generation time (years)
b
Lag (years)
Generation time (years)
c
Population change
Lag (years)
Declines
Increases
d
Richness change
Lag (years)
Declines
Increases
e
Population change
Density
Peak forest loss
Peak population change
f
Richness change
Density
Peak forest loss
Peak richness change
g
Turnover
Density
Peak forest loss
Peak turnover
h
Turnover
Density
Population change
836
Richness change
581
Turnover
1770
Years

**Fig. S14. Lags in population and biodiversity change following peak forest loss occur frequently.** Temporal lags in mammal and bird population change following peak forest cover change increased with longer species' generation times (**a**-**b**), and were similar across population declines and increases, and species richness losses and gains. In general, the years in which peak forest loss occurred across sites coincided with the most population and biodiversity change (**e**-**g**). Plot h shows the distribution of lag values across the three metrics we studied – population change, richness change and turnover. The y-axis in plots e-h refers to the probability density function for the kernel density estimation per unit on the x-axis, and the distributions are relative to one another. Peak forest cover change refers to the timing of the largest forest cover change event across the duration of each time series. We categorized lags as time periods of three years (dashed horizontal line) or more between peak forest loss during the monitoring for each time series, and peak population and biodiversity change (Fig. 2B). Mammal generation times were extracted from the Pacifici *et al.* 2013 Database (*8*, N = 77). Bird generation times were extracted from the BirdLife Database (9). We found a positive relationship between temporal lag in population change and bird and mammal generation time (for birds: slope = 0.84, CI = 0.41 to 1.29; for mammals: slope = 0.31, CI = 0.04 to 0.58, grey shades indicate the 50%, 80% and 95% credible intervals for the model predictions and points represent raw data). Boxplots in **b** and **c** show the mean, with the whiskers extending to the mean ± SD. See Table S1 for full model outputs.

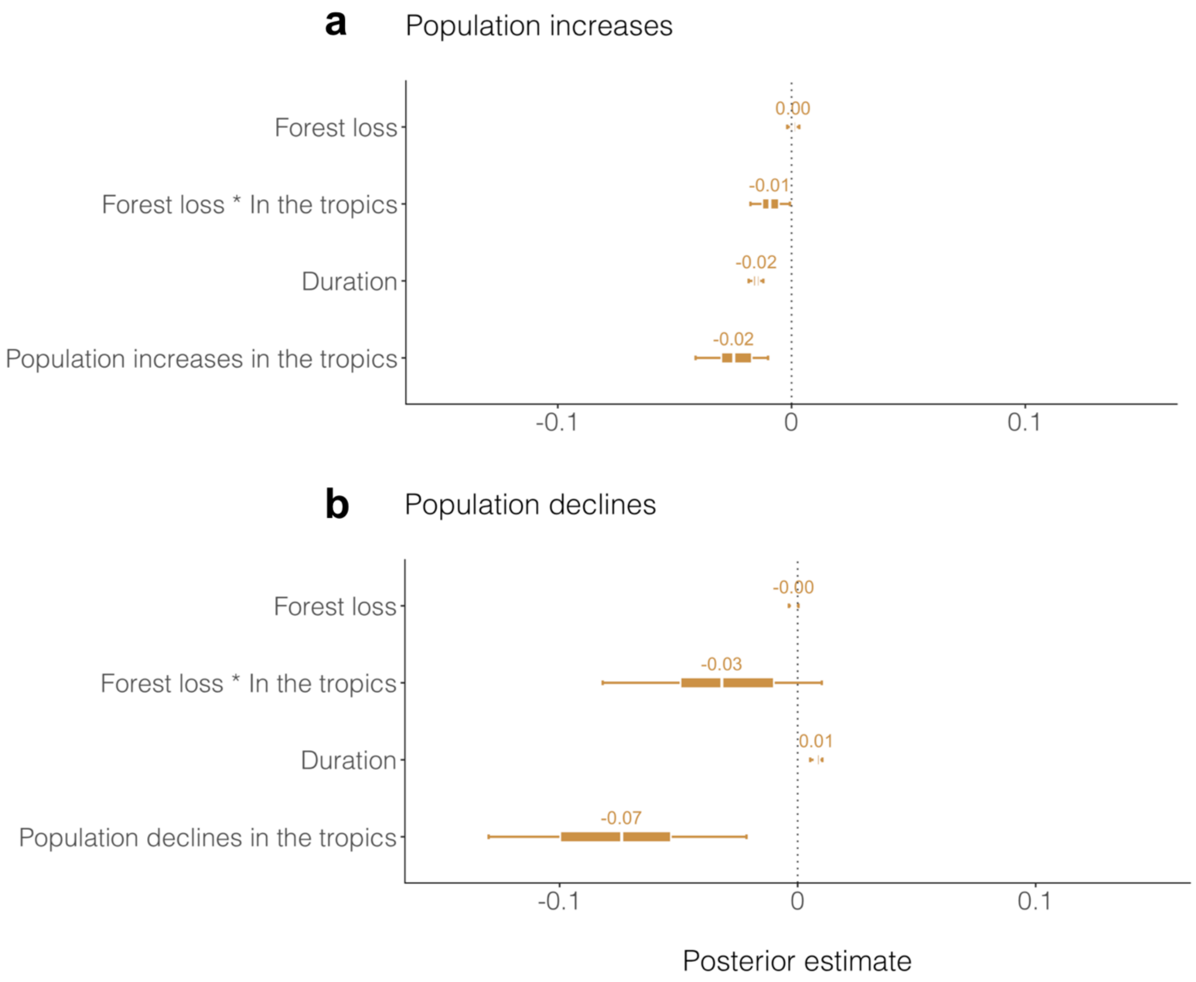


**Fig. S15. The effects of forest loss on population change were more negative in the tropics compared to the rest of the planet.** Graph shows effect sizes and 95% credible intervals from models which included a binary categorical variable (in the tropics or not). We categorized time series based on whether or not they fall within the Tropics of Capricorn and Cancer. See Table S1 for full model outputs.

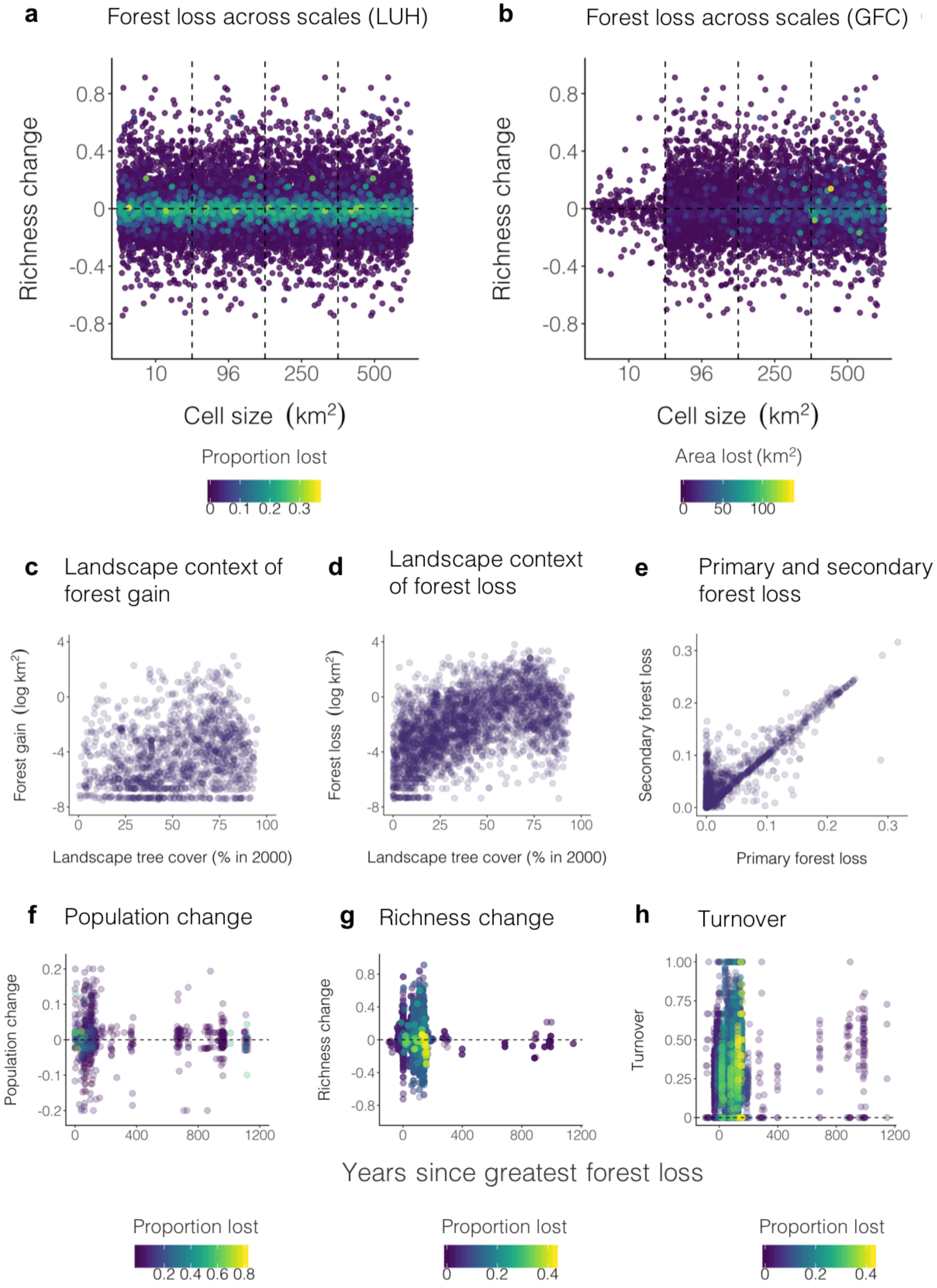

a
Forest loss across scales (LUH)
Richness change
Cell size (km²)
Proportion lost
b
Forest loss across scales (GFC)
Area lost (km²)
c
Landscape context of forest gain
Forest gain (log km²)
Landscape tree cover (% in 2000)
d
Landscape context of forest loss
Forest loss (log km²)
e
Primary and secondary forest loss
Secondary forest loss
Primary forest loss
f
Population change
g
Richness change
h
Turnover
Years since greatest forest loss


636

**Fig. S16. Sensitivity analyses. a,** Proportion forest cover lost (based on LUH Database (*22*) and **b**, amount of forest cover loss (based on GFC Database (*25*) across different cell sizes. The detected forest loss increased with larger cell sizes, but regardless of the cell size, we found both positive and negative richness trends associated with forest loss. **c** and **d,** Landscape context (500 $km^2$ cell size) of forest cover change in 2000-2016, as detected by the GFC Database. **e**, Secondary forest loss scaled positively with primary forest loss. Forest loss was measured as a proportion between zero and one. **f**, **g** and **h**, Time since all-time greatest loss in forest cover (based on LUH Database) was a poor predictor of recent population and biodiversity change. Colors on **a**-**b** and **f**-**h** indicate the magnitude of forest cover change.

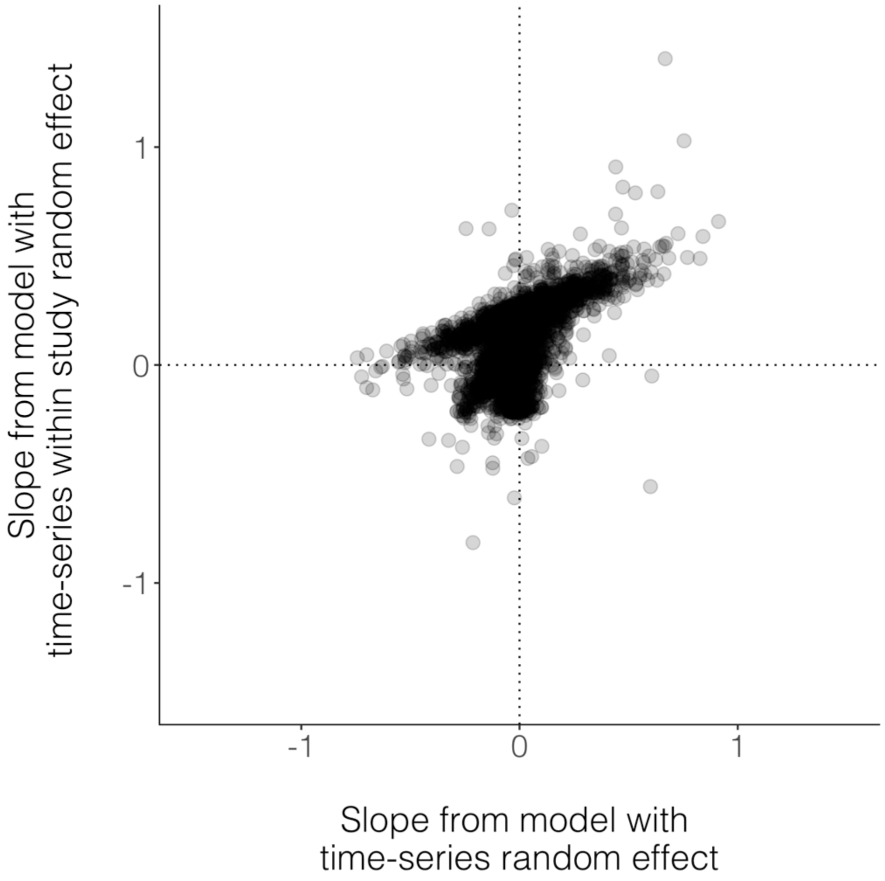


**Fig. S17. Estimating slopes of species richness change over time was not sensitive to the inclusion of a nested random effect (time series id within study).** Because some of the biodiversity studies we included in our analyses covered very large areas, we split those studies into multiple standardized cells of around 96 km2 (see methods for further details). In 28% of cases, there were more than one time series from the same study. In our analyses, we used slopes of richness change over time derived from a model including a time series id random intercept and random slope because this model converged better relative to a model with a nested random effect (time series id within study).

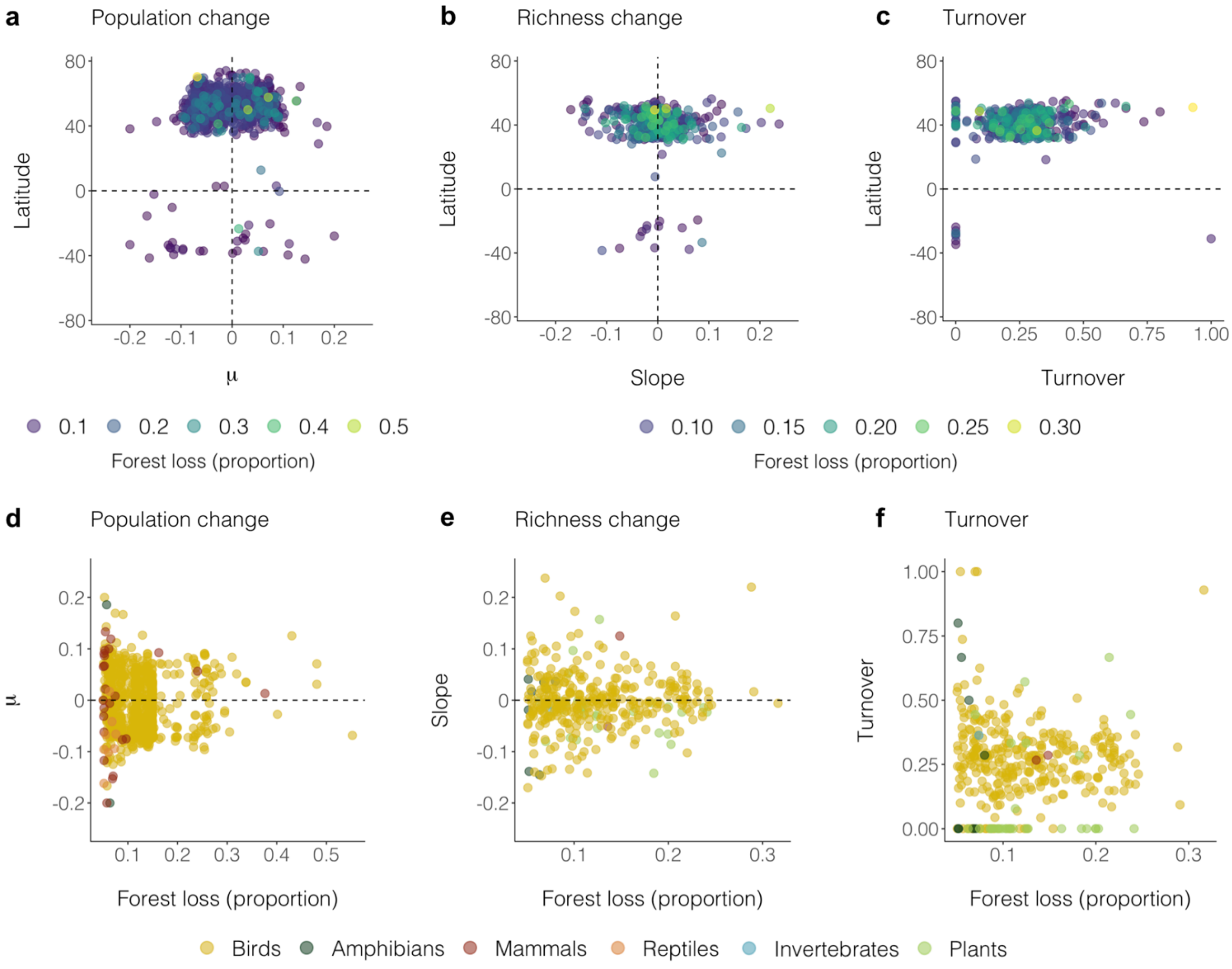


**Fig. S18. Lack of latitudinal and taxonomic patterning in the relationships between forest loss, population and biodiversity change.** Plots show population change, richness change and turnover over time across latitude (**a**, **b**, and **c**) and across different taxa (**d**, **e**, and **f**) for time series that experienced at least 5% loss in forest cover across their duration. Forest loss was calculated using the LUH database as proportions bounded between zero and one across the same duration as that of each individual time series. The geographic and taxonomic gaps in the Living Planet and BioTIME databases reflect current differences in survey effort and public availability of data.

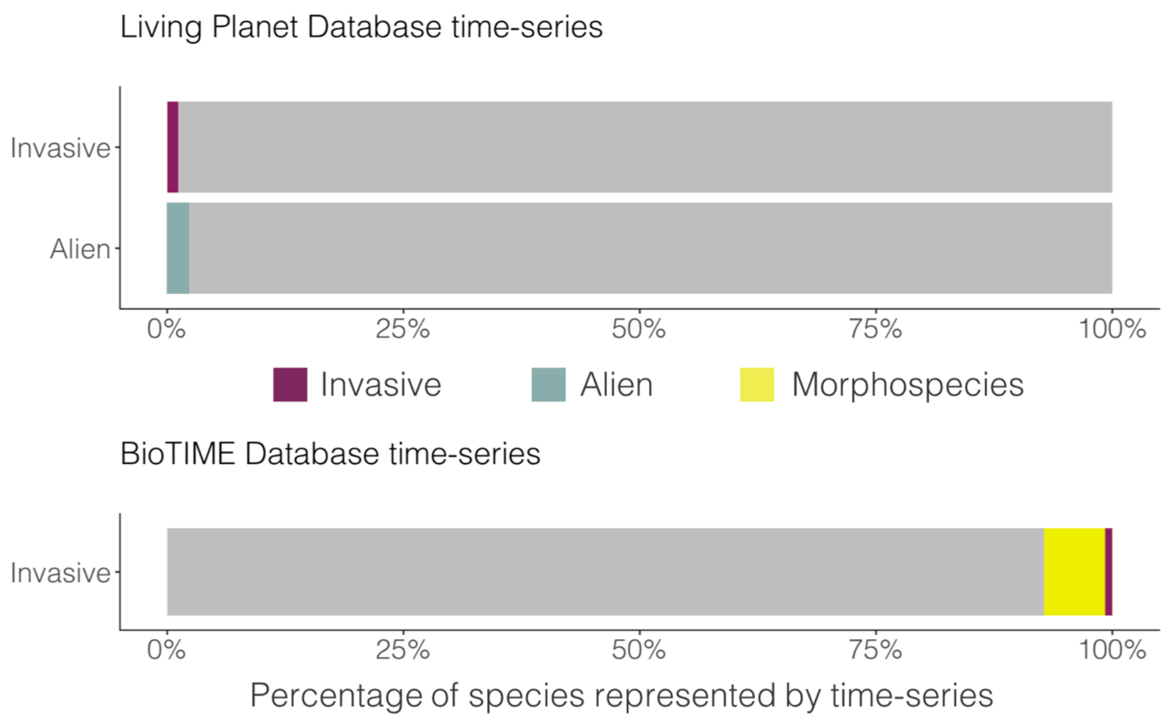


**Fig. S19. The majority of the species represented by the databases we analyzed are not classified as alien or invasive (grey bars).** All species in the Living Planet Database were identified to the species level, whereas in the BioTIME database, 7% of species were recorded as morphospecies. Alien and invasive categorizations were available from the Living Planet Database. To classify the species from the BioTIME database, we used the Global Invasive Species Database (*57*).

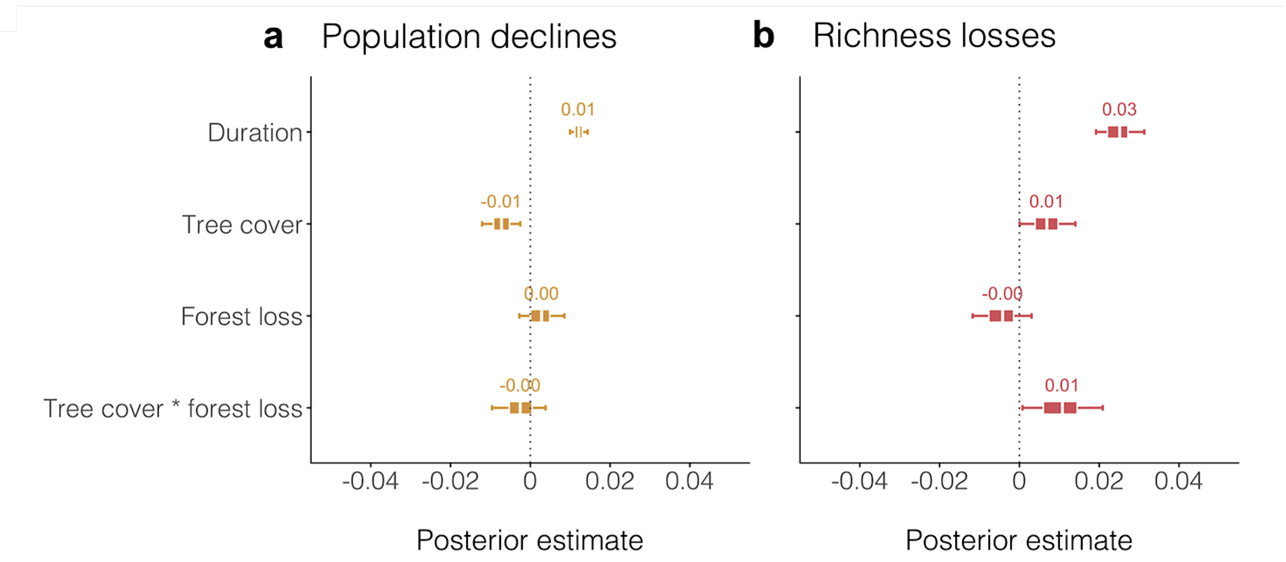


**Fig. S20. The effects of forest loss on species richness over time were less negative in places with higher tree cover.** We did not detect a significant interaction between tree cover and forest loss on population declines. We calculated tree cover using the GFC Database. See Table S1 for full model outputs.

**Table S1. Model outputs for all analyses.** See methods for justification of models. Scaled variables were mean centered on zero. Term names starting with “b” refer to fixed effects and term names starting with “r” refer to random intercepts. Sigma indicates the residual variance. For turnover models, “zoi” refers to the probability of being a zero or a one, “coi” refers to the conditional probability of being a one (given an observation is a zero or a one), and “phi” is the precision parameter of zero-one inflated beta distribution. “Scaled” indicates variables which were centered with a mean of zero. Numbers (N) refer to the sample size for each model and are presented as number of time series out of the overall number of population (4,228) and biodiversity (2,339) time series we analyzed. The sample sizes for each model were determined based on the aim of the model (e.g., testing specifically time series where populations declined, and the sites experienced more than 5% forest loss). See methods for details on analyses.

| Model | Term | Estimate | Std. error | Lower 95% CI | Upper 95% CI |
|---|---|---|---|---|---|
| **Population declines before/after forest loss** (N = 467 / 4,228) | b.intercept | -0.040 | 0.011 | -0.061 | -0.025 |
| | b.after.forest.loss | -0.037 | 0.002 | -0.041 | -0.033 |
| | b.duration.scaled | 0.010 | 0.001 | 0.008 | 0.012 |
| | sigma | 0.033 | 0.001 | 0.032 | 0.035 |
| | biome.boreal.forests.taiga | 0.016 | 0.011 | 0 | 0.036 |
| | biome.flooded.grasslands.and.savannas | -0.036 | 0.033 | -0.083 | 0.006 |
| | biome.mediterranean.forests.woodlands.and.scrub | 0.013 | 0.021 | -0.017 | 0.056 |
| | biome.temperate.broadleaf.and.mixed.forests | 0.008 | 0.011 | -0.009 | 0.027 |
| | biome.temperate.coniferous.forests | 0.009 | 0.011 | -0.008 | 0.029 |
| | biome.temperate.grasslands.savannas.and.shrublands | 0.008 | 0.012 | -0.011 | 0.031 |
| | biome.tropical.and.subtropical.coniferous.forests | -0.004 | 0.016 | -0.032 | 0.027 |
| | biome.tropical.and.subtropical.grasslands.savannas.and.shrublands | -0.012 | 0.014 | -0.038 | 0.009 |
| | biome.tropical.and.subtropical.moist.broadleaf.forests | -0.026 | 0.023 | -0.060 | 0.006 |
| | biome.tundra | 0.015 | 0.013 | -0.003 | 0.036 |
| **Population increases before/after forest loss** (N = 343 / 4,228) | b.intercept | 0.034 | 0.004 | 0.028 | 0.041 |
| | b.after.forest.loss | 0.022 | 0.003 | 0.017 | 0.027 |
| | b.duration.scaled | -0.010 | 0.002 | -0.013 | -0.008 |
| | sigma | 0.038 | 0.001 | 0.036 | 0.040 |
| | biome.boreal.forests.taiga | -0.002 | 0.003 | -0.010 | 0.002 |
| | biome.flooded.grasslands.and.savannas | 0 | 0.004 | -0.008 | 0.012 |
| | biome.mediterranean.forests.woodlands.and.scrub | -0.001 | 0.004 | -0.013 | 0.008 |
| | biome.temperate.broadleaf.and.mixed.forests | 0 | 0.003 | -0.006 | 0.006 |
| | biome.temperate.coniferous.forests | -0.001 | 0.003 | -0.010 | 0.005 |

| | | | | | |
|---|---|---|---|---|---|
| | biome.temperate.grasslands.savannas.and.shrublands | 0.001 | 0.004 | -0.005 | 0.012 |
| | biome.tropical.and.subtropical.grasslands.savannas.and.shrublands | 0.001 | 0.004 | -0.007 | 0.013 |
| | biome.tundra | 0 | 0.003 | -0.008 | 0.008 |
| **Population change before/during/after peak deforestation** (N = 1,941 / 4,228) | b.intercept | 0.002 | 0.006 | -0.010 | 0.011 |
| | b.periodduring | -0.007 | 0.003 | -0.012 | -0.001 |
| | b.after.forest.loss | -0.009 | 0.004 | -0.015 | -0.003 |
| | b.duration.scaled | -0.006 | 0.002 | -0.009 | -0.003 |
| | sigma | 0.057 | 0.001 | 0.056 | 0.059 |
| | biome.boreal.forests.taiga | 0.002 | 0.007 | -0.009 | 0.014 |
| | biome.deserts.and.xeric.shrublands | -0.030 | 0.012 | -0.049 | -0.012 |
| | biome.flooded.grasslands.and.savannas | -0.011 | 0.017 | -0.040 | 0.016 |
| | biome.mediterranean.forests.woodlands.and.scrub | 0.010 | 0.008 | -0.003 | 0.023 |
| | biome.montane.grasslands.and.shrublands | 0.013 | 0.016 | -0.012 | 0.039 |
| | biome.temperate.broadleaf.and.mixed.forests | 0.009 | 0.007 | -0.002 | 0.020 |
| | biome.temperate.coniferous.forests | 0.001 | 0.007 | -0.011 | 0.013 |
| | biome.temperate.grasslands.savannas.and.shrublands | -0.004 | 0.008 | -0.017 | 0.009 |
| | biome.tropical.and.subtropical.coniferous.forests | 0.003 | 0.016 | -0.023 | 0.033 |
| | biome.tropical.and.subtropical.dry.broadleaf.forests | -0.022 | 0.015 | -0.048 | 0.001 |
| | biome.tropical.and.subtropical.grasslands.savannas.and.shrublands | -0.003 | 0.008 | -0.016 | 0.010 |
| | biome.tropical.and.subtropical.moist.broadleaf.forests | 0.018 | 0.009 | 0.004 | 0.032 |
| | biome.tundra | 0.011 | 0.008 | -0.002 | 0.025 |
| **Richness losses before/after peak deforestation** (N = 188 / 2,339) | b.intercept | -0.100 | 0.033 | -0.163 | -0.043 |
| | b.after.forest.loss | -0.018 | 0.011 | -0.036 | -0.001 |
| | b.duration.scaled | 0.035 | 0.006 | 0.026 | 0.044 |
| | sigma | 0.097 | 0.004 | 0.091 | 0.103 |
| | biome.boreal.forests.taiga | 0.032 | 0.042 | -0.039 | 0.105 |
| | biome.deserts.and.xeric.shrublands | 0.007 | 0.039 | -0.065 | 0.071 |
| | biome.mediterranean.forests.woodlands.and.scrub | -0.026 | 0.046 | -0.105 | 0.048 |
| | biome.montane.grasslands.and.shrublands | -0.146 | 0.076 | -0.268 | -0.031 |
| | biome.temperate.broadleaf.and.mixed.forests | -0.035 | 0.033 | -0.091 | 0.029 |
| | biome.temperate.conifer.forests | 0.053 | 0.037 | -0.011 | 0.116 |
| | biome.temperate.grasslands.savannas.and.shrublands | 0.056 | 0.035 | 0 | 0.124 |
| | biome.tropical.and.subtropical.grasslands.savannas.and.shrublands | 0.048 | 0.059 | -0.041 | 0.154 |
| **Richness gains before/after peak deforestation** (N = 266 / 2,339) | b.intercept | 0.030 | 0.005 | 0.020 | 0.039 |
| | b.after.forest.loss | -0.002 | 0.001 | -0.004 | 0 |
| | b.duration.scaled | 0.002 | 0.001 | 0 | 0.003 |
| | sigma | 0.016 | 0.001 | 0.015 | 0.017 |
| | biome.boreal.forests.taiga | 0.004 | 0.007 | -0.008 | 0.015 |
| | biome.deserts.and.xeric.shrublands | 0.010 | 0.008 | -0.002 | 0.024 |
| | biome.mediterranean.forests.woodlands.and.scrub | 0.004 | 0.009 | -0.009 | 0.020 |
| | biome.temperate.broadleaf.and.mixed.forests | 0 | 0.005 | -0.010 | 0.010 |
| | biome.temperate.conifer.forests | -0.005 | 0.006 | -0.016 | 0.004 |
| | biome.temperate.grasslands.savannas.and.shrublands | -0.013 | 0.006 | -0.024 | -0.003 |
| **Turnover before/after peak deforestation** | b.intercept | -1.009 | 0.095 | -1.219 | -0.831 |
| | b.zoi.intercept | -1.266 | 0.076 | -1.392 | -1.146 |

| | | | | | |
|---|---|---|---|---|---|
| (N = 389 / 2,339) | b.coi.intercept | -2.553 | 0.235 | -2.937 | -2.190 |
| | b.after.forest.loss | -0.036 | 0.043 | -0.105 | 0.031 |
| | b.duration.scaled | -0.121 | 0.044 | -0.195 | -0.052 |
| | phi | 13.448 | 0.684 | 12.352 | 14.551 |
| | biome.boreal.forests.taiga | 0.211 | 0.174 | -0.026 | 0.493 |
| | biome.deserts.and.xeric.shrublands | 0.210 | 0.171 | -0.023 | 0.500 |
| | biome.mediterranean.forests.woodlands.and.scrub | -0.088 | 0.168 | -0.450 | 0.164 |
| | biome.montane.grasslands.and.shrublands | -0.001 | 0.204 | -0.467 | 0.486 |
| | biome.temperate.broadleaf.and.mixed.forests | -0.042 | 0.094 | -0.214 | 0.165 |
| | biome.temperate.conifer.forests | 0.010 | 0.104 | -0.165 | 0.243 |
| | biome.temperate.grasslands.savannas.and.shrublands | -0.069 | 0.104 | -0.273 | 0.133 |
| | biome.tropical.and.subtropical.dry.broadleaf.forests | -0.142 | 0.225 | -0.671 | 0.160 |
| | biome.tropical.and.subtropical.grasslands.savannas.and.shrublands | -0.050 | 0.172 | -0.434 | 0.268 |
| | biome.tropical.and.subtropical.moist.broadleaf.forests | -0.005 | 0.194 | -0.463 | 0.432 |
| **Population change and magnitude of peak forest loss** (N = 618 / 4,228) | b.intercept | 0.028 | 0.006 | 0.017 | 0.039 |
| | b.max.loss.scaled | -0.002 | 0.002 | -0.005 | 0.001 |
| | b.duration.scaled | -0.021 | 0.002 | -0.024 | -0.018 |
| | sigma | 0.043 | 0.001 | 0.041 | 0.045 |
| | biome.boreal.forests.taiga | -0.001 | 0.006 | -0.013 | 0.010 |
| | biome.flooded.grasslands.and.savannas | 0.009 | 0.014 | -0.010 | 0.043 |
| | biome.mediterranean.forests.woodlands.and.scrub | -0.001 | 0.010 | -0.024 | 0.022 |
| | biome.temperate.broadleaf.and.mixed.forests | 0.008 | 0.007 | -0.002 | 0.021 |
| | biome.temperate.coniferous.forests | 0.001 | 0.007 | -0.011 | 0.014 |
| | biome.temperate.grasslands.savannas.and.shrublands | 0 | 0.008 | -0.016 | 0.015 |
| | biome.tropical.and.subtropical.coniferous.forests | 0.001 | 0.011 | -0.023 | 0.027 |
| | biome.tropical.and.subtropical.grasslands.savannas.and.shrublands | -0.017 | 0.018 | -0.049 | 0.005 |
| | biome.tropical.and.subtropical.moist.broadleaf.forests | 0 | 0.011 | -0.025 | 0.024 |
| | biome.tundra | -0.002 | 0.008 | -0.017 | 0.011 |
| **Richness change and magnitude of peak forest loss** (N = 386 / 2,339) | b.intercept | 0.028 | 0.005 | 0.019 | 0.040 |
| | b.max.loss.scaled | -0.003 | 0.001 | -0.005 | -0.001 |
| | b.duration.scaled | -0.002 | 0.002 | -0.006 | 0.001 |
| | sigma | 0.027 | 0.001 | 0.025 | 0.028 |
| | biome.boreal.forests.taiga | 0.007 | 0.008 | -0.003 | 0.021 |
| | biome.deserts.and.xeric.shrublands | -0.005 | 0.007 | -0.020 | 0.006 |
| | biome.mediterranean.forests.woodlands.and.scrub | 0 | 0.006 | -0.014 | 0.012 |
| | biome.montane.grasslands.and.shrublands | 0.013 | 0.018 | -0.006 | 0.058 |
| | biome.temperate.broadleaf.and.mixed.forests | -0.001 | 0.005 | -0.013 | 0.007 |
| | biome.temperate.conifer.forests | -0.003 | 0.006 | -0.016 | 0.005 |
| | biome.temperate.grasslands.savannas.and.shrublands | -0.006 | 0.006 | -0.019 | 0.002 |
| | biome.tropical.and.subtropical.grasslands.savannas.and.shrublands | -0.002 | 0.009 | -0.027 | 0.015 |
| **Turnover and magnitude of peak forest loss** (N = 386 / 2,339) | b.intercept | 0.220 | 0.115 | 0.013 | 0.425 |
| | b.max.loss.scaled | 0.005 | 0.007 | -0.006 | 0.016 |
| | b.duration.scaled | -0.010 | 0.004 | -0.017 | -0.003 |
| | sigma | 0.091 | 0.003 | 0.087 | 0.096 |
| | biome.boreal.forests.taiga | -0.153 | 0.117 | -0.355 | 0.059 |
| | biome.deserts.and.xeric.shrublands | -0.121 | 0.118 | -0.327 | 0.087 |

| | | | | | |
|---|---|---|---|---|---|
| | biome.mediterranean.forests.woodlands.and.scrub | -0.070 | 0.118 | -0.278 | 0.138 |
| | biome.montane.grasslands.and.shrublands | 0.699 | 0.141 | 0.473 | 0.957 |
| | biome.temperate.broadleaf.and.mixed.forests | -0.125 | 0.115 | -0.333 | 0.080 |
| | biome.temperate.conifer.forests | -0.122 | 0.116 | -0.322 | 0.090 |
| | biome.temperate.grasslands.savannas.and.shrublands | -0.125 | 0.116 | -0.333 | 0.080 |
| | biome.tropical.and.subtropical.grasslands.savannas.and.shrublands | 0.004 | 0.142 | -0.223 | 0.254 |
| **Population declines and overall forest loss** (N = 470 / 4,228) | b.intercept | -0.052 | 0.003 | -0.062 | -0.047 |
| | b.forest.loss.scaled | -0.001 | 0.001 | -0.003 | 0.001 |
| | b.duration.scaled | 0.012 | 0.001 | 0.009 | 0.014 |
| | sigma | 0.026 | 0.001 | 0.025 | 0.028 |
| | biome.boreal.forests.taiga | 0.001 | 0.003 | -0.004 | 0.013 |
| | biome.deserts.and.xeric.shrublands | -0.003 | 0.007 | -0.031 | 0.007 |
| | biome.mediterranean.forests.woodlands.and.scrub | -0.001 | 0.005 | -0.018 | 0.007 |
| | biome.temperate.broadleaf.and.mixed.forests | 0 | 0.003 | -0.006 | 0.010 |
| | biome.temperate.coniferous.forests | 0.001 | 0.004 | -0.006 | 0.012 |
| | biome.temperate.grasslands.savannas.and.shrublands | 0.001 | 0.004 | -0.007 | 0.012 |
| | biome.tropical.and.subtropical.coniferous.forests | -0.001 | 0.005 | -0.021 | 0.011 |
| | biome.tropical.and.subtropical.grasslands.savannas.and.shrublands | -0.003 | 0.006 | -0.024 | 0.006 |
| | biome.tropical.and.subtropical.moist.broadleaf.forests | 0.002 | 0.005 | -0.008 | 0.022 |
| | biome.tundra | 0.006 | 0.009 | -0.003 | 0.024 |
| **Population increases and overall forest loss** (N = 369 / 4,228) | b.intercept | 0.047 | 0.003 | 0.042 | 0.054 |
| | b.forest.loss.scaled | 0 | 0.002 | -0.003 | 0.002 |
| | b.duration.scaled | -0.012 | 0.002 | -0.015 | -0.009 |
| | sigma | 0.029 | 0.001 | 0.027 | 0.031 |
| | biome.boreal.forests.taiga | 0 | 0.002 | -0.008 | 0.006 |
| | biome.mediterranean.forests.woodlands.and.scrub | 0.001 | 0.003 | -0.005 | 0.017 |
| | biome.temperate.broadleaf.and.mixed.forests | 0 | 0.002 | -0.008 | 0.004 |
| | biome.temperate.coniferous.forests | 0 | 0.002 | -0.009 | 0.006 |
| | biome.temperate.grasslands.savannas.and.shrublands | 0 | 0.002 | -0.010 | 0.005 |
| | biome.tropical.and.subtropical.grasslands.savannas.and.shrublands | -0.001 | 0.003 | -0.017 | 0.006 |
| | biome.tropical.and.subtropical.moist.broadleaf.forests | 0.001 | 0.003 | -0.006 | 0.023 |
| | biome.tundra | -0.001 | 0.003 | -0.011 | 0.005 |
| **Population declines and forest cover loss (2000-2016)** (N = 532 / 4,228) | b.intercept | -0.052 | 0.002 | -0.057 | -0.046 |
| | b.forest.loss.scaled | -0.002 | 0.001 | -0.004 | 0.001 |
| | b.duration.scaled | 0.012 | 0.001 | 0.009 | 0.014 |
| | sigma | 0.027 | 0.001 | 0.025 | 0.028 |
| | biome.boreal.forests.taiga | 0 | 0.002 | -0.006 | 0.006 |
| | biome.mediterranean.forests.woodlands.and.scrub | 0 | 0.003 | -0.007 | 0.009 |
| | biome.temperate.broadleaf.and.mixed.forests | 0 | 0.002 | -0.006 | 0.005 |
| | biome.temperate.coniferous.forests | 0.001 | 0.003 | -0.004 | 0.009 |
| | biome.temperate.grasslands.savannas.and.shrublands | 0 | 0.003 | -0.008 | 0.009 |
| | biome.tropical.and.subtropical.dry.broadleaf.forests | -0.001 | 0.003 | -0.014 | 0.008 |
| | biome.tropical.and.subtropical.grasslands.savannas.and.shrublands | 0.001 | 0.004 | -0.007 | 0.017 |
| | biome.tropical.and.subtropical.moist.broadleaf.forests | -0.003 | 0.004 | -0.017 | 0.003 |
| | b.intercept | 0.051 | 0.003 | 0.047 | 0.057 |

| | | | | | |
|---|---|---|---|---|---|
| **Population increases and forest cover loss (2000-2016)** (N = 411 / 4,228) | b.forest.loss.scaled | -0.001 | 0.002 | -0.004 | 0.002 |
| | b.duration.scaled | -0.015 | 0.002 | -0.018 | -0.012 |
| | sigma | 0.032 | 0.001 | 0.030 | 0.034 |
| | biome.boreal.forests.taiga | -0.002 | 0.003 | -0.009 | 0.003 |
| | biome.deserts.and.xeric.shrublands | 0 | 0.003 | -0.009 | 0.008 |
| | biome.mediterranean.forests.woodlands.and.scrub | 0 | 0.003 | -0.009 | 0.006 |
| | biome.temperate.broadleaf.and.mixed.forests | 0.001 | 0.002 | -0.004 | 0.007 |
| | biome.temperate.coniferous.forests | 0 | 0.003 | -0.005 | 0.007 |
| | biome.temperate.grasslands.savannas.and.shrublands | 0.001 | 0.003 | -0.006 | 0.009 |
| | biome.tropical.and.subtropical.grasslands.savannas.and.shrublands | 0 | 0.003 | -0.006 | 0.011 |
| | biome.tropical.and.subtropical.moist.broadleaf.forests | 0 | 0.003 | -0.008 | 0.008 |
| | biome.tundra | 0 | 0.003 | -0.011 | 0.007 |
| **Population increases and forest cover gain (2000-2016)** (N = 54 / 4,228) | b.intercept | 0.049 | 0.048 | -0.064 | 0.266 |
| | b.forest.gain.scaled | -0.003 | 0.005 | -0.012 | 0.005 |
| | b.duration.scaled | 0.002 | 0.012 | -0.014 | 0.028 |
| | sigma | 0.027 | 0.003 | 0.023 | 0.032 |
| | biome.temperate.broadleaf.and.mixed.forests | -0.005 | 0.049 | -0.233 | 0.103 |
| | biome.temperate.coniferous.forests | 0.019 | 0.062 | -0.109 | 0.238 |
| | biome.tundra | -0.013 | 0.045 | -0.197 | 0.109 |
| **Richness losses and overall forest loss** (N = 201 / 2,339) | b.intercept | -0.036 | 0.005 | -0.045 | -0.027 |
| | b.forest.loss.scaled | 0.006 | 0.003 | 0.001 | 0.011 |
| | b.duration.scaled | 0.003 | 0.003 | -0.002 | 0.008 |
| | sigma | 0.039 | 0.002 | 0.036 | 0.042 |
| | biome.boreal.forests.taiga | 0 | 0.005 | -0.013 | 0.016 |
| | biome.deserts.and.xeric.shrublands | 0.002 | 0.005 | -0.008 | 0.015 |
| | biome.mediterranean.forests.woodlands.and.scrub | 0 | 0.005 | -0.015 | 0.012 |
| | biome.montane.grasslands.and.shrublands | 0 | 0.005 | -0.016 | 0.013 |
| | biome.temperate.broadleaf.and.mixed.forests | 0 | 0.004 | -0.009 | 0.009 |
| | biome.temperate.conifer.forests | -0.004 | 0.006 | -0.017 | 0.004 |
| | biome.temperate.grasslands.savannas.and.shrublands | -0.001 | 0.005 | -0.013 | 0.010 |
| | biome.tropical.and.subtropical.grasslands.savannas.and.shrublands | 0.001 | 0.005 | -0.012 | 0.017 |
| | biome.tropical.and.subtropical.moist.broadleaf.forests | 0.001 | 0.006 | -0.011 | 0.018 |
| **Richness gains and overall forest loss** (N = 156 / 2,339) | b.intercept | 0.045 | 0.009 | 0.026 | 0.063 |
| | b.forest.loss.scaled | 0.001 | 0.006 | -0.009 | 0.009 |
| | b.duration.scaled | -0.015 | 0.006 | -0.024 | -0.006 |
| | sigma | 0.063 | 0.004 | 0.057 | 0.068 |
| | biome.boreal.forests.taiga | -0.002 | 0.014 | -0.043 | 0.028 |
| | biome.deserts.and.xeric.shrublands | 0.009 | 0.016 | -0.013 | 0.049 |
| | biome.mediterranean.forests.woodlands.and.scrub | -0.004 | 0.013 | -0.040 | 0.019 |
| | biome.montane.grasslands.and.shrublands | -0.003 | 0.013 | -0.040 | 0.022 |
| | biome.temperate.broadleaf.and.mixed.forests | -0.001 | 0.009 | -0.020 | 0.020 |
| | biome.temperate.conifer.forests | 0.013 | 0.017 | -0.007 | 0.042 |
| | biome.temperate.grasslands.savannas.and.shrublands | -0.003 | 0.011 | -0.026 | 0.019 |
| | biome.tropical.and.subtropical.dry.broadleaf.forests | -0.003 | 0.014 | -0.042 | 0.028 |
| **Richness losses and forest cover loss (2000-2016)** | b.intercept | -0.070 | 0.012 | -0.088 | -0.048 |
| | b.forest.loss.scaled | 0.001 | 0.004 | -0.005 | 0.006 |

| | | | | | |
|---|---|---|---|---|---|
| (N = 493 / 2,339) | b.duration.scaled | 0.027 | 0.004 | 0.021 | 0.033 |
| | sigma | 0.079 | 0.003 | 0.075 | 0.083 |
| | biome.boreal.forests.taiga | -0.012 | 0.016 | -0.041 | 0.010 |
| | biome.deserts.and.xeric.shrublands | 0.001 | 0.020 | -0.039 | 0.043 |
| | biome.mediterranean.forests.woodlands.and.scrub | 0.010 | 0.018 | -0.019 | 0.044 |
| | biome.montane.grasslands.and.shrublands | 0.006 | 0.020 | -0.027 | 0.047 |
| | biome.temperate.broadleaf.and.mixed.forests | -0.021 | 0.013 | -0.043 | 0 |
| | biome.temperate.conifer.forests | -0.004 | 0.012 | -0.027 | 0.016 |
| | biome.temperate.grasslands.savannas.and.shrublands | -0.002 | 0.019 | -0.037 | 0.033 |
| | biome.tropical.and.subtropical.grasslands.savannas.and.shrublands | 0.003 | 0.020 | -0.034 | 0.043 |
| | biome.tropical.and.subtropical.moist.broadleaf.forests | 0.020 | 0.018 | -0.008 | 0.048 |
| **Richness gains and forest cover loss (2000-2016)** (N = 377 / 2,339) | b.intercept | 0.095 | 0.015 | 0.065 | 0.121 |
| | b.forest.loss.scaled | 0.012 | 0.006 | 0.002 | 0.022 |
| | b.duration.scaled | -0.033 | 0.006 | -0.042 | -0.023 |
| | sigma | 0.114 | 0.004 | 0.108 | 0.122 |
| | biome.boreal.forests.taiga | 0.044 | 0.028 | 0 | 0.087 |
| | biome.deserts.and.xeric.shrublands | 0.004 | 0.027 | -0.045 | 0.063 |
| | biome.flooded.grasslands.and.savannas | -0.001 | 0.028 | -0.066 | 0.050 |
| | biome.mediterranean.forests.woodlands.and.scrub | -0.007 | 0.024 | -0.056 | 0.033 |
| | biome.montane.grasslands.and.shrublands | -0.011 | 0.028 | -0.074 | 0.034 |
| | biome.temperate.broadleaf.and.mixed.forests | 0.008 | 0.017 | -0.019 | 0.041 |
| | biome.temperate.conifer.forests | -0.006 | 0.016 | -0.035 | 0.023 |
| | biome.temperate.grasslands.savannas.and.shrublands | 0.003 | 0.023 | -0.039 | 0.048 |
| | biome.tropical.and.subtropical.grasslands.savannas.and.shrublands | -0.014 | 0.029 | -0.076 | 0.032 |
| | biome.tropical.and.subtropical.moist.broadleaf.forests | -0.012 | 0.025 | -0.065 | 0.028 |
| **Richness losses and forest cover gain (2000-2016)** (N = 73 / 2,339) | b.intercept | -0.041 | 0.021 | -0.074 | 0.003 |
| | b.forest.gain.scaled | 0.009 | 0.008 | -0.004 | 0.022 |
| | b.duration.scaled | 0.027 | 0.009 | 0.012 | 0.042 |
| | sigma | 0.068 | 0.006 | 0.059 | 0.078 |
| | biome.boreal.forests.taiga | 0.012 | 0.025 | -0.028 | 0.072 |
| | biome.mediterranean.forests.woodlands.and.scrub | 0.007 | 0.027 | -0.044 | 0.074 |
| | biome.montane.grasslands.and.shrublands | 0.004 | 0.027 | -0.053 | 0.066 |
| | biome.temperate.broadleaf.and.mixed.forests | -0.016 | 0.023 | -0.064 | 0.017 |
| | biome.temperate.conifer.forests | -0.025 | 0.028 | -0.076 | 0.010 |
| | biome.tropical.and.subtropical.moist.broadleaf.forests | 0.011 | 0.022 | -0.025 | 0.061 |
| **Richness gains and forest cover gain (2000-2016)** (N = 41 / 2,339) | b.intercept | 0.070 | 0.040 | -0.001 | 0.133 |
| | b.forest.gain.scaled | 0.012 | 0.028 | -0.035 | 0.056 |
| | b.duration.scaled | -0.075 | 0.026 | -0.115 | -0.031 |
| | sigma | 0.128 | 0.016 | 0.102 | 0.153 |
| | biome.boreal.forests.taiga | -0.014 | 0.042 | -0.103 | 0.063 |
| | biome.mediterranean.forests.woodlands.and.scrub | -0.004 | 0.047 | -0.108 | 0.093 |
| | biome.montane.grasslands.and.shrublands | -0.014 | 0.047 | -0.120 | 0.064 |
| | biome.temperate.broadleaf.and.mixed.forests | 0.056 | 0.051 | -0.011 | 0.138 |
| | biome.temperate.conifer.forests | 0.014 | 0.042 | -0.055 | 0.109 |
| | biome.tropical.and.subtropical.grasslands.savannas.and.shrublands | -0.009 | 0.049 | -0.124 | 0.081 |
| | biome.tropical.and.subtropical.moist.broadleaf.forests | -0.017 | 0.046 | -0.122 | 0.068 |

| | | | | | |
|---|---|---|---|---|---|
| **Turnover and overall forest loss** (N = 357 / 2,339) | b.intercept | -0.891 | 0.057 | -0.993 | -0.775 |
| | b.zoi.intercept | -1.652 | 0.138 | -1.877 | -1.434 |
| | b.coi.intercept | -1.533 | 0.296 | -1.998 | -1.044 |
| | b.forest.loss.scaled | -0.018 | 0.036 | -0.076 | 0.042 |
| | b.duration.scaled | -0.156 | 0.071 | -0.265 | -0.037 |
| | phi | 12.932 | 1.013 | 11.364 | 14.591 |
| | biome.boreal.forests.taiga | 0.011 | 0.057 | -0.122 | 0.225 |
| | biome.deserts.and.xeric.shrublands | -0.003 | 0.055 | -0.164 | 0.133 |
| | biome.mediterranean.forests.woodlands.and.scrub | -0.004 | 0.058 | -0.194 | 0.153 |
| | biome.montane.grasslands.and.shrublands | 0 | 0.058 | -0.201 | 0.183 |
| | biome.temperate.broadleaf.and.mixed.forests | -0.015 | 0.046 | -0.142 | 0.080 |
| | biome.temperate.conifer.forests | 0.010 | 0.049 | -0.092 | 0.145 |
| | biome.temperate.grasslands.savannas.and.shrublands | 0.011 | 0.051 | -0.091 | 0.160 |
| | biome.tropical.and.subtropical.dry.broadleaf.forests | 0.001 | 0.057 | -0.170 | 0.195 |
| | biome.tropical.and.subtropical.grasslands.savannas.and.shrublands | 0.002 | 0.058 | -0.159 | 0.202 |
| | biome.tropical.and.subtropical.moist.broadleaf.forests | -0.017 | 0.064 | -0.280 | 0.120 |
| **Turnover and forest cover loss (2000-2016)** (N = 870 / 2,339) | b.intercept | -0.524 | 0.104 | -0.728 | -0.373 |
| | b.zoi.intercept | -1.705 | 0.092 | -1.841 | -1.548 |
| | b.coi.intercept | -1.308 | 0.192 | -1.636 | -1.021 |
| | b.forest.loss.scaled | -0.018 | 0.026 | -0.061 | 0.023 |
| | b.duration.scaled | -0.274 | 0.030 | -0.321 | -0.227 |
| | phi | 7.298 | 0.361 | 6.747 | 7.886 |
| | biome.boreal.forests.taiga | 0.054 | 0.116 | -0.133 | 0.288 |
| | biome.deserts.and.xeric.shrublands | 0.045 | 0.168 | -0.228 | 0.498 |
| | biome.flooded.grasslands.and.savannas | -0.063 | 0.172 | -0.538 | 0.237 |
| | biome.mediterranean.forests.woodlands.and.scrub | 0.034 | 0.151 | -0.267 | 0.384 |
| | biome.montane.grasslands.and.shrublands | 0.001 | 0.171 | -0.419 | 0.414 |
| | biome.temperate.broadleaf.and.mixed.forests | 0.054 | 0.101 | -0.094 | 0.274 |
| | biome.temperate.conifer.forests | 0.135 | 0.122 | -0.025 | 0.353 |
| | biome.temperate.grasslands.savannas.and.shrublands | 0.037 | 0.133 | -0.191 | 0.330 |
| | biome.tropical.and.subtropical.grasslands.savannas.and.shrublands | -0.196 | 0.251 | -0.681 | 0.109 |
| | biome.tropical.and.subtropical.moist.broadleaf.forests | -0.118 | 0.154 | -0.422 | 0.082 |
| **Turnover and forest cover gain (2000-2016)** (N = 111 / 2,339) | b.intercept | -0.924 | 0.283 | -1.616 | -0.366 |
| | b.zoi.intercept | -1.382 | 0.216 | -1.720 | -1.033 |
| | b.coi.intercept | -0.552 | 0.348 | -1.126 | 0.012 |
| | b.forest.gain.scaled | -0.150 | 0.090 | -0.297 | -0.010 |
| | b.duration.scaled | -0.393 | 0.093 | -0.542 | -0.234 |
| | phi | 10.729 | 1.647 | 8.198 | 13.400 |
| | biome.boreal.forests.taiga | 0.411 | 0.431 | -0.172 | 1.206 |
| | biome.mediterranean.forests.woodlands.and.scrub | -0.002 | 0.499 | -1.299 | 1.151 |
| | biome.montane.grasslands.and.shrublands | 0 | 0.500 | -1.179 | 1.365 |
| | biome.temperate.broadleaf.and.mixed.forests | 0.067 | 0.286 | -0.512 | 0.746 |
| | biome.temperate.conifer.forests | 0.116 | 0.291 | -0.377 | 0.897 |
| | biome.tropical.and.subtropical.grasslands.savannas.and.shrublands | -0.637 | 0.658 | -1.816 | 0.148 |
| | biome.tropical.and.subtropical.moist.broadleaf.forests | -0.019 | 0.334 | -0.723 | 0.694 |
| | b.classmammals | 6.684 | 1.369 | 4.429 | 8.916 |

| | | | | | |
|---|---|---|---|---|---|
| **Population change lags across taxa** (N = 841 / 4,228) | b.classbirds | 9.136 | 1.087 | 7.193 | 10.944 |
| | b.classamphibians | 7.900 | 3.819 | 1.893 | 13.935 |
| | b.classreptiles | 2.357 | 2.679 | -1.893 | 6.608 |
| | sigma | 8.137 | 0.197 | 7.830 | 8.464 |
| | biome.boreal.forests.taiga | -0.225 | 1.174 | -2.260 | 1.695 |
| | biome.flooded.grasslands.and.savannas | -0.648 | 2.476 | -4.770 | 3.849 |
| | biome.mediterranean.forests.woodlands.and.scrub | -1.707 | 2.168 | -5.133 | 1.891 |
| | biome.temperate.broadleaf.and.mixed.forests | -2.251 | 1.157 | -4.068 | -0.184 |
| | biome.temperate.coniferous.forests | -0.843 | 1.350 | -3.148 | 1.348 |
| | biome.temperate.grasslands.savannas.and.shrublands | 5.008 | 1.398 | 2.702 | 7.240 |
| | biome.tropical.and.subtropical.coniferous.forests | 0.033 | 2.682 | -4.539 | 4.665 |
| | biome.tropical.and.subtropical.dry.broadleaf.forests | -2.146 | 2.534 | -6.585 | 2.077 |
| | biome.tropical.and.subtropical.grasslands.savannas.and.shrublands | 2.528 | 1.746 | -0.297 | 5.370 |
| | biome.tropical.and.subtropical.moist.broadleaf.forests | 1.796 | 1.987 | -1.342 | 5.050 |
| | biome.tundra | -1.318 | 1.535 | -3.772 | 1.270 |
| **Richness change lags across taxa** (N = 728 / 2,339) | b.taxa.amphibians | 9.691 | 2.065 | 6.142 | 13.057 |
| | b.taxa.invertebrates | -1.536 | 2.430 | -5.687 | 2.300 |
| | b.taxa.birds | 8.370 | 2.029 | 4.812 | 11.588 |
| | b.taxa.mammals | 6.937 | 2.407 | 3.041 | 11.038 |
| | b.taxa.otherplants | 9.207 | 2.647 | 5.159 | 13.557 |
| | b.taxa.trees | 17.389 | 2.526 | 13.379 | 21.693 |
| | sigma | 4.585 | 0.122 | 4.387 | 4.779 |
| | biome.map.boreal.forests.taiga | 2.581 | 2.879 | -2.134 | 7.321 |
| | biome.map.deserts.and.xeric.shrublands | -0.649 | 2.101 | -3.914 | 2.991 |
| | biome.map.multiple.ecoregions | -4.292 | 2.047 | -7.532 | -0.694 |
| | biome.map.temperate.broadleaf.and.mixed.forests | 6.755 | 2.257 | 3.076 | 10.476 |
| | biome.map.temperate.coniferous.forest | -6.899 | 2.614 | -11.178 | -2.646 |
| | biome.map.temperate.grasslands..savannas.and.shrublands | 2.136 | 2.032 | -1.121 | 5.758 |
| | biome.map.tropical.and.subtropical.dry.broadleaf.forests | -2.233 | 3.527 | -7.837 | 3.680 |
| | biome.map.tropical.and.subtropical.grasslands..savannas.and.shrublands | 4.577 | 2.564 | 0.213 | 8.611 |
| | biome.map.tropical.and.subtropical.moist.broadleaf.forests | -1.601 | 3.243 | -7.023 | 3.518 |
| **Turnover lags across taxa** (N = 2,157 / 2,339) | b.taxa.trees | 18.493 | 2.876 | 13.600 | 23.276 |
| | b.taxa.otherplants | 12.571 | 2.724 | 7.872 | 17.129 |
| | b.taxa.mammals | 11.017 | 2.708 | 6.457 | 15.569 |
| | b.taxa.birds | 8.341 | 2.476 | 4.159 | 12.557 |
| | b.taxa.amphibians | 10.042 | 2.500 | 6.098 | 14.552 |
| | b.taxa.invertebrates | 11.500 | 2.584 | 7.214 | 15.858 |
| | sigma | 4.662 | 0.069 | 4.551 | 4.773 |
| | biome.map.boreal.forests.taiga | -0.647 | 3.328 | -6.219 | 4.794 |
| | biome.map.deserts.and.xeric.shrublands | -3.153 | 2.494 | -7.478 | 0.989 |
| | biome.map.multiple.ecoregions | -5.443 | 2.465 | -9.816 | -1.399 |
| | biome.map.temperate.broadleaf.and.mixed.forests | -4.990 | 2.504 | -9.266 | -0.785 |
| | biome.map.temperate.coniferous.forest | -9.358 | 2.703 | -13.802 | -4.656 |
| | biome.map.temperate.grasslands..savannas.and.shrublands | 3.057 | 2.485 | -1.238 | 7.187 |

| | | | | | |
|---|---|---|---|---|---|
| | biome.map.tropical.and.subtropical.dry.broadleaf.forests | 4.616 | 4.323 | -2.065 | 12.130 |
| | biome.map.tropical.and.subtropical.grasslands..savannas.and.shrublands | 13.696 | 3.059 | 8.664 | 18.792 |
| | biome.map.tropical.and.subtropical.moist.broadleaf.forests | -6.432 | 2.801 | -11.174 | -1.780 |
| | biome.map.tundra | 8.372 | 2.566 | 3.894 | 12.843 |
| **Mammal generation time and population change temporal lags** (N = 83 / 117) | b.intercept | 4.047 | 1.228 | 2.131 | 6.003 |
| | b.generation.time | 0.336 | 0.137 | 0.110 | 0.551 |
| | sigma | 6.448 | 0.494 | 5.687 | 7.251 |
| **Bird generation time and population change temporal lags** (N = 545 / 599) | b_Intercept | 6.976 | 1.151 | 4.683 | 9.228 |
| | b_mean_gentime | 0.842 | 0.223 | 0.411 | 1.291 |
| | sigma | 9.158 | 0.279 | 8.644 | 9.723 |
| **Population declines, overall forest loss and species' geographic range** (N = 359 / 4,228) | b.intercept | -0.060 | 0.012 | -0.087 | -0.042 |
| | b.forest.loss.scaled | -0.001 | 0.001 | -0.003 | 0.002 |
| | b.range.scaled | -0.004 | 0.001 | -0.006 | -0.001 |
| | b.duration.scaled | 0.007 | 0.002 | 0.005 | 0.010 |
| | b.forest.loss.scaled.range.scaled | 0 | 0.002 | -0.003 | 0.002 |
| | sigma | 0.024 | 0.001 | 0.023 | 0.025 |
| | biome.boreal.forests.taiga | 0.011 | 0.013 | -0.008 | 0.038 |
| | biome.temperate.broadleaf.and.mixed.forests | 0.007 | 0.012 | -0.013 | 0.033 |
| | biome.temperate.coniferous.forests | 0.003 | 0.012 | -0.017 | 0.029 |
| | biome.temperate.grasslands.savannas.and.shrublands | 0.004 | 0.013 | -0.018 | 0.031 |
| | biome.tropical.and.subtropical.coniferous.forests | -0.011 | 0.019 | -0.045 | 0.022 |
| | biome.tropical.and.subtropical.grasslands.savannas.and.shrublands | -0.044 | 0.030 | -0.086 | 0.001 |
| | biome.tundra | 0.026 | 0.016 | 0 | 0.051 |
| **Population increases, overall forest loss and species' geographic range** (N = 302 / 4,228) | b.intercept | 0.050 | 0.008 | 0.037 | 0.076 |
| | b.forest.loss.scaled | 0.002 | 0.002 | -0.001 | 0.005 |
| | b.range.scaled | -0.001 | 0.002 | -0.004 | 0.002 |
| | b.duration.scaled | -0.010 | 0.002 | -0.014 | -0.007 |
| | b.forest.loss.scaled.range.scaled | 0.001 | 0.002 | -0.002 | 0.004 |
| | sigma | 0.029 | 0.001 | 0.027 | 0.031 |
| | biome.boreal.forests.taiga | -0.003 | 0.008 | -0.028 | 0.013 |
| | biome.mediterranean.forests.woodlands.and.scrub | 0.009 | 0.013 | -0.010 | 0.038 |
| | biome.temperate.broadleaf.and.mixed.forests | -0.005 | 0.008 | -0.032 | 0.008 |
| | biome.temperate.coniferous.forests | -0.003 | 0.008 | -0.030 | 0.011 |
| | biome.temperate.grasslands.savannas.and.shrublands | -0.004 | 0.009 | -0.031 | 0.011 |
| | biome.tropical.and.subtropical.grasslands.savannas.and.shrublands | -0.004 | 0.013 | -0.042 | 0.019 |
| | biome.tropical.and.subtropical.moist.broadleaf.forests | 0.035 | 0.046 | -0.005 | 0.101 |
| | biome.tundra | -0.009 | 0.013 | -0.039 | 0.006 |
| **Population declines, overall forest loss and species' mean population size** (N = 36 / 4,228) | b.intercept | 0.014 | 0.041 | -0.050 | 0.082 |
| | b.forest.loss.scaled | 0.020 | 0.027 | -0.022 | 0.066 |
| | b.meanpop.scaled | 0.031 | 0.044 | -0.044 | 0.099 |
| | b.duration.scaled | 0.031 | 0.008 | 0.019 | 0.044 |
| | b.forest.loss.scaled.meanpop.scaled | 0.035 | 0.049 | -0.045 | 0.115 |
| | sigma | 0.040 | 0.006 | 0.031 | 0.050 |
| | biome.boreal.forests.taiga | -0.005 | 0.024 | -0.055 | 0.036 |

| | | | | | |
|---|---|---|---|---|---|
| | biome.deserts.and.xeric.shrublands | -0.014 | 0.026 | -0.067 | 0.025 |
| | biome.mediterranean.forests.woodlands.and.scrub | -0.016 | 0.023 | -0.056 | 0.018 |
| | biome.temperate.broadleaf.and.mixed.forests | -0.015 | 0.018 | -0.046 | 0.012 |
| | biome.temperate.coniferous.forests | 0.031 | 0.030 | -0.009 | 0.079 |
| | biome.temperate.grasslands.savannas.and.shrublands | -0.010 | 0.020 | -0.050 | 0.021 |
| | biome.tropical.and.subtropical.grasslands.savannas.and.shrublands | -0.005 | 0.020 | -0.046 | 0.030 |
| | biome.tundra | 0.037 | 0.032 | -0.007 | 0.083 |
| **Population increases, overall forest loss and species' mean population size** (N = 43 / 4,228) | b.intercept | 0.038 | 0.022 | 0.002 | 0.077 |
| | b.forest.loss.scaled | 0.009 | 0.009 | -0.005 | 0.023 |
| | b.meanpop.scaled | 0.091 | 0.044 | 0.012 | 0.158 |
| | b.duration.scaled | -0.025 | 0.010 | -0.041 | -0.010 |
| | b.forest.loss.scaled.meanpop.scaled | 0.089 | 0.047 | 0.011 | 0.163 |
| | sigma | 0.049 | 0.006 | 0.040 | 0.059 |
| | biome.boreal.forests.taiga | 0 | 0.015 | -0.036 | 0.036 |
| | biome.mediterranean.forests.woodlands.and.scrub | 0.006 | 0.014 | -0.017 | 0.040 |
| | biome.temperate.broadleaf.and.mixed.forests | 0 | 0.011 | -0.024 | 0.024 |
| | biome.temperate.grasslands.savannas.and.shrublands | -0.009 | 0.017 | -0.055 | 0.016 |
| | biome.tropical.and.subtropical.grasslands.savannas.and.shrublands | 0.001 | 0.014 | -0.026 | 0.041 |
| | biome.tropical.and.subtropical.moist.broadleaf.forests | 0.006 | 0.015 | -0.019 | 0.050 |
| | biome.tundra | -0.007 | 0.014 | -0.045 | 0.013 |
| **Population declines, overall forest loss and species' habitat specificity** (N = 376 / 4,228) | b.intercept | -0.054 | 0.004 | -0.065 | -0.047 |
| | b.forest.loss.scaled | -0.001 | 0.002 | -0.003 | 0.002 |
| | b.habspec.scaled | -0.003 | 0.001 | -0.006 | -0.001 |
| | b.duration.scaled | 0.012 | 0.002 | 0.010 | 0.015 |
| | b.forest.loss.scaled.habspec.scaled | 0 | 0.002 | -0.002 | 0.003 |
| | sigma | 0.026 | 0.001 | 0.025 | 0.028 |
| | biome.boreal.forests.taiga | 0.002 | 0.004 | -0.006 | 0.013 |
| | biome.deserts.and.xeric.shrublands | -0.006 | 0.011 | -0.035 | 0.007 |
| | biome.mediterranean.forests.woodlands.and.scrub | -0.001 | 0.006 | -0.018 | 0.012 |
| | biome.temperate.broadleaf.and.mixed.forests | 0 | 0.004 | -0.007 | 0.011 |
| | biome.temperate.coniferous.forests | -0.001 | 0.005 | -0.011 | 0.009 |
| | biome.temperate.grasslands.savannas.and.shrublands | 0.001 | 0.005 | -0.009 | 0.014 |
| | biome.tropical.and.subtropical.coniferous.forests | -0.002 | 0.008 | -0.025 | 0.013 |
| | biome.tropical.and.subtropical.grasslands.savannas.and.shrublands | -0.006 | 0.009 | -0.028 | 0.006 |
| | biome.tropical.and.subtropical.moist.broadleaf.forests | 0.004 | 0.008 | -0.009 | 0.027 |
| | biome.tundra | 0.012 | 0.012 | -0.002 | 0.029 |
| **Population increases, overall forest loss and species' habitat specificity** (N = 316 / 4,228) | b.intercept | 0.047 | 0.003 | 0.041 | 0.054 |
| | b.forest.loss.scaled | 0.001 | 0.002 | -0.002 | 0.004 |
| | b.habspec.scaled | -0.003 | 0.002 | -0.007 | 0 |
| | b.duration.scaled | -0.011 | 0.002 | -0.014 | -0.008 |
| | b.forest.loss.scaled.habspec.scaled | 0.002 | 0.003 | -0.002 | 0.006 |
| | sigma | 0.029 | 0.001 | 0.027 | 0.031 |
| | biome.boreal.forests.taiga | 0 | 0.003 | -0.009 | 0.006 |
| | biome.mediterranean.forests.woodlands.and.scrub | 0.002 | 0.004 | -0.004 | 0.020 |
| | biome.temperate.broadleaf.and.mixed.forests | -0.001 | 0.003 | -0.009 | 0.004 |
| | biome.temperate.coniferous.forests | 0 | 0.003 | -0.010 | 0.006 |

| | | | | | |
|---|---|---|---|---|---|
| | biome.temperate.grasslands.savannas.and.shrublands | -0.001 | 0.003 | -0.011 | 0.006 |
| | biome.tropical.and.subtropical.grasslands.savannas.and.shrublands | -0.001 | 0.004 | -0.017 | 0.006 |
| | biome.tropical.and.subtropical.moist.broadleaf.forests | 0.002 | 0.005 | -0.007 | 0.024 |
| | biome.tundra | 0 | 0.003 | -0.012 | 0.007 |
| **Population declines, overall forest loss in the tropics and elsewhere** (N = 470 / 4,228) | b.intercept | -0.050 | 0.003 | -0.057 | -0.044 |
| | b.forest.loss.scaled | -0.002 | 0.001 | -0.004 | 0 |
| | b.tropicaltrue | -0.074 | 0.034 | -0.127 | -0.018 |
| | b.duration.scaled | 0.008 | 0.002 | 0.005 | 0.010 |
| | b.forest.loss.scaled.tropicaltrue | -0.032 | 0.029 | -0.078 | 0.014 |
| | sigma | 0.026 | 0.001 | 0.024 | 0.027 |
| | biome.boreal.forests.taiga | 0.001 | 0.003 | -0.005 | 0.009 |
| | biome.deserts.and.xeric.shrublands | -0.002 | 0.006 | -0.022 | 0.009 |
| | biome.mediterranean.forests.woodlands.and.scrub | -0.002 | 0.005 | -0.017 | 0.006 |
| | biome.temperate.broadleaf.and.mixed.forests | -0.001 | 0.003 | -0.008 | 0.005 |
| | biome.temperate.coniferous.forests | 0 | 0.003 | -0.008 | 0.007 |
| | biome.temperate.grasslands.savannas.and.shrublands | 0.001 | 0.004 | -0.006 | 0.013 |
| | biome.tropical.and.subtropical.coniferous.forests | -0.001 | 0.005 | -0.018 | 0.009 |
| | biome.tropical.and.subtropical.grasslands.savannas.and.shrublands | -0.001 | 0.004 | -0.014 | 0.009 |
| | biome.tropical.and.subtropical.moist.broadleaf.forests | 0.003 | 0.006 | -0.007 | 0.023 |
| | biome.tundra | 0.004 | 0.006 | -0.003 | 0.018 |
| **Population increases, overall forest loss in the tropics and elsewhere** (N = 369 / 4,228) | b.intercept | 0.049 | 0.004 | 0.044 | 0.061 |
| | b.forest.loss.scaled | 0.001 | 0.002 | -0.002 | 0.003 |
| | b.tropicaltrue | -0.025 | 0.010 | -0.041 | -0.009 |
| | b.duration.scaled | -0.015 | 0.002 | -0.018 | -0.012 |
| | b.forest.loss.scaled.tropicaltrue | -0.009 | 0.005 | -0.018 | -0.001 |
| | sigma | 0.029 | 0.001 | 0.027 | 0.031 |
| | biome.boreal.forests.taiga | 0 | 0.003 | -0.012 | 0.006 |
| | biome.mediterranean.forests.woodlands.and.scrub | 0.002 | 0.005 | -0.005 | 0.020 |
| | biome.temperate.broadleaf.and.mixed.forests | -0.002 | 0.004 | -0.015 | 0.004 |
| | biome.temperate.coniferous.forests | -0.001 | 0.004 | -0.013 | 0.006 |
| | biome.temperate.grasslands.savannas.and.shrublands | -0.001 | 0.004 | -0.014 | 0.006 |
| | biome.tropical.and.subtropical.grasslands.savannas.and.shrublands | 0 | 0.004 | -0.012 | 0.015 |
| | biome.tropical.and.subtropical.moist.broadleaf.forests | 0.004 | 0.007 | -0.006 | 0.039 |
| | biome.tundra | -0.002 | 0.005 | -0.019 | 0.004 |
| **Population declines, overall forest loss and starting tree cover** (N = 481 / 2,339) | b.intercept | -0.050 | 0.003 | -0.055 | -0.045 |
| | b.forest.loss.scaled | 0.003 | 0.004 | -0.002 | 0.009 |
| | b.sum.area.scaled | -0.007 | 0.003 | -0.012 | -0.002 |
| | b.duration.scaled | 0.012 | 0.001 | 0.010 | 0.014 |
| | b.forest.loss.scaled.sum.area.scaled | -0.003 | 0.004 | -0.009 | 0.004 |
| | sigma | 0.026 | 0.001 | 0.025 | 0.028 |
| | biome.boreal.forests.taiga | -0.001 | 0.002 | -0.007 | 0.002 |
| | biome.mediterranean.forests.woodlands.and.scrub | 0 | 0.002 | -0.006 | 0.006 |
| | biome.temperate.broadleaf.and.mixed.forests | 0 | 0.002 | -0.003 | 0.005 |
| | biome.temperate.coniferous.forests | 0 | 0.002 | -0.004 | 0.006 |
| | biome.temperate.grasslands.savannas.and.shrublands | 0 | 0.002 | -0.006 | 0.006 |
| | biome.tropical.and.subtropical.dry.broadleaf.forests | 0 | 0.002 | -0.008 | 0.005 |

| | | | | | |
|---|---|---|---|---|---|
| | biome.tropical.and.subtropical.grasslands.savannas.and.shrublands | 0 | 0.002 | -0.005 | 0.010 |
| | biome.tropical.and.subtropical.moist.broadleaf.forests | 0 | 0.002 | -0.006 | 0.006 |
| **Richness losses, overall forest loss and starting tree cover** (n = 493 / 2,339) | b.intercept | -0.069 | 0.013 | -0.090 | -0.044 |
| | b.forest.loss.scaled | -0.004 | 0.005 | -0.011 | 0.003 |
| | b.sum.area.scaled | 0.007 | 0.004 | 0 | 0.013 |
| | b.duration.scaled | 0.025 | 0.004 | 0.019 | 0.031 |
| | b.forest.loss.scaled.sum.area.scaled | 0.011 | 0.006 | 0.002 | 0.021 |
| | sigma | 0.079 | 0.003 | 0.075 | 0.083 |
| | biome.boreal.forests.taiga | -0.016 | 0.018 | -0.047 | 0.011 |
| | biome.deserts.and.xeric.shrublands | 0.001 | 0.023 | -0.043 | 0.046 |
| | biome.mediterranean.forests.woodlands.and.scrub | 0.012 | 0.020 | -0.018 | 0.048 |
| | biome.montane.grasslands.and.shrublands | 0.009 | 0.022 | -0.032 | 0.047 |
| | biome.temperate.broadleaf.and.mixed.forests | -0.025 | 0.015 | -0.050 | -0.002 |
| | biome.temperate.conifer.forests | -0.005 | 0.014 | -0.029 | 0.019 |
| | biome.temperate.grasslands.savannas.and.shrublands | -0.001 | 0.021 | -0.040 | 0.034 |
| | biome.tropical.and.subtropical.grasslands.savannas.and.shrublands | 0.003 | 0.022 | -0.036 | 0.048 |
| | biome.tropical.and.subtropical.moist.broadleaf.forests | 0.022 | 0.018 | -0.007 | 0.053 |

**Table S2. Number of time series across woody biomes of the world.**

| Database | Biome | Number of time series |
|---|---|---|
| *Living Planet Database* | Boreal forests/taiga | 919 |
| | Deserts and xeric shrublands | 20 |
| | Flooded grasslands and savannas | 3 |
| | Mediterranean forests woodlands and scrub | 128 |
| | Montane grasslands and shrublands | 2 |
| | Temperate broadleaf and mixed forests | 853 |
| | Temperate coniferous forests | 301 |
| | Temperate grasslands savannas and shrublands | 241 |
| | Tropical and subtropical coniferous forests | 1 |
| | Tropical and subtropical dry broadleaf forests | 5 |
| | Tropical and subtropical grasslands savannas and shrublands | 70 |
| | Tropical and subtropical moist broadleaf forests | 58 |
| | Tundra | 128 |
| *BioTIME* | Boreal forests/taiga | 222 |
| | Deserts and xeric shrublands | 42 |
| | Flooded grasslands and savannas | 4 |
| | Mangroves | 3 |
| | Mediterranean forests woodlands and scrub | 40 |
| | Montane grasslands and shrublands | 57 |
| | Temperate broadleaf and mixed forests | 1881 |
| | Temperate conifer forests | 532 |
| | Temperate grasslands savannas and shrublands | 129 |
| | Tropical and subtropical dry broadleaf forests | 3 |
| | Tropical and subtropical grasslands savannas and shrublands | 18 |
| | Tropical and subtropical moist broadleaf forests | 54 |
| | Tundra | 11 |

**Table S3. BioTIME study references. See http://biotime.st-andrews.ac.uk for further information about the BioTIME database.**

| StudyID | References |
|---|---|
| 10 | (*1*) |
| 18 | (*2*) |
| 39 | (*3-6*) |
| 41 | (*7*) |
| 42 | (*8-14*) |
| 44 | (*15-17*) |
| 46 | (*18*) |
| 47 | (*19*) |
| 51 | (*20, 21*) |
| 52 | (*22, 23*) |
| 53 | (*24, 25*) |
| 54 | (*26*) |
| 56 | (*27*) |
| 58 | (*28, 29*) |
| 59 | (*30*) |
| 60 | (*31-36*) |
| 63 | (*37, 38*) |
| 67 | (*39*) |
| 70 | (*40*) |
| 194 | (*41*) |
| 195 | (*42*) |
| 201 | (*43*). |
| 202 | (*44*) |
| 214 | (*45*) |
| 215 | (*46*) |
| 216 | (*47*) |
| 217 | (*48*) |
| 218 | (*49*) |

| | |
|---|---|
| 219 | (*50*) |
| 220 | (*50*) |
| 221 | (*51*) |
| 224 | (*52, 53*) |
| 225 | (*54*) |
| 226 | (*55*) |
| 233 | (*56*) |
| 234 | (*57-63*) |
| 235 | (*64-67*) |
| 239 | (*68*) |
| 240 | (*69*) |
| 241 | (*35, 70*) |
| 242 | (*71*) |
| 243 | (*72, 73*) |
| 248 | (*74*) |
| 249 | (*75*) |
| 255 | (*76*) |
| 270 | (*77*) |
| 275 | (*78*). |
| 277 & 279 | (*15-17*) |
| 293 | (*79*) |
| 294 | (*80, 81*) |
| 298 | (*82*) |
| 299 | (*83*) |
| 300 | (*84*) |
| 301 | (*85, 86*) |
| 302 | (*87*) |
| 303 | (*88*) |
| 304 | (*89-91*) |
| 305 | (*92*) |
| 307 | (*93*) |
| 308 | (*94*) |

| | |
|---|---|
| 309 | (*95*) |
| 311 | (*96*) |
| 312 | (*97*) |
| 313 | (*98*) |
| 314 | (*98*) |
| 315 | (*99*) |
| 316 | (*100*) |
| 317 | (*101*) |
| 318 | (*102*) |
| 319 | (*103*) |
| 321 | (*104*). |
| 322 | (*105*) |
| 323 | (*106*) |
| 324-326 | (*107*) |
| 327 | (*108*) |
| 329 | (*109*) |
| 331 | (*110-113*) |
| 333 | (*114*) |
| 334 | (*115-117*) |
| 336 | (*30*) |
| 337 | (*118*) |
| 339 | (*119*) |
| 340 | (*120*) |
| 341 | (*121*) |
| 342 | (*122*) |
| 343 | (*123*) |
| 344 | (*124*) |
| 345 | (*125*) |
| 346 | (*126*) |
| 348 | (*127*) |
| 352 | (*128*) |
| 353 | (*129*) |

| | |
|---|---|
| 355 | (*130*) |
| 356 | (*131*) |
| 357 | (*132*) |
| 358 | (*133*) |
| 360 | (*134-140*) |
| 361 | (*141*) |
| 362 | (*142, 143*) |
| 363 | (*144*) |
| 366 | (*145*) |
| 369 | (*146*) |
| 372 | (*147*). |
| 373 | (*148*) |
| 375 | (*149*) |
| 376 | (*150*) |
| 377 | (*149*) |
| 380 | (*151, 152*) |
| 381 | (*153*) |
| 382 | (*152, 154*) |
| 383-401 | (*155*) |
| 404 | (*156*) |
| 405 | (*157*) |
| 406 | (*158*) |
| 413 | (*159*) |
| 414 | (*159*) |
| 415 | (*159*) |
| 416 | (*159*) |
| 420 | (*160*) |
| 421 | (*161*) |
| 422-423 | (*162*) |
| 424 | (*163*) |
| 439-441 | (*164*) |
| 442-443 | (*165*) |

| | |
|---|---|
| 444 | (*166*) |
| 445 | (*167*) |
| 446 | (*168*) |
| 447 | (*169*) |
| 448 | (*170*) |
| 449 | (*171*) |
| 458-465 | (*172-174*) |
| 471 | (*175*) |
| 473 | (*176*) |
| 475 | (*177*) |
| 476 | (*178*) |
| 479 | (*179-184*) |
| 480-483 | (*181-184*) |
| 484 | (*181-185*) |
| 485-498 | (*181-184*) |
| 502 | (*186*) |
| 508-509 | (*181-184*) |
| 510 | (*187*) |
| 512 | (*188, 189*) |
| 515 | (*190-193*) |
| 516 | (*194-198*) |
| 518 | (*199, 200*) |
| 521 | (*201*) |
| 522 | (*202*) |
| 523 | (*203*) |
| 524 | (*204*) |

692

### References (including Supplementary Materials references)


1. IPBES, Summary for policymakers of the global assessment report on biodiversity and ecosystem services of the Intergovernmental Science-Policy Platform on Biodiversity and Ecosystem Services (2019).

11. T. Newbold, L. N. Hudson, S. L. L. Hill, S. Contu, I. Lysenko, R. A. Senior, L. Börger, D. J. Bennett, A. Choimes, B. Collen, J. Day, A. De Palma, S. Díaz, S. Echeverria-Londoño, M. J. Edgar, A. Feldman, M. Garon, M. L. K. Harrison, T. Alhusseini, D. J. Ingram, Y. Itescu, J. Kattge, V. Kemp, L. Kirkpatrick, M. Kleyer, D. L. P. Correia, C. D. Martin, S. Meiri, M. Novosolov, Y. Pan, H. R. P. Phillips, D. W. Purves, A. Robinson, J. Simpson, S. L. Tuck, E. Weiher, H. J. White, R. M. Ewers, G. M. Mace, J. P. W. Scharlemann, A. Purvis, Global effects of land use on local terrestrial biodiversity. *Nature*. **520**, 45–50 (2015).

24. S. Channan, K. Collins, W. R. Emanuel, Global mosaics of the standard MODIS land cover type data. University of Maryland and the Pacific Northwest National Laboratory, College Park, Maryland, USA. (2014).

25. LPI, Living Planet Index database. (2016) (available at www.livingplanetindex.org).

26. M. Dornelas, L. H. Antão, F. Moyes, A. E. Bates, A. E. Magurran, D. Adam, A. A. Akhmetzhanova, W. Appeltans, J. M. Arcos, H. Arnold, N. Ayyappan, G. Badihi, A. H. Baird, M. Barbosa, T. E. Barreto, C. Bässler, A. Bellgrove, J. Belmaker, L. Benedetti-Cecchi, B. J. Bett, A. D. Bjorkman, M. Błażewicz, S. A. Blowes, C. P. Bloch, T. C. Bonebrake, S. Boyd, M. Bradford, A. J. Brooks, J. H. Brown, H. Bruelheide, P. Budy, F. Carvalho, E. Castañeda-Moya, C. A. Chen, J. F. Chamblee, T. J. Chase, L. Siegwart Collier, S. K. Collinge, R. Condit, E. J. Cooper, J. H. C. Cornelissen, U. Cotano, S. Kyle Crow, G. Damasceno, C. H. Davies, R. A. Davis, F. P. Day, S. Degraer, T. S. Doherty, T. E. Dunn, G. Durigan, J. E. Duffy, D. Edelist, G. J. Edgar, R. Elahi, S. C. Elmendorf, A. Enemar, S. K. M. Ernest, R. Escribano, M. Estiarte, B. S. Evans, T.-Y. Fan, F. Turini Farah, L. Loureiro Fernandes, F. Z. Farneda, A. Fidelis, R. Fitt, A. M. Fosaa, G. A. Daher Correa Franco, G. E. Frank, W. R. Fraser, H. García, R. Cazzolla Gatti, O. Givan, E. Gorgone-Barbosa, W. A. Gould, C. Gries, G. D. Grossman, J. R. Gutierréz, S. Hale, M. E. Harmon, J. Harte, G. Haskins, D. L. Henshaw, L. Hermanutz, P. Hidalgo, P. Higuchi, A. Hoey, G. Van Hoey, A. Hofgaard, K. Holeck, R. D. Hollister, R. Holmes, M. Hoogenboom, C. Hsieh, S. P. Hubbell, F. Huettmann, C. L. Huffard, A. H. Hurlbert, N. Macedo Ivanauskas, D. Janík, U. Jandt, A. Jażdżewska, T. Johannessen, J. Johnstone, J. Jones, F. A. M. Jones, J. Kang, T. Kartawijaya, E. C. Keeley, D. A. Kelt, R. Kinnear, K. Klanderud, H. Knutsen, C. C. Koenig, A. R. Kortz, K. Král, L. A. Kuhnz, C.-Y. Kuo, D. J. Kushner, C. Laguionie-Marchais, L. T. Lancaster, C.

Min Lee, J. S. Lefcheck, E. Lévesque, D. Lightfoot, F. Lloret, J. D. Lloyd, A. López-Baucells, M. Louzao, J. S. Madin, B. Magnússon, S. Malamud, I. Matthews, K. P. McFarland, B. McGill, D. McKnight, W. O. McLarney, J. Meador, P. L. Meserve, D. J. Metcalfe, C. F. J. Meyer, A. Michelsen, N. Milchakova, T. Moens, E. Moland, J. Moore, C. Mathias Moreira, J. Müller, G. Murphy, I. H. Myers-Smith, R. W. Myster, A. Naumov, F. Neat, J. A. Nelson, M. Paul Nelson, S. F. Newton, N. Norden, J. C. Oliver, E. M. Olsen, V. G. Onipchenko, K. Pabis, R. J. Pabst, A. Paquette, S. Pardede, D. M. Paterson, R. Pélissier, J. Peñuelas, A. Pérez-Matus, O. Pizarro, F. Pomati, E. Post, H. H. T. Prins, J. C. Priscu, P. Provoost, K. L. Prudic, E. Pulliainen, B. R. Ramesh, O. Mendivil Ramos, A. Rassweiler, J. E. Rebelo, D. C. Reed, P. B. Reich, S. M. Remillard, A. J. Richardson, J. P. Richardson, I. van Rijn, R. Rocha, V. H. Rivera-Monroy, C. Rixen, K. P. Robinson, R. Ribeiro Rodrigues, D. de Cerqueira Rossa-Feres, L. Rudstam, H. Ruhl, C. S. Ruz, E. M. Sampaio, N. Rybicki, A. Rypel, S. Sal, B. Salgado, F. A. M. Santos, A. P. Savassi-Coutinho, S. Scanga, J. Schmidt, R. Schooley, F. Setiawan, K.-T. Shao, G. R. Shaver, S. Sherman, T. W. Sherry, J. Siciński, C. Sievers, A. C. da Silva, F. Rodrigues da Silva, F. L. Silveira, J. Slingsby, T. Smart, S. J. Snell, N. A. Soudzilovskaia, G. B. G. Souza, F. Maluf Souza, V. Castro Souza, C. D. Stallings, R. Stanforth, E. H. Stanley, J. Mauro Sterza, M. Stevens, R. Stuart-Smith, Y. Rondon Suarez, S. Supp, J. Yoshio Tamashiro, S. Tarigan, G. P. Thiede, S. Thorn, A. Tolvanen, M. Teresa Zugliani Toniato, Ø. Totland, R. R. Twilley, G. Vaitkus, N. Valdivia, M. I. Vallejo, T. J. Valone, C. Van Colen, J. Vanaverbeke, F. Venturoli, H. M. Verheye, M. Vianna, R. P. Vieira, T. Vrška, C. Quang Vu, L. Van Vu, R. B. Waide, C. Waldock, D. Watts, S. Webb, T. Wesołowski, E. P. White, C. E. Widdicombe, D. Wilgers, R. Williams, S. B. Williams, M. Williamson, M. R. Willig, T. J. Willis, S. Wipf, K. D. Woods, E. J.

Woehler, K. Zawada, M. L. Zettler, BioTIME: A database of biodiversity time series for the Anthropocene. *Global Ecology and Biogeography*. **27**, 760–786 (2018).

33. S. A. Blowes, S. R. Supp, L. H. Antão, A. Bates, H. Bruelheide, J. M. Chase, F. Moyes, A. Magurran, B. McGill, I. H. Myers-Smith, M. Winter, A. D. Bjorkman, D. E. Bowler, J. E. K.

Byrnes, A. Gonzalez, J. Hines, F. Isbell, H. P. Jones, L. M. Navarro, P. L. Thompson, M. Vellend, C. Waldock, M. Dornelas, The geography of biodiversity change in marine and terrestrial assemblages. *Science*. **366**, 339–345 (2019).

47. J. W. Veldman, J. C. Aleman, S. T. Alvarado, T. M. Anderson, S. Archibald, W. J. Bond, T. W. Boutton, N. Buchmann, E. Buisson, J. G. Canadell, M. de S. Dechoum, M. H. Diaz-Toribio, G. Durigan, J. J. Ewel, G. W. Fernandes, A. Fidelis, F. Fleischman, S. P. Good, D. M. Griffith, J.-M. Hermann, W. A. Hoffmann, S. Le Stradic, C. E. R. Lehmann, G. Mahy, A. N. Nerlekar, J. B. Nippert, R. F. Noss, C. P. Osborne, G. E. Overbeck, C. L. Parr, J. G.

Pausas, R. T. Pennington, M. P. Perring, F. E. Putz, J. Ratnam, M. Sankaran, I. B. Schmidt, C. B. Schmitt, F. A. O. Silveira, A. C. Staver, N. Stevens, C. J. Still, C. A. E. Strömberg, V. M. Temperton, J. M. Varner, N. P. Zaloumis, Comment on “The global tree restoration potential.” *Science*. **366**, eaay7976 (2019).
